# Genetic and phylogenetic features of the repetitive ITS copies in the *Hirsutella sinensis* genome and the genomically independent AT-biased *Ophiocordyceps sinensis* genotypes

**DOI:** 10.1101/2022.12.09.519819

**Authors:** Xiu-Zhang Li, Yu-Ling Li, Ya-Nan Wang, Jia-Shi Zhu

**Affiliations:** State Key Laboratory of Plateau Ecology and Agriculture, Qinghai Academy of Animal and Veterinary Sciences, Qinghai University, Xining, Qinghai 810016, China; State Key Laboratory of Breeding Base of Dao-di Herbs, National Resource Center for Chinese Materia Medica, China Academy of Chinese Medical Sciences, Beijing 100700, China; Institute of Biopharmaceutical and Health Engineering, Shenzhen International Graduate School, Tsinghua University, Shenzhen 518055, China

**Keywords:** Multiple genotypes of *Ophiocordyceps sinensis*, genomic independence, repeat-induced point mutations (RIP), insertion/deletion, transversion and transition point mutations, 5.8S gene transcript, transcriptional silencing, repetitive copies of authentic genes in multiple genomic loci

## Abstract

It has been hypothesized that AT-biased genotypes of *Ophiocordyceps sinensis* are generated through repeat-induced point mutation (RIP) and coexist as permanently nonfunctional internal transcribed spacer (ITS) pseudogenes in the genome of *Hirsutella sinensis* (GC-biased Genotype #1 of *O. sinensis*). This study examined the *H. sinensis* genome, which contains multiple repetitive ITS copies (GC content: 64.7±0.33%) with multiple insertion/deletion and transversion alleles, which were not generated through RIP mutagenesis that theoretically causes cytosine-to-thymine (C-to-T) and guanine-to-adenine (G-to-A) transitions. The repetitive ITS copies in the *H. sinensis* genome were found to be genetically and phylogenetically distinct from the AT-biased *O. sinensis* genotypes (GC content: 51.1±1.69%), which possess multiple transition alleles. The sequences of Genotypes #2–17, both GC- and AT-biased, are absent from the *H. sinensis* genome; these genotypes belong to interindividual *O. sinensis* fungi and differentially occur in different compartments of natural *Cordyceps sinensis,* with dynamic alterations in abundance occurring in an asynchronous, disproportional manner during *C. sinensis* maturation. Metatranscriptomic analyses of natural *C. sinensis* revealed the transcriptional silencing of 5.8S genes in all *C. sinensis*- colonizing fungi, including *H. sinensis*. The transcription assay reported by Li *et al*. [1] provided unsound, controversial evidence indicating that the 5.8S genes of AT-biased genotypes are nonfunctional pseudogenes. In addition to the single ITS locus analysis, repetitive genomic copies were also examined at multiple loci in the *H. sinensis* genome, and approximately 8.2% of the authentic genes had repetitive copies, including various transitions, transversions, and insertions/deletions. The transcripts for the repetitive copies, regardless of the decreases, increases, or bidirectional changes in the AT content, were identified in the *H. sinensis* transcriptome. These results are inconsistent with those of RIP mutagenesis, which generates pseudogenic, nonfunctional, repetitive copies. In conclusion, AT-biased genotypes of *O. sinensis* might have evolved through evolutionary mechanisms from a common ancestor over the long course of evolution, in parallel with GC-biased Genotype #1 *H. sinensis*.

## I. Introduction

Natural *Cordyceps sinensis* is one of most valued therapeutic agents in traditional Chinese medicine and has a rich history of clinical use in health maintenance, disease amelioration, postdisease and postsurgery recovery, and antiaging therapy [2–4]. The Chinese Pharmacopeia defines natural *C. sinensis* as an insect-fungal complex that includes the remains of a Hepialidae moth larva [5–9] and >90 fungal species from >37 genera [10–12], with differential co-occurrence of some of 17 *Ophiocordyceps sinensis* genotypes in different compartments of natural *C. sinensis* [5–9,13–32], *i.e.*, the natural *C. sinensis* insect-fungal complex ≠ the entomopathogenic fungus *O. sinensis*. However, since the 1840s, the Latin name *C. sinensis* has been indiscriminately used for both the natural insect-fungal complex and the teleomorph/holomorph of the *C. sinensis* fungus [6–8,31,33–34]. The fungus was renamed *Ophiocordyceps sinensis* using *Hirsutella sinensis* Strain EFCC 7287 (GC-biased Genotype #1 of *O. sinensis*) as the nomenclature reference, while the indiscriminately used name of the natural and cultivated insect-fungal complexes has remained unchanged [8,31,35]. Since then, especially after the improper implementation of the “One Fungus = One Name” nomenclature rule in *O. sinensis* research [36–38], the name *O. sinensis* has been used to refer to multiple teleomorphic and anamorphic fungi that have distinct genomes and has been indiscriminately applied to the natural insect-fungal complex [5–8,31,33]. The confusion surrounding Latin name use continues and has even spread from the scientific community to the public media and mass markets, causing a significant decrease in the wholesale and retail prices of natural *C. sinensis*. Because a consensus regarding a Latin name for the natural insect-fungal product has not been reached by taxonomists and because multiple genotypes of *O. sinensis* with undetermined taxonomic positions are referred to by the same Latin name [6–9,31,33,39–40], we temporarily refer to the 17 genotypes of the fungus/fungi as *O. sinensis*, including Genotype #1 *H. sinensis*, and we continue customarily to refer to the natural and cultivated insect-fungal complexes as *C. sinensis*; however, this practice only partially ameliorates the academic confusion arising from the indiscriminate use of Latin names and will likely be replaced by the exclusive use of distinct Latin names in the future. Zhang *et al*. [39] suggested using the non-Latin term “Chinese cordyceps” for the insect-fungi complex, however, this proposal has not been generally accepted because governmental regulations worldwide require every natural medicinal product to have a unique, exclusive Latin name.

Wei *et al*. [19] hypothesized that the GC-biased Genotype #1 *H. sinensis* is the sole anamorph of the solely teleomorphic *O. sinensis*. However, numerous studies have identified 17 mutant genotypes of *O. sinensis* in natural *C. sinensis*, which are indiscriminately referred to by the same Latin name [5–9,14,17–18,20–34,42]. Stensrud *et al*. [18] analyzed 71 ITS sequences that were annotated as belonging to *C. sinensis* (renamed *O. sinensis* by Sung *et al*. [35]) in GenBank under taxid: 72228 and reported the existence of 3 phylogenetic clades of mutant *O. sinensis* sequences using a Bayesian phylogenetic algorithm: Group A (GC-biased Genotype #1) and Groups B‒C (AT-biased Genotypes #4‒5). The authors reported that the variation in the 5.8S gene sequences of *O. sinensis* in Groups A‒C “far exceeds what is normally observed in fungi … even at higher taxonomic levels (genera and family)” and that the genotypes “share a common ancestor” and originated from “accelerated nrDNA evolution”. Zhang *et al*. [41] conducted a GenBank data mining study involving 397 ITS sequences annotated as belonging to *C. sinensis* or *O. sinensis*, followed by phylogenetic analysis using a minimum evolutionary algorithm, and confirmed the existence of 3 phylogenetic clades of mutant *O. sinensis* sequences: Clade A (GC-biased genotypes) and Clade B‒C (AT-biased Genotypes #4‒5). In contrast to the homogenous GC-biased Group-A sequences containing only Genotype #1 reported by Stensrud *et al*. [18], the Clade A reported by Zhang *et al*. [41] was quite divergent, containing the GC-biased sequences of Genotype #1 *H. sinensis* and several other mutant sequences that were phylogenetically distant from GC-biased Genotype #1, making it questionable whether they truly belonged to Genotype #1 *H. sinensis*.

Kinjo & Zang [14] and Mao *et al*. [29] reported that *O. sinensis* genotypes share the same *H. sinensis*- like morphological and growth characteristics. Xiao *et al*. [21] concluded that the genetically variable genotypic sequences likely belonged to independent *O. sinensis* fungi based on the differential coexistence of multiple *O. sinensis* mutants in the stroma and caterpillar body of natural *C. sinensis* in different maturation stages. In the debate regarding “cryptic species” [18] or independent fungi of *O. sinensis* [21], Zhang *et al*. [41] disproved the former and supported the latter based on the results of Kimura 2-parameter (K2P) analysis. Wei *et al*. [19] reported the existence of the sole GC-biased Genotype #1 teleomorph of *O. sinensis* in natural *C. sinensis*. However, the authors reported the sole AT-biased Genotype #4 teleomorph of *O. sinensis* in cultivated *C. sinensis*, contradicting the anamorphic inoculants of GC-biased *H. sinensis* strains that were used in the industrial cultivation project.

Li *et al*. [43] reported the differential occurrence of the mating-type genes of the *MAT1-1* and *MAT1-2* idiomorphs in the *H. sinensis* genome and the differential transcription and silencing of the mating-type genes, which invalidated the “self-fertilization” hypothesis under homothallism and pseudohomothallism [44–45] and instead suggested that the self-sterility of *O. sinensis* requires mating partners to accomplish sexual reproduction *via* physiological heterothallism or hybridization. In addition to identifying the molecular heterogeneity of genotypic *O. sinensis* fungi and the requirement of sexual partners for *O. sinensis* reproduction, Figure 3 of [45] illustrates the multicellular heterokaryotic structure of *C. sinensis* hyphae and ascospores, which include multiple mononucleated, binucleated, trinucleated, and tetranucleated cells. Zhang & Zhang [46] suggested that the various heterokaryotic cells of *C. sinensis* hyphae and ascospores likely contain different genetic materials.

The molecular heterogeneity and self-sterility features of *O. sinensis* and the multicellular heterokaryotic structure of natural *C. sinensis* challenge the sole anamorph hypothesis for *H. sinensis* (GC-biased Genotype #1 of *O. sinensis*). Responding to this academic challenge, Li *et al*. [1] proposed the “ITS pseudogene” hypothesis for all AT-biased genotypes “in a single genome” of GC-biased Genotype #1 *H. sinensis* based on (1) the identification of heterogeneous ITS sequences of both GC-biased Genotype #1 and AT-biased Genotype #5 of *O. sinensis* in 8 out of 15 clones following 25 days of liquid incubation of *C. sinensis* mono-ascospores and (2) the identification of the 5.8S gene cDNA of Genotype #1 but not of Genotype #5 in a cDNA library of Strain 1220, one of 8 genetically heterogeneous, ascosporic clones. According to this “ITS pseudogene” hypothesis, the simultaneous identification of homogenous Genotype #1 in 7 other clones derived from the same mono-ascospores was apparently ignored. The authors [1] overgeneralized the “ITS pseudogene” hypothesis to all AT-biased genotypes (#4‒6 and #15‒17) of *O. sinensis* based on insufficient evidence while attempting to justify AT-biased genotypes as pseudogenes in the single genome of GC-biased *H. sinensis* under the sole anamorph hypothesis for *H. sinensis* [19]. Li *et al*. [47] further postulated that all AT-biased *O. sinensis* genotypes “could have emerged either before or after” a new GC-biased “ITS haplotype was generated” through “repeat-induced point mutations (RIP)” that cause C-to-T (cytosine-to-thymine) and G-to-A (guanine-to-adenine) transitions and cooccur as AT-biased, nonfunctional repetitive ITS copies in “a single genome” of GC-biased *H. sinensis*.

Because of the unavailability of pure cultures of AT-biased *O. sinensis* genotypes for taxonomic determination, which prevents genomic, transcriptomic, and multigene/multilocus analyses of hypothetical pseudogenes, the present study included analysis of relevant genomic and transcriptomic sequences of GC-biased Genotype #1 available in GenBank to examine the “ITS pseudogene” hypothesis and the extended hypothesis that AT-biased genotypes formed after a new GC-biased “ITS haplotype was generated” in “a single genome” of *H. sinensis* through “RIP” mutagenesis. We also extended the single ITS locus analysis to genomic multilocus analysis for the repetitive copies of 104 of 1271 authentic *H. sinensis* genes. We discuss the genetic and phylogenetic characteristics of multiple repetitive copies (including repetitive ITS copies) in the genome of Genotype #1 of *O. sinensis*, the genomic independence of AT-biased genotypes, and whether sufficient evidence exists to conclude that the 5.8S genes of AT-biased genotypes are permanently nonfunctional pseudogenes.

## II. Materials and Methods

### II-1. Reagents

Common laboratory reagents such as agarose, electrophoresis reagents, nylon membranes, and blocking solution were purchased from Beijing Bioland Technology Company. The endonucleases *Ava*I, *Dra*I, and *Eco*RI used were produced by New England BioLabs, United States. Mercuric chloride (0.1%) for surface sterilization of freshly collected *C. sinensis* specimens was a gift from the Institute of Microbiology, Chinese Academy of Sciences. The Universal DNA Purification Kit was obtained from TIANGEN BIOTECH (China). The DNeasy Plant Mini Kit was obtained from the Qiagen, Germany. The Gel Extraction Kit was obtained from Omega Bio-Tek (United States). The *Taq* PCR Reagent Kit and Vector NTI Advance 9 software were purchased from Invitrogen (United States). All primers used were synthesized by Invitrogen Beijing Lab. or Beijing Bomaide Technology Co. The DIG High Prime DNA Labeling and Detection Kit was purchased from Roche Diagnostics (Indiana, USA).

### II-2. Fungal species in natural *C. sinensis*

Information regarding intrinsic fungal species in natural *C. sinensis* was summarized by Jiang & Yao [48], and subsequent discoveries of additional species in natural *C. sinensis* were reviewed by Li *et al*. [5–8,31]. The sequences of 17 genotypes of *O. sinensis* have been obtained and uploaded to GenBank by many research groups worldwide since 2001 and analyzed genetically and phylogenetically [5–9,13–14,16–32,41–42.49].

### II-3. Genetic, genomic, transcriptomic and protein sequences from GenBank

Five genome assemblies, ANOV00000000, JAAVMX000000000, LKHE00000000, LWBQ00000000, and NGJJ00000000, for the *H. sinensis* Strains Co18, IOZ07, 1229, ZJB12195, and CC1406-203, respectively, are available in GenBank [44,50–53] and were used for the analysis of repetitive copies of authentic genes in the ITS locus and multiple loci and multiple GC- and AT-biased *O. sinensis* genotypes.

The GenBank database includes at least 668 *O. sinensis* ITS sequences under GenBank taxid: 72228, 440 of which belong to GC-biased Genotypes #1‒3 and #7‒14 and 228 of which belong to AT-biased Genotypes #4‒6 and #15‒17 [16,18,22,31–32,34,36,41].

The mRNA transcriptome assembly GCQL00000000 for the *H. sinensis* Strain L0106 was uploaded to GenBank by Liu *et al*. [54]. Two metatranscriptome studies of natural *C. sinensis* have been reported. Xiang *et al*. [55] uploaded the GAGW00000000 metatranscriptome assembly to GenBank; this assembly was obtained from natural specimens (unknown maturation stages) collected from Kangding County in Sichuan Province, China. Xia *et al*. [56] uploaded the unassembled SRR5428527 Shotgun Read Archive (SRA) to GenBank for mature *C. sinensis* specimens collected from Deqin County in Yunnan Province, China; however, the assembled metatranscriptome sequences were uploaded to www.plantkingdomgdb.com/Ophiocordyceps_sinensis/data/cds/Ophiocordyceps_sinensis_CDS.fas, which is currently inaccessible, but a previously downloaded file was used for analysis.

GenBank does not provide annotations for authentic genes in the genome assemblies ANOV00000000, JAAVMX000000000, LKHE00000000, LWBQ00000000, and NGJJ00000000 of the *H. sinensis* Strains Co18, IOZ07, 1229, ZJB12195, and CC1406-203, respectively. One (JAACLJ010000002) of the 13 genome contigs for *Ophiocordyceps camponoti-floridani* Strain EC05 contained 1271 authentic genes [57], which was used to cross-reference, position, and annotate the authentic *H. sinensis* genes and their repetitive copies in the *H. sinensis* genome.

The transcription of the authentic *H. sinensis* genes and their repetitive genomic copies, as well as the translated protein sequences, were explored against the mRNA transcriptome assembly GCQL00000000 for the *H. sinensis* Strain L0106 and the metatranscriptome assemblies for natural *C. sinensis* using the GenBank BLAST algorithm (https://blast.ncbi.nlm.nih.gov/blast/).

### II-4. Genetic and phylogenetic analyses

The ITS sequences of multiple genotypes of *O. sinensis* and multiple repetitive genomic ITS copies as well as metatranscriptomic/transcriptomic sequences were analyzed using the GenBank BLAST algorithms (https://blast.ncbi.nlm.nih.gov/blast/). The DNA sequences and transcripts of authentic *H. sinensis* genes and their repetitive copies in the *H. sinensis* genome identified during multilocus analysis were also analyzed using BLAST algorithms.

A Bayesian phylogenetic tree of the ITS sequences and their repetitive copies was inferred using MrBayes v3.2.7 software (Markov chain Monte Carlo [MCMC] algorithm) based on 3x10^4^ samples, with a sampling frequency of 10^3^ iterations after discarding the first 25% of the samples from a total of 4x10^6^ iterations [8–9,58]. This phylogenetic analysis was performed at Nanjing Genepioneer Biotechnologies Co.

### II-5. Collection and on-site processing of fresh *C. sinensis* primers

Fresh *C. sinensis* specimens were purchased from a local market in Kangding County in Sichuan Province, China (altitude 3800‒4600 m; latitude 30°049’N, longitude 101°959’E) [42,49]. Immature *C. sinensis* specimens were characterized by the presence of a plump caterpillar body and very short stroma (1.0‒2.0 cm). Maturing *C. sinensis* specimens were characterized by the presence of a plump caterpillar body and a stroma that was 2.5‒5.0 cm in length, without the formation of an expanded fertile portion close to the stromal tip. Mature *C. sinensis* specimens were characterized by the presence of a plump or slightly less plump caterpillar body and long stroma (>5.0 cm) and by the formation of an expanded fertile portion close to the stromal tip, which was densely covered with numerous ascocarps [42–43]. Governmental permission was not required for *C. sinensis* purchases in local markets, and the collection of *C. sinensis* specimens from sales by local farmers fall under governmental regulations for traditional Chinese herbal products.

Freshly collected *C. sinensis* specimens were washed thoroughly in running water with gentle brushing and surface sterilized in 0.1% mercuric chloride for 10 min, followed by washing 3 times with sterile water [9,42]. The thoroughly cleaned and sterilized *C. sinensis* specimens were immediately frozen in liquid nitrogen and kept frozen during transport and storage prior to further processing.

### II-6. Extraction of genomic DNA and preparation of *C. sinensis* DNA

Caterpillar bodies and stromata of sterilized, frozen *C. sinensis* were separately ground into powder in liquid nitrogen. Genomic DNA was individually extracted from the powder samples using a DNeasy Plant Mini Kit [23,42].

### II-7. PCR amplification and amplicon sequencing

The genomic DNA of immature, maturing and mature stromata of natural *C. sinensis* was used as the template for PCR amplification. *H. sinensis*-specific primers were designed (forward primer, *Hsprp1*: 5’ ATTATCGAGTCACCACTCCCAAACCCCC 3’; reverse primer, *Hsprp3*: 5’ AGGTTCTCAGCGAGCTA 3’) based on alignment analysis of the ITS sequences (AB067721, AB067740, and AB067744, representing Genotypes #1 and #4‒5 of *O. sinensis*, respectively).

A touch-down PCR protocol was used, in which the annealing temperature was initially set at 70°C and subsequently decreased stepwise by 0.3°C every cycle for a total of 36 cycles [23,42]. The PCR amplicons were subjected to agarose gel electrophoresis, and the DNA bands were recovered, purified using a gel extraction kit, and sequenced either directly or after cloning the amplicons.

### II-8. Preparation of the fungus specific ITS1 probe and 18S internal control probe

An *H. sinensis*-specific probe (158 bp) was designed based on the ITS1 sequence of GC-biased Genotype #1 *H. sinensis* and generated through PCR using the forward primer *Hsprp1* described above and the reverse primer *Hsprp2* (5’ ATTTGCTTGCTTGACTGAGAGATGCC 3’) [42].

A nonspecific 18S internal control probe was designed based on alignments of GenBank 18S rDNA sequences (DQ838795, AB187268, AB067735, AB067736, and AB080090) of several *C. sinensis*- associated fungi. A region highly homologous to the 18S rDNA region, located more than 700 bp upstream of the ITS1 region of *C. sinensis*-associated fungi, was selected. The 18S control probe (401 bp) was generated through PCR using the forward primer *Icprp1* (5’ ACCGTCGTAGTCTTAACCAT 3’) and the reverse primer *Icprp2* (5’ ATCGGCTTGAGCCGATAGTC 3’) [42].

Both amplicons were confirmed by sequencing, and the probes were labeled using the DIG High Prime DNA Labeling and Detection Kit [42].

### II-9. Southern blotting analysis

The stromata and caterpillar bodies of immature and mature *C. sinensis* were subjected to Southern blotting examination. Genomic DNA was extracted from the samples and prepared by endonuclease digestion using *Ava*I, *Dra*I, and *Eco*RI [42]. The digested genomic DNA (5 µg of each sample) was subjected to agarose gel electrophoresis and transferred onto a nylon membrane through capillary transfer, after which the membrane was incubated in an oven at 58°C for 4 hours for fixation, followed by blocking of the binding sites on the membrane in blocking solution and hybridization with the labeled *H. sinensis*- specific probe at 40°C overnight. The hybridized transblotted membrane was viewed by colorimetric photography using a gel imaging system (Bio-Rad) and then stripped and reprobed with the labeled 18S internal control probe.

## III. Results

### III-1. Multiple GC-biased repetitive ITS copies in the *H. sinensis* genome

In general, fungal genomes contain multiple repetitive copies of the nrDNA ITS1-5.8S-ITS2 sequences. Li *et al*. [47] submitted their paper to IMA Fungus in January 2020, when 3 genome assemblies, namely, ANOV00000000, LKHE00000000, and LWBQ00000000, of the *H. sinensis* Strains Co18, 1229, and ZJB12195, respectively [44,51–52], were available in GenBank. However, only single copies of the ITS1-5.8S-ITS2 sequences could be identified in each of these genomes (the segments within ANOV01021709, LKHE01000582, and LWBQ01000008), probably because the repetitive ITS copies were discarded during the assembly of the genome shotgun reads, which indicated that Li *et al*. [1,47] might not have been able to obtain real and non-abridged genomic data for repetitive ITS copies before manuscript submission as the basis for backup of their “ITS pseudogene” hypotheses. On the other hand, multiple repetitive ITS copies were identified in 2 additional genome assemblies, JAAVMX000000000 and NGJJ00000000, for the *H. sinensis* Strains IOZ07 and CC1406-203, respectively [50,53], which were available in GenBank after June 2020. Seventeen repetitive ITS1-5.8S-ITS2 copies were found within JAAVMX000000000, 7 repetitive ITS1-5.8S-ITS2 copies within NGJJ00000000 (Table 1), and 2 partial ITS segments within NGJJ00000000 were not included in our analysis.

**Table 1.**
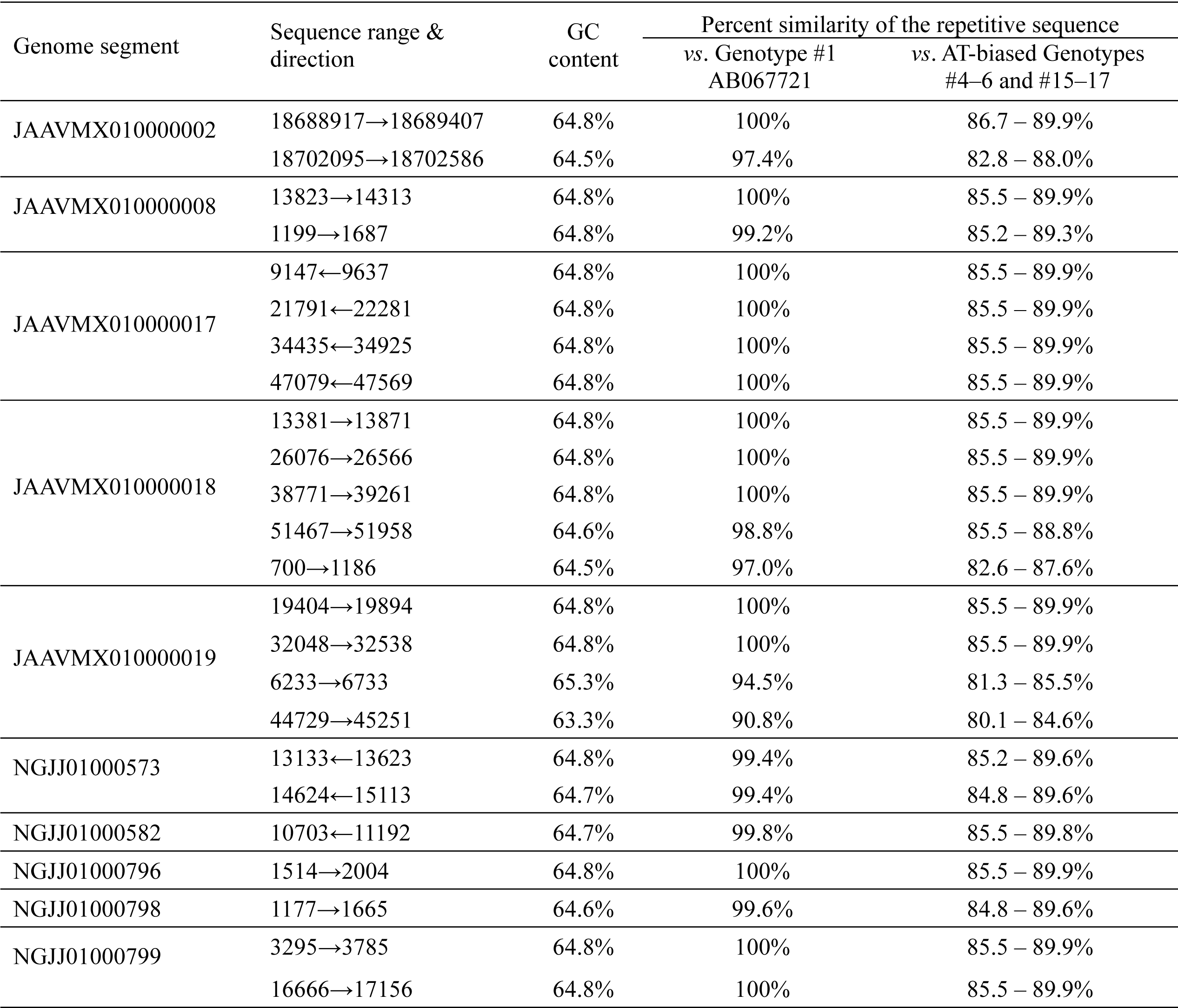
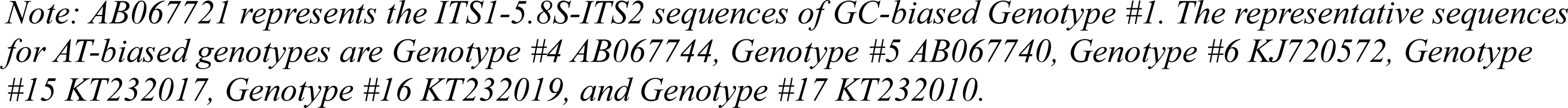
GC content of the repetitive ITS1-5.8S-ITS2 sequences from the genome assemblies of 2 H. sinensis strains and percent similarities of the repetitive ITS sequences to the sequences of GC-biased Genotype #1 (AB067721) and AT-biased Genotypes #4‒6 and #15‒17 of O. sinensis.

Most of the repetitive ITS sequences listed in Table 1 were highly homologous (98.8–100%) to AB067721, the reference sequence of GC-biased Genotype #1 *H. sinensis*. Two variable repetitive copies (6233→6733 and 44729→45251) within JAAVMX010000019 showed 94.5% and 90.8% similarity to AB067721, respectively (Table 1). Two other copies, 18702095→18702586 within JAAVMX010000002 and 700→1186 within JAAVMX010000018, showed similarities of 97.4% and 97.0%, respectively, to AB067721. The repetitive ITS sequences showed 80.1‒89.9% similarity to the AT-biased Genotype #4‒6 and #15‒17 sequences of *O. sinensis* (Table 1).

### III-2. Genetic characteristics of the multiple repetitive ITS sequences in the H. sinensis genome

The repetitive ITS sequences within the genome assemblies JAAVMX000000000 and NGJJ00000000 for the *H. sinensis* Strains IOZ07 and CC1406-203, which are listed in Table 1, and the single genomic ITS copies within ANOV01021709, LKHE01000582, and LWBQ01000008 for the *H. sinensis* Strains Co18, 1229, and ZJB12195, respectively, which are listed in Table 2 [44,50–53], were GC biased; their average GC content was 64.7±0.33% (*cf*. Table 1), nearly identical to the GC content (64.8%) of the AB067721 Genotype #1 sequence but significantly greater (*P*<0.001) than that of the AT-biased Genotypes #4‒6 and #15‒17, *i.e*., 51.1±1.69% (44.8‒53.1%) [8–9]. Apparently, these genetic features of the variable repetitive ITS copies were distinct from those of AT-biased *O. sinensis* genotypes, which included multiple scattered transition alleles.

**Table 2.** Percent similarities between nrDNA ITS1-5.8S-ITS2 sequences of 5 genome assemblies of Genotype #1 H. sinensis and 17 O. sinensis genotype sequences. (Modified from. **Table 1 of [9]).**

| Genotype # and<br>GenBank<br>accession # | Percent similarity between the *H. sinensis* genomic ITS segments and 17 *O. sinensis* genotypes |  |  |  |  |
| --- | --- | --- | --- | --- | --- |
|  | ANOV01021709<br>838→1504 | LKHE01000582<br>2074→2740 | LWBQ01000008<br>991739→992405 | JAAVMX010000017<br>9029←9695 | NGJJ01000799<br>3237→3903 |
| #1 AB067721 | 99.7% | 100% | 99.6% | 100% | 100% |
| #2 MG770309 | 94.4% | 94.7% | 94.7% | 94.7% | 94.7% |
| #3 HM595984 | 95.9% | 96.1% | 95.6% | 95.3% | 95.3% |
| #7 AJ488254 | 94.7% | 95.0% | 95.3% | 95.0% | 95.0% |
| #8 GU246286 | 89.5% | 89.5% | 89.2% | 89.5% | 89.5% |
| #9 GU246288 | 95.0% | 95.2% | 94.6% | 95.2% | 95.2% |
| #10 GU246287 | 83.0% | 83.2% | 83.0% | 83.2% | 83.2% |
| #11 JQ695935 | 78.9% | 79.1% | 78.5% | 79.1% | 79.1% |
| #12 GU246296 | 94.3% | 94.5% | 93.9% | 94.5% | 94.5% |
| #13 KT339190 | 87.5% | 87.5% | 87.2% | 87.5% | 87.5% |
| #14 KT339178 | 89.3% | 89.6% | 89.1% | 89.6% | 89.6% |
| #4 AB067744 | 89.1% | 89.4% | 89.3% | 89.4% | 89.4% |
| #5 AB067740 | 86.3% | 86.7% | 86.2% | 86.7% | 86.7% |
| #6 KJ720572 | 85.5% | 85.5% | 85.5% | 85.5% | 85.5% |
| #15 KT232017 | 89.6% | 89.9% | 89.4% | 89.9% | 89.9% |
| #16 KT232019 | 87.4% | 87.7% | 87.6% | 87.7% | 87.7% |
| #17 KT232010 | 87.7% | 88.1% | 87.5% | 88.1% | 88.1% |

Note: “GT” represents the genotype. GC-biased genotypes are listed in *blue* in the upper panel. AT-biased genotypes are listed in *red* in the lower panel. The GenBank accession numbers are hyperlinked to the GenBank database.

The sequences of GC-biased repetitive ITS copies that were variable (94.5% and 90.8% similarity) in comparison to the reference sequence AB067721 of Genotype #1 predominantly contained insertion/deletion point mutations and included less frequent transversion alleles, while only a few transition alleles were found (Table 3). The variable ITS repeats 6233→6733 and 44729→45251 within JAAVMX010000019 (*cf*. Table 1) contained 22 and 32 insertion/deletion alleles and 4 and 11 transversion alleles, respectively (in blue in Figure 1; Table 3); they contained only 1 and 5 transition alleles, having allelic ratios of 26:1 and 43:5, respectively, for insertion/deletion/transversion *vs*. transition.

**Figure 1.**
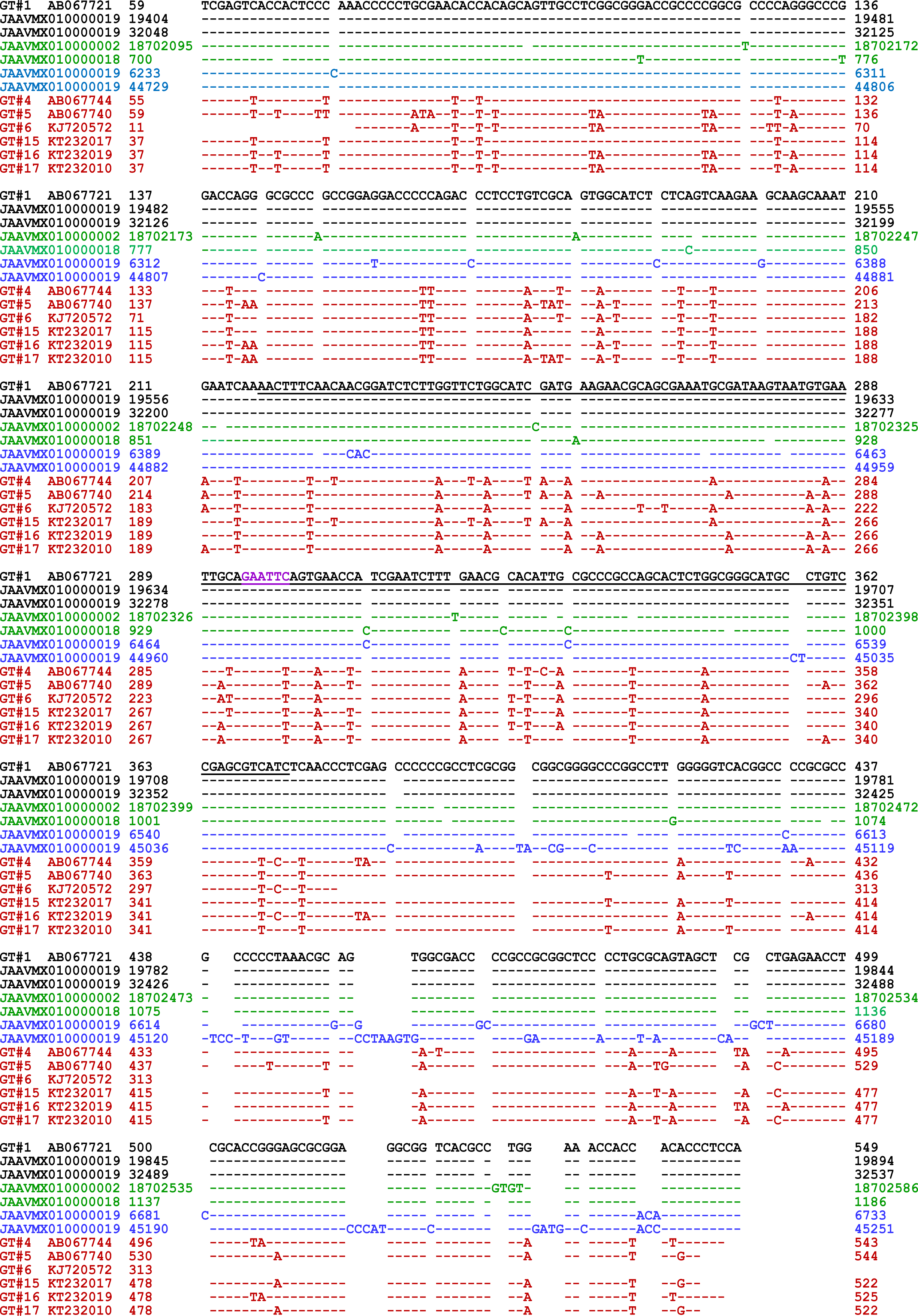
ITS sequence alignment of AB067721, variable and less variable repetitive ITS copies within the genome JAAVMX000000000 of the H. sinensis Strain IOZ07, and AT-biased genotypes of O. sinensis. The ITS sequences contained complete or partial ITS1-5.8S-ITS2 nrDNA segments. “GT” denotes the O. sinensis genotype. The underlined sequence in black represents the 5.8S gene of GC-biased Genotype #1 H. sinensis. AB067721 is the ITS sequence of GC-biased Genotype #1 H. sinensis. The genome assemblies JAAVMX010000002, JAAVMX010000018, and JAAVMX010000019 were obtained from the H. sinensis Strain IOZ07 [50]. One copy each within JAAVMX010000002 and JAAVMX010000018, indicated in green, shares 97.4% or 97.0% similarity with AB067721. JAAVMX010000019 contains 4 repetitive ITS copies, including 2 black sequences (19404→19894 and 32048→32537), which are 100% identical to AB067721, and 2 other blue sequences (6233→6733 and 44729→45251), with 94.5% and 90.8% similarity to AB067721. The sequences in red, namely, AB067744, AB067740, KJ720572, KT232017, KT232019, and KT232010, represent AT-biased Genotypes #4‒6 and #15‒17 of O. sinensis, respectively. The underlined “GAATTC” sequences in purple represent the EcoRI endonuclease cleavage sites in the sequences of GC-biased Genotype #1 and the GC-biased genome assembly JAAVMX000000000. EcoRI endonuclease cleavage sites are absent in AT-biased ITS sequences due to a single-base cytosine-to-thymine (C-to-T) transition. Hyphens indicate identical bases, and spaces denote unmatched sequence gaps.

**Table 3.**
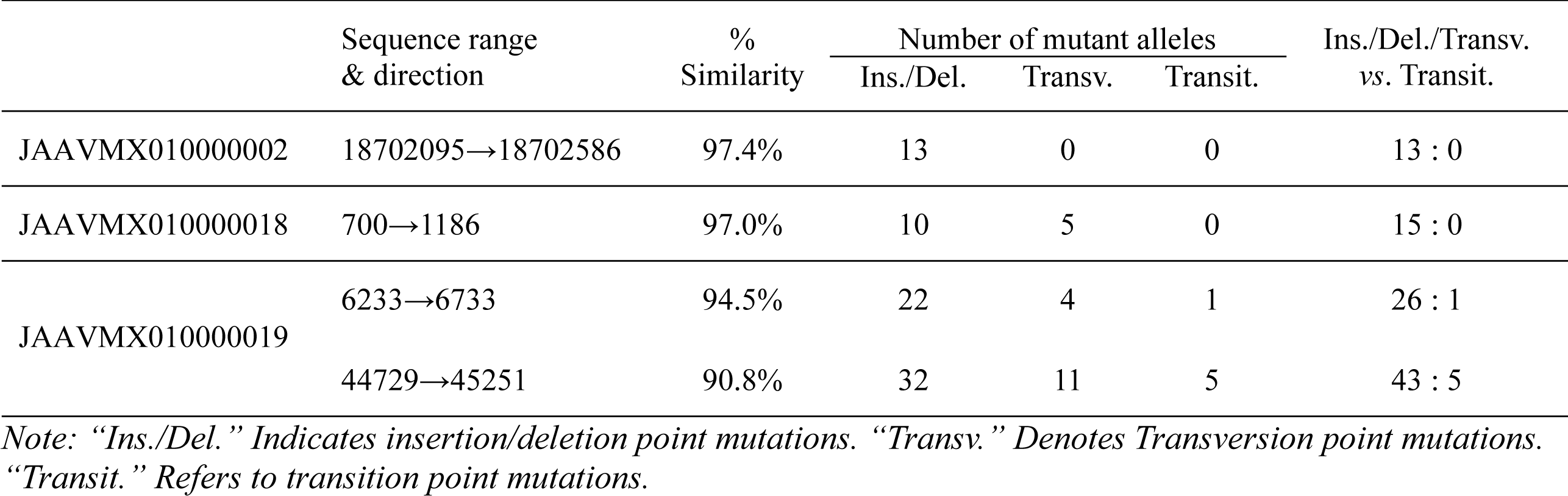
Types of mutations in the variable and slightly less variable repetitive ITS sequences compared with those of GC-biased Genotype #1 H. sinensis (AB067721).

The sequences of repetitive ITS copies, 18702095→18702586 within JAAVMX010000002 and 700→1186 within JAAVMX010000018 (*cf*. Table 1), were less variable (97.4% and 97.0% similarity to AB067721); they contained no transition alleles and predominantly included 13 and 15 insertion, deletion and transversion alleles, with allelic ratios of 13:0 and 15:0, respectively, for insertion/deletion/transversion *vs*. transition (in green in Figure 1; Table 3). These genetic characteristics of the variable and less variable repetitive ITS copies indicate that they were not generated through RIP mutagenesis, which theoretically causes cytosine-to-thymine (C-to-T) and guanine-to-adenine (G-to-A) transition mutations. Other repetitive ITS copies with 98.1‒99.8% similarity to AB067721 are listed in Table 1 and contained only a few insertion/deletion point mutations.

### III-3. Phylogenetic features of multiple repetitive ITS copies in the H. sinensis genome

As shown in Tables 1‒2 and Figure 1, the sequences of GC-biased Genotypes #2–3 and #7–14 and of AT-biased Genotypes #4‒6 and #15‒17 of *O. sinensis* are absent in the genome assemblies ANOV00000000, JAAVMX000000000, LKHE00000000, LWBQ00000000, and NGJJ00000000 of the *H. sinensis* Strains Co18, IOZ07, 1229, ZJB12195, and CC1406-203, respectively, and instead belong to the genomes of independent *O. sinensis* fungi [5–9,16,18,20–22,31,41,44,50–53]. The genomic repetitive ITS copies were phylogenetically clustered into the GC-biased Genotype #1 clade, shown in blue alongside the Bayesian majority rule consensus tree in Figure 2, and were phylogenetically distant from the AT-biased clades containing Genotypes #4‒6 and #15‒17 of *O. sinensis*, shown in red alongside the tree, similar to what has previously been demonstrated in phylogenetic trees that were conferred using different phylogenetic algorithms [5–9,18,20,31,34,41]. Figure 2 includes the 2 variable genomic repetitive ITS copies with low similarity (94.5% and 90.8%) and the 2 other repetitive ITS copies with relatively low similarity (97.4% and 97.0%), which are shown in Tables 1 and 3. The repetitive ITS copies are individually scattered within the GC-biased Genotype #1 clade. The 2 variable repetitive ITS sequences, 6233→6733 (94.5% similarity) and 44729→45251 (90.8% similarity), within the JAAVMX010000019 genome assembly (*cf*. Tables 1 and 3), showed greater phylogenetic distances (horizontal lines) from other GC-biased ITS copies within the GC-biased clades. GC-biased Genotypes #2‒3 and #7‒14 of *O. sinensis* showed greater phylogenetic distances from the sequences of GC-biased Genotype #1 and the repetitive ITS copies. These results indicate that genetic variations in the GC-biased repetitive ITS sequences shown in blue in Figure 1 had a significant impact on the Bayesian phylogenetic clustering analysis.

**Figure 2.**
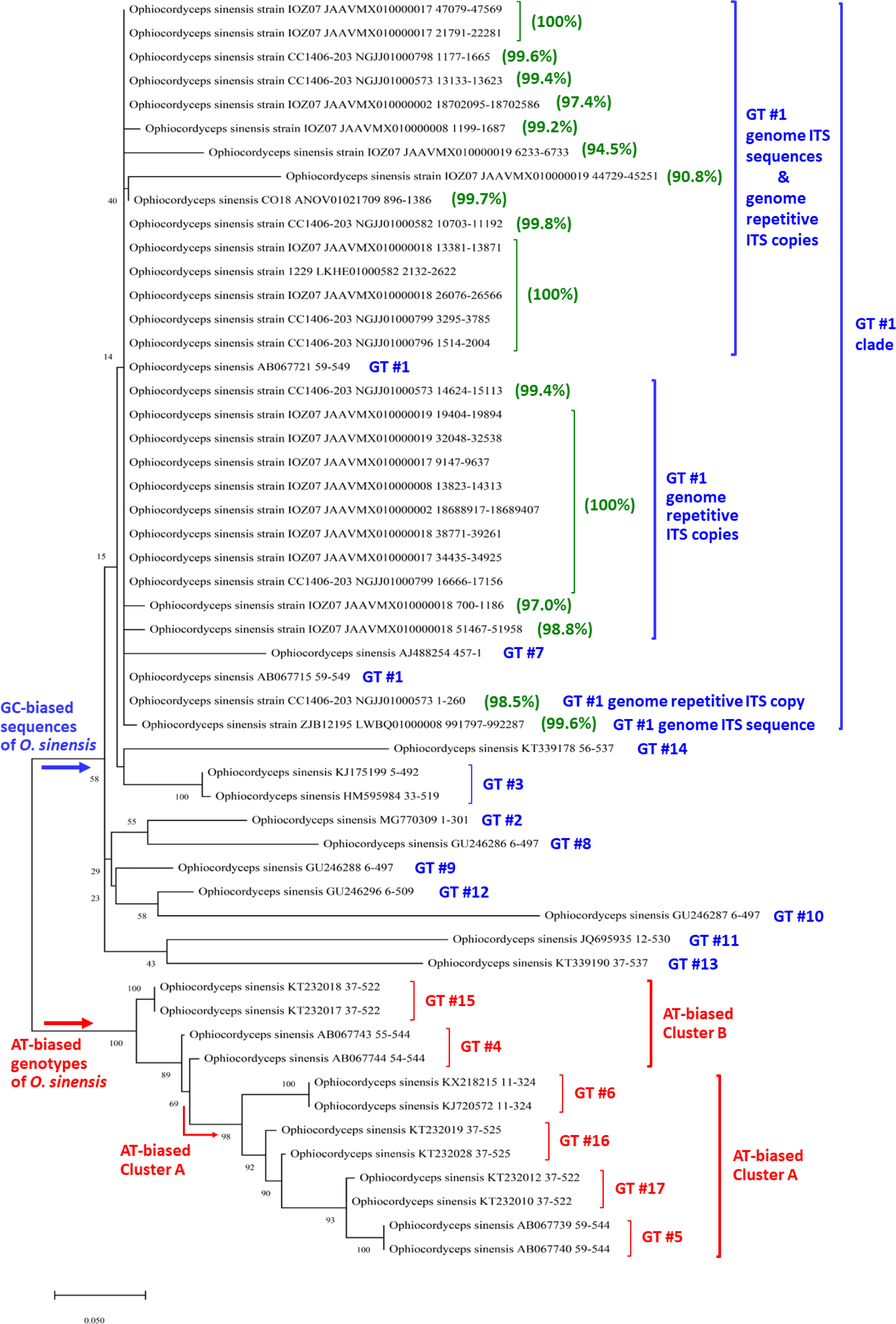
A Bayesian majority rule consensus phylogenetic tree. “GT” represents the genotype. Twenty-eight ITS segments within the genome assemblies (ANOV01021709, LKHE01000582, LWBQ01000008, JAAVMX010000002, JAAVMX010000008, JAAVMX0100000017, JAAVMX0100000018, JAAVMX010000019, NGJJ01000573, NGJJ01000582, NGJJ01000796, NGJJ01000798, and NGJJ01000799) of H. sinensis strains (Co18, 1229, ZJB12195, IOZ07, and CC1406-203, respectively) and 25 ITS sequences of GC-biased Genotypes #1–3 and #7–14 (in blue alongside the tree) and AT-biased Genotypes #4–6 and #15–17 of O. sinensis (in red alongside the tree) were analyzed phylogenetically using MrBayes v3.2.7 software (cf. Section II-4). Genome assemblies JAAVMX000000000 and NGJJ00000000 contain multiple repetitive ITS copies (cf. Table 1). The percent similarities of the genomic sequences of repetitive ITS copies with the representative Genotype #1 sequence (AB067721) are shown in green alongside the tree.

### III-4. Multiple GC-biased genotypes of *O. sinensis* in relation to the number of repetitive genomic ITS copies

In addition to the phylogenetic analysis of multiple genotypes of *O. sinensis* shown in Figure 2, the sequences of GC-biased Genotypes #1‒3 and #7‒12 were further analyzed (Figure 3) and compared with the genomic repetitive ITS copies. GC-biased Genotypes #13‒14 were characterized by large DNA segment reciprocal substitutions and recombination of genetic material between the genomes of the 2 parental fungi, Genotype #1 *H. sinensis* and an AB067719-type fungus [9], and they were not included in the analysis in Figure 3. Figure 2 shows that GC-biased Genotypes #2‒3 and #8‒14 formed several GC-biased phylogenetic branches outside the Genotype #1 clade with great phylogenetic distances (horizontal lines), and they were phylogenetically and genetically distinct from the genomic repetitive ITS copies of Genotype #1 *H. sinensis* (*cf*. Table 2 and Figure 3). Genotype #7 was unique among the GC-biased genotypes and was placed within the Genotype #1 clade with great phylogenetic distance (horizontal line) in the Bayesian tree in Figure 2. The sequences of Genotype #7 were 94.7‒95.3% similar to Genotype #1, which had repetitive ITS sequences within the genome assemblies JAAVMX000000000 and NGJJ00000000, which had numerous mutant alleles (*cf*. Table 2, Figure 3).

**Figure 3.**
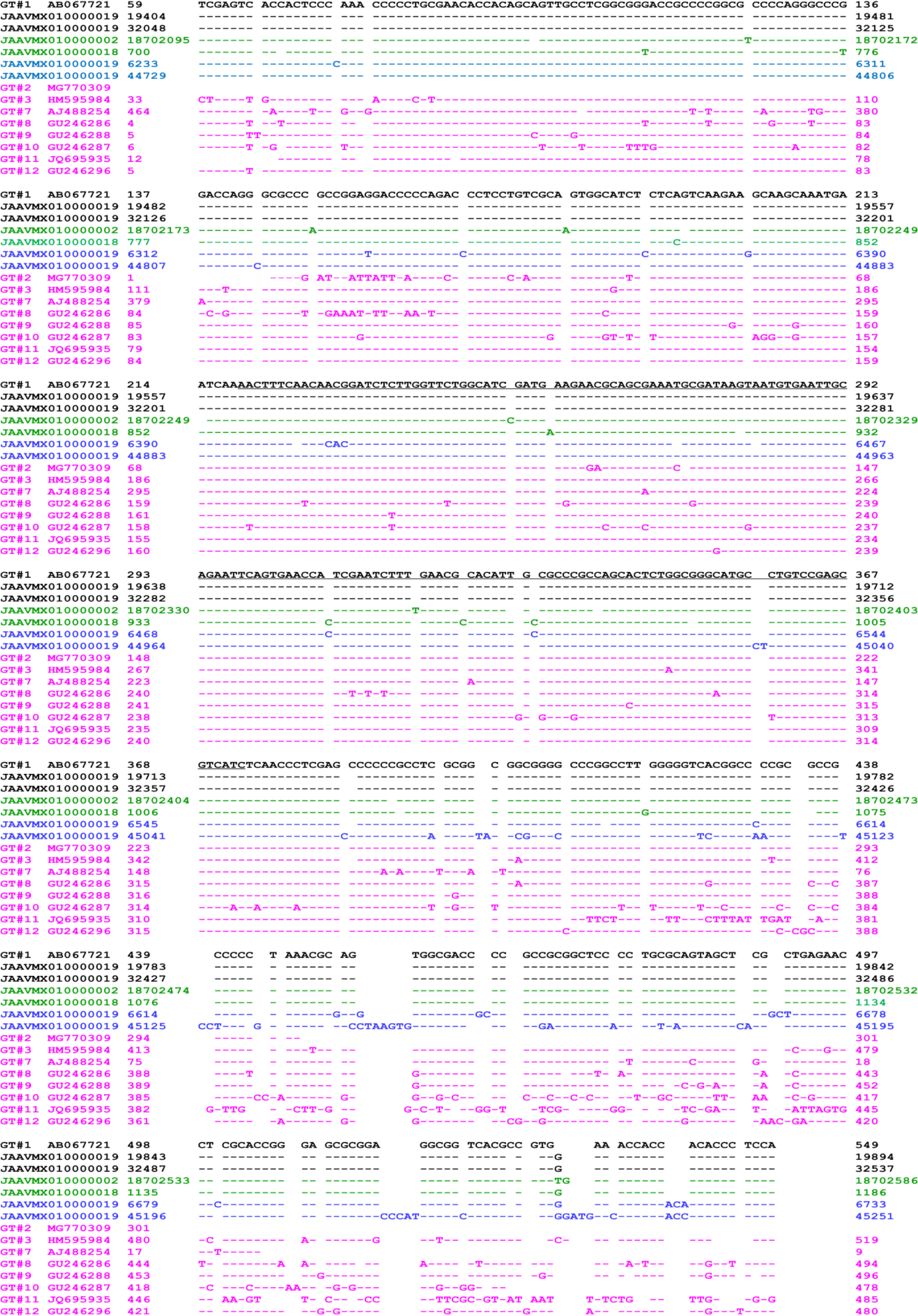
Alignment of repetitive ITS copies within the genome JAAVMX000000000 of the H. sinensis Strain IOZ07 and ITS sequences of GC-biased genotypes of O. sinensis. The genome segments JAAVMX010000002, JAAVMX010000018, and JAAVMX010000019 were obtained from the H. sinensis Strain IOZ07 [50]. One copy each within JAAVMX010000002 and JAAVMX010000018, indicated in **green**, shares 97.4% or 97.0% similarity with AB067721. JAAVMX010000019 contains 4 repetitive ITS copies, including 2 sequences in **black** (19404→19894 and 32048→32537), which are 100% identical to AB067721, and 2 other sequences in **blue** (6233→6733 and 44729→45251), with 94.5% and 90.8% similarity to AB067721. “GT” denotes the O. sinensis genotype. AB067721, MG770309, HM595984, AJ488254, GU246286, GU246288, GU246287, JQ695935, and GU246296 are the ITS sequences in pink of GC-biased Genotypes #1‒3 and #7‒12 of O. sinensis, respectively. The underlined sequence in **black** represents the 5.8S gene of GC-biased Genotype #1 H. sinensis. Hyphens indicate identical bases, and spaces denote unmatched sequence gaps.

The Genotype #7 sequence (AJ488254) was originally identified in the stroma of a *C. sinensis* specimen (H1023) collected from Qinghai Province, China and uploaded to GenBank by Chen *et al*. [16]. The authors identified the GC-biased Genotype #1 ITS sequence (AJ488255) from the tail of the caterpillar body of the same specimen, leading to questions regarding whether the sequences of AJ488254 and AJ488255 coexisted in a single genome of GC-biased Genotype #1 *H. sinensis* as 2 repetitive ITS copies or belonged to the genomes of independent *O. sinensis* fungi cooccurring in different compartments of the same *C. sinensis* specimen. The study conducted by Chen *et al*. [16] did not involve the purification of the *O. sinensis* strain(s) within the same natural specimen. Thus, there is no evidence to date for determining whether the sequence (AJ488254) of Genotype #7 is indeed present in the genome of Genotype #1 *H. sinensis* as one of the repetitive ITS copies, although Genotype #7 was placed within the Genotype #1 phylogenetic clade shown in Figure 2 and was phylogenetically distant from GC-biased Genotypes #2–3 and #8–14, which were placed outside the Genotype #1 clade in the Bayesian tree shown in Figure 2.

Notably, GC-biased Genotypes #1 and #2 were simultaneously detected in the stromata of the immature, maturing and mature *C. sinensis* samples collected from Kangding County in Sichuan Province [42]. These 2 genotypes showed completely different developmental patterns in an asynchronous, disproportional manner in *C. sinensis* stromata during maturation [9,24,42], indicating the genomic independence of the 2 GC-biased genotypes. The Bayesian tree shown in Figure 2 confirms that the sequences of Genotypes #1 and #2 were placed in separate GC-biased phylogenetic clades and were unlikely to be represented as repetitive ITS copies in a single *O. sinensis* genome.

### III-5. Multilocus analysis of repetitive copies of authentic genes in the *H. sinensis* genome and transcriptome

To explore the hypothesis that repetitive AT-biased pseudogenes are present in a single genome of GC-biased *H. sinensis* and the postulated cause of “RIP” mutagenesis, we extended the analysis of the single ITS locus to multiple loci of the *H. sinensis* genome and transcriptome. Unfortunately, GenBank does not provide annotations for authentic genes in the genome assemblies (ANOV00000000, JAAVMX000000000, LKHE00000000, LWBQ00000000, and NGJJ00000000) of the *H. sinensis* Strains Co18, IOZ07, 1229, ZJB12195, and CC1406-203, respectively. Cross-referencing based on gene annotations for one (JAACLJ010000002) of the 13 genome contigs of *O. camponoti-floridani* Strain EC05 [57], which contains 5,126,525 bp for 1271 authentic genes, the *H. sinensis* genes and their repetitive genomic copies were then positioned and annotated in the genome assemblies of 5 *H. sinensis* strains, and their transcription was analyzed in the mRNA transcriptome of the *H. sinensis* Strain L0106. Table 4 outlines the results of the multilocus analysis of 1271 *H. sinensis* genes, 1167 (91.8%) of which had no repetitive genomic copies, 37 (2.9%) of which had repetitive copies in the genome of only one *H. sinensis* strain, and 67 (5.3%) of which had multiple repetitive copies in the genomes of 5 *H. sinensis* strains.

**Table 4.** Summary of 1271 authentic genes and their repetitive copies in the genomes of the H. sinensis Strains 1229, CC1406-203, Co18, IOZ07, and ZJB12195.

|  | # of genes |  | Change in the AT content | # of genes having repetitive copies |
| --- | --- | --- | --- | --- |
| No genomic repetitive copies | 1167 (91.8%) |  |  |  |
| Single repetitive copy in only one of 5 *H. sinensis* genomes | 37 (2.9%) | Homologous ($\geq 97\%$ similarity) | | 36 |
|  |  | 96.5% similarity | ↓ 0.2% | 1 |
| Multiple repetitive copies in genomes of 5 *H. sinensis* strains | 67 (5.3%) | Essentially no change | ↑ or ↓ within $\pm 1\%$ | 6 |
| | | Slight decreases | ↓ $\leq 5\%$ | 11 |
| | | Large decreases | ↓ $> 5\%$ | 7 |
| | | Slight increases | ↑ $\leq 5\%$ | 17 |
| | | Large increases | ↑ $> 5\%$ | 11 |
|  |  | Bidirectional changes |  | 15 |
| Total: | 1271 |  |  |  |

Note: “↑” and “↓” indicate an increase or decrease in the AT content, respectively.

The 37 authentic genes that had repetitive copies in only one *H. sinensis* genome were normally transcribed, and the transcripts were found in the mRNA transcriptome of the *H. sinensis* Strain L0106. The repetitive copies of 36 of the 37 genes were homologous to the authentic genes. However, one gene (32562→34036 within LWBQ01000158) encoding a PNGase family protein had a single repetitive copy (414324←415851 within LWBQ01000021) in the genome of the *H. sinensis* Strain ZJB12195; this repetitive sequence showed 96.5% similarity to the authentic gene with a 53-nt insertion but no transition or transversion point mutations. The AT content did not change substantially, decreasing from 35.1% in the authentic gene to 34.9% in the repetitive copy. The authentic gene with an 87-nt intron (33112→33198 within LWBQ01000158) was normally transcribed (transcript 610→1996 within GCQL01000547 in the transcriptome of the *H. sinensis* Strain L0106), which encodes a PNGase family protein (accession #EQL03505 containing 462 aa residues, according to the GenBank protein annotation for the genome of the *H. sinensis* Strain Co18). The repetitive copy was also transcribed, and the transcript 1159←1996 within GCQL01000547 had an extra deletion (53 nt) in addition to the 87-nt intron (Figure 4) and encoded the same PNGase family protein but missing 180 aa residues at the N-terminus (181→458 of EQL03505). EQL03505 showed 73.8–82.8% similarity to the PNGase family proteins of 48 other fungi.

**Figure 4.**
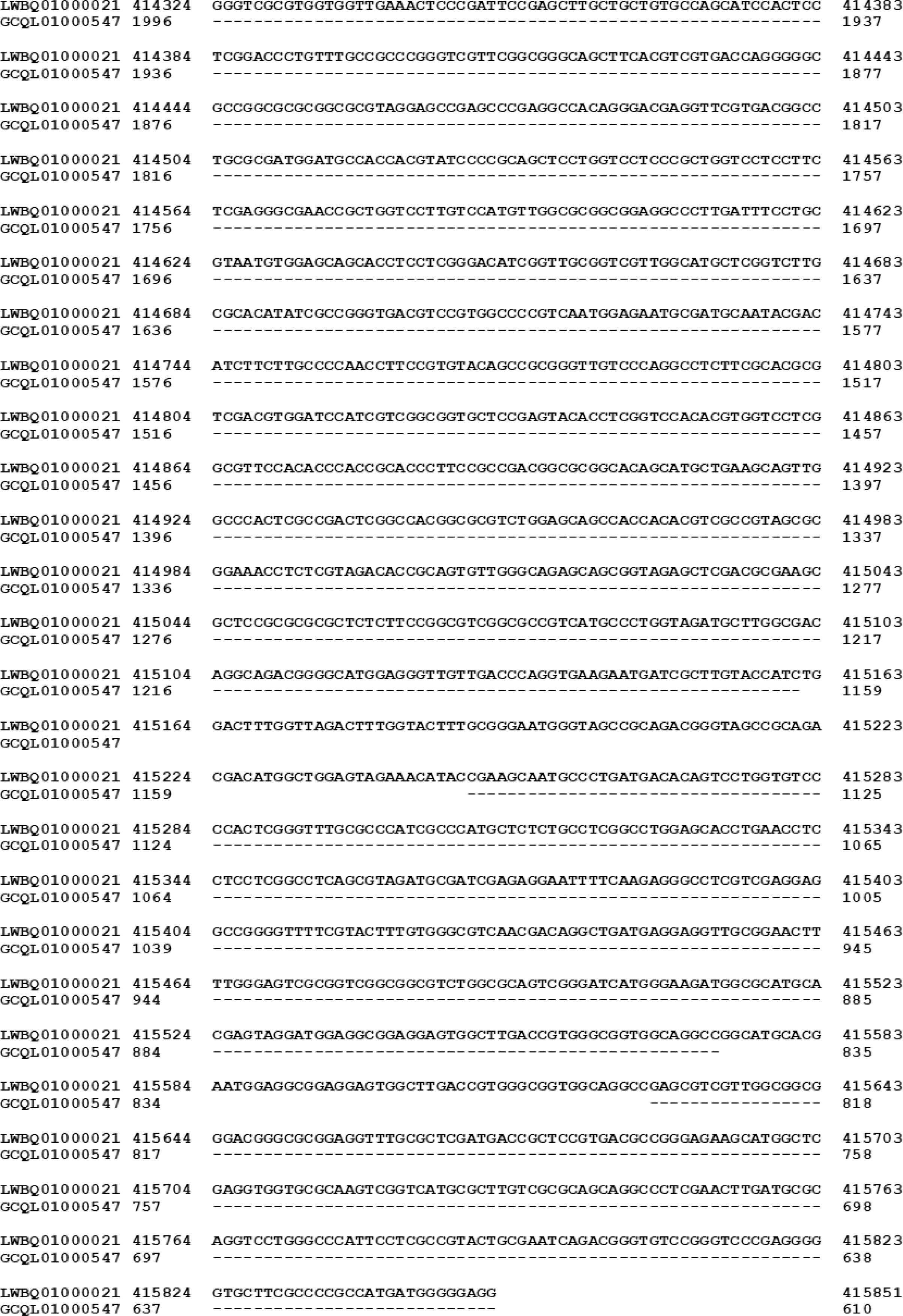
Alignments of the genomic sequence of the repetitive copy (414324←415851 within LWBQ01000021) of the H. sinensis Strain ZJB12195 and the transcriptomic sequence (610←1996 within GCQL01000547) of the Strain L0106 for the PNGase family protein [53–54]. Hyphens indicate identical bases, and spaces denote unmatched sequence gaps.

The remaining 67 authentic genes in the genomes of the *H. sinensis* Strains 1229, CC1406-203, Co18, IOZ07, and ZJB12195 had multiple repetitive copies in the *H. sinensis* genome assemblies with various changes (decreases, increases, or bidirectional changes) in the AT content.

#### III-5.1. Slight decreases in the AT content of repetitive genomic copies

Two of the 18 authentic genes (the query sequences) shown in Table 4 had multiple repetitive genomic copies (the subject sequences) with slight decreases (**↓** ≤5) or large decreases (**↓** >5) in the AT content in comparison with the sequences of authentic genes for the triose-phosphate transporter (Table 5) and heavy metal tolerance protein precursor (Table 6), respectively.

**Table 5:**
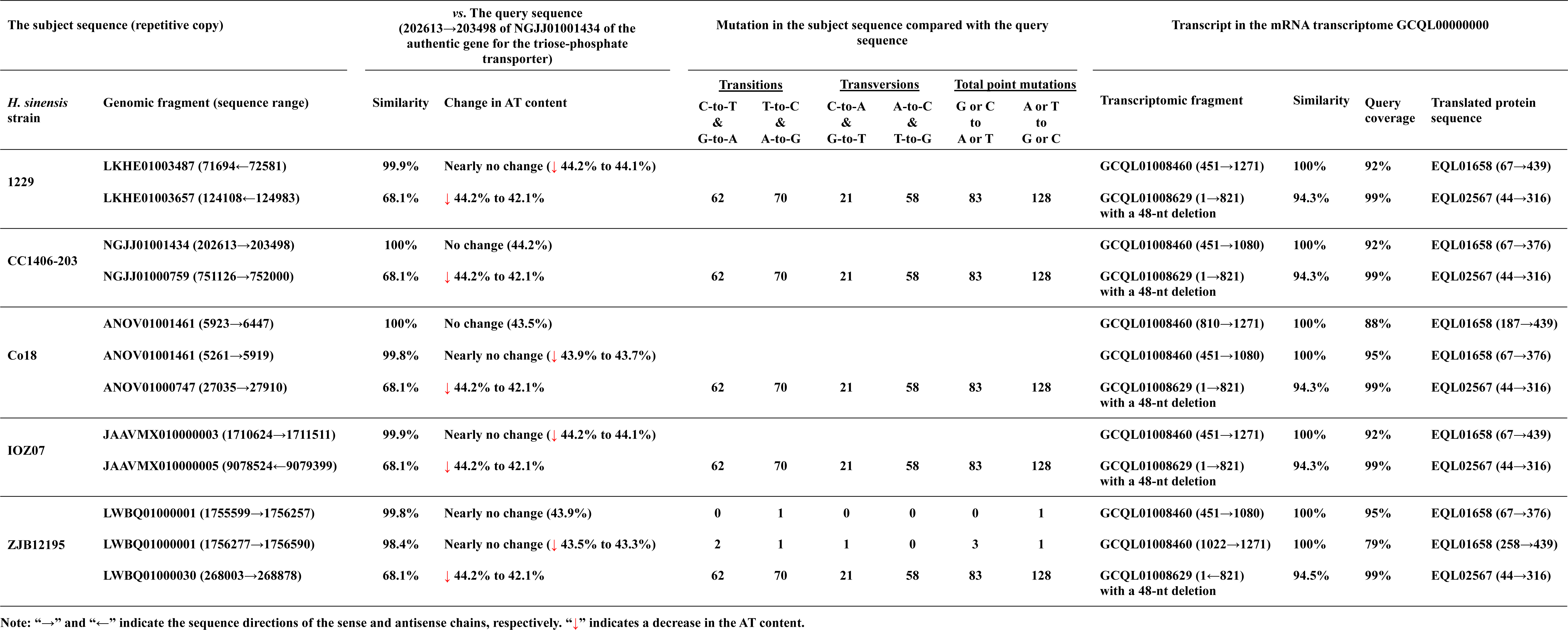
The authentic H. sinensis genes for the triose-phosphate transporter (the query sequence) and the repetitive genomic copies (the subject sequences) with slight decreases (≤ 5%) in the AT content.

**Table 6:**
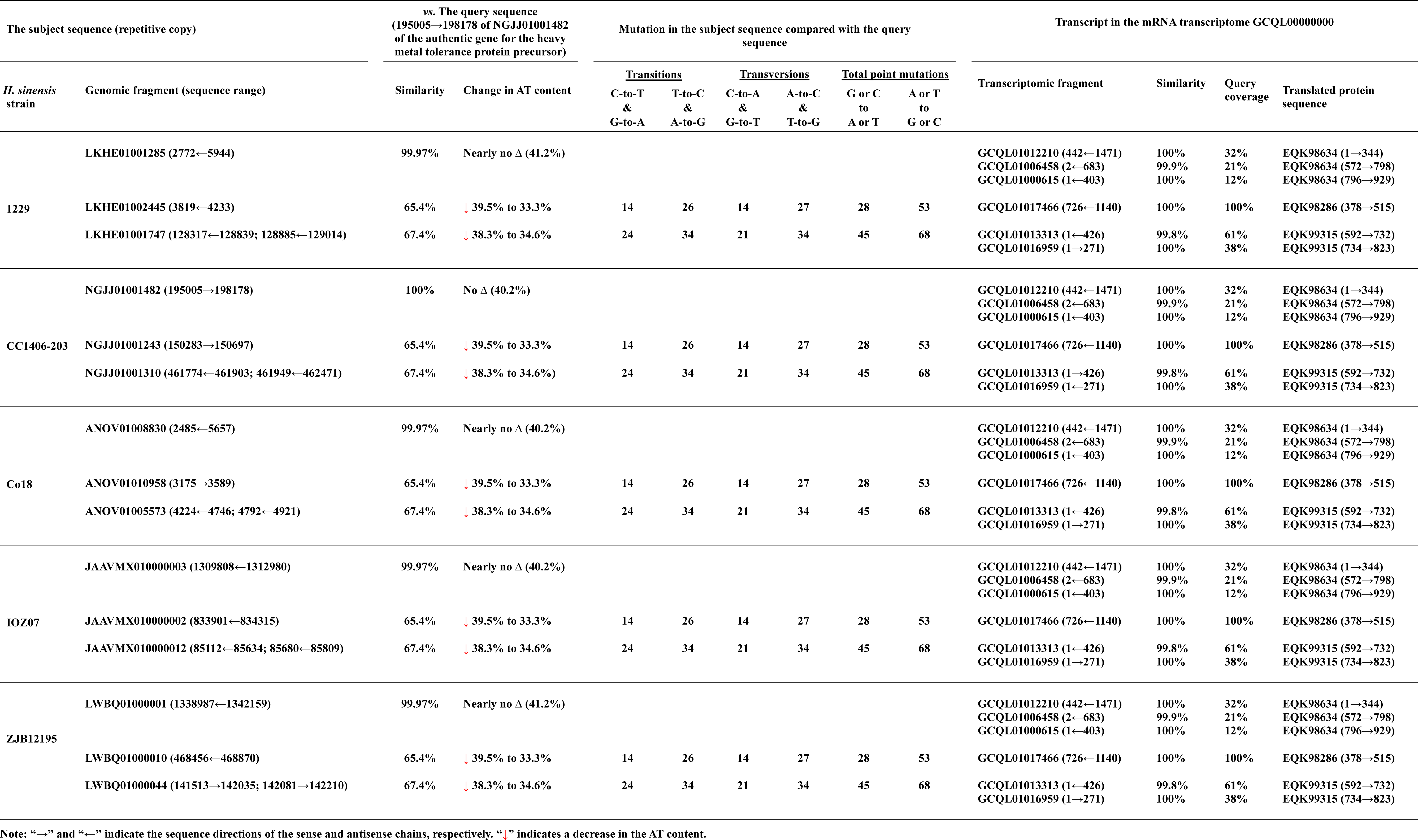
Authentic H. sinensis genes for the heavy metal tolerance protein precursor (the query sequence) and repetitive genomic copies whose AT content decreased markedly (by > 5%).

Table 5 shows that repetitive copies (124108←124983 within LKHE01003657; 751126→752000 within NGJJ01000759; 27035→27910 within ANOV01000747; 9078524←9079399 within JAAVMX010000005; and 268003→268878 within LWBQ01000030) of the authentic gene for the triose- phosphate transporter (71694←72581 within LKHE01003487; 202613→203498 within NGJJ01001434; 5923→6447 and 5261→5919 within ANOV01001461; 1710624→1711511 within JAAVMX010000003; and 1755599→1756257 and 1756277→1756590 within LWBQ01000001, respectively) had slightly reduced AT content (**↓** 2.1%), which was attributed to a greater number of T-to-C and A-to-G transitions (70) and T-to-G and A-to-C transversions (58) than of postulated RIP mutagenesis-related C-to-T and G-to- A transitions (62) and G-to-T and C-to-A transversions (21). Overall, the number (128) of A or T to G or C mutations was much greater than the number (83) of G or C to A or T mutations in the repetitive copies in the 5 *H. sinensis* genomes, resulting in 68.1% sequence similarity and a reduction in the AT content from 44.2% in the authentic genes to 42.1% in the repetitive copies.

The authentic genes for the triose-phosphate transporter (the genome segments within ANOV01001461, JAAVMX010000003, LKHE01003487, LWBQ01000001, and NGJJ01001434), shown in Table 5, were 98.4–100% homologous and were normally transcribed. The transcript (451→1271 within GCQL01008460) in the transcriptome of the *H. sinensis* Strain L0106 was 100% homologous to the gene for the protein (accession #EQL01658 containing 440 aa residues), with query coverages of 79–99%. EQL01658 was defined as the triose-phosphate transporter according to the GenBank protein annotation for the genome of the *H. sinensis* Strain Co18 and showed 61.6–80.8% similarity to the triose-phosphate transporter of 29 other microbes (*Colletotrichum* sp., *Coniochaeta ligniaria*, *Diplogelasinospora grovesii*, *Leotiomycetes* sp., *Phialemonium atrogriseum*, *Ophiocordyceps* sp., *Rhexocercosporidium* sp., *Thozetella* sp.).

The repetitive copies (the genome segments within ANOV01000747, LKHE01003657, JAAVMX010000005, LWBQ01000030, and NGJJ01000759; *cf*. Table 5) were also normally transcribed. The transcript (1←821 within GCQL01008629 of the *H. sinensis* Strain L0106) for the triose-phosphate transporter (accession #EQL02567 containing 330 aa residues) was 94.3–94.5% similar to the repetitive copy sequences with a 48-nt deletion, with a query coverage of 99%. Both EQL01658 and EQL02567, which were encoded by the authentic genes and their repetitive copies, respectively, were annotated as the triose-phosphate transporter; however, these 2 protein sequences were only 65.4% similar to each other, with a query coverage of 89%. EQL02567, encoded by the *H. sinensis* genomic repetitive copies, exhibited (1) 80.3% similarity to the sequence XP_044716728 for eamA-like transporter family domain-containing protein of *Hirsutella rhossiliensis*, (2) 61.3–67.4% similarity to 20 protein sequences of the solute carrier family 35 member E3 proteins of many microbes (*Colletotrichum* sp., *Coniochaeta* sp., *Diaporthaceae* sp., *Hymenoscyphus varicosporioides*, *Lasiosphaeria hispida*, *Pyricularia oryzae*, *Tolypocladium ophioglossoides*, *Tolypocladium paradoxum*), and (3) 59.7–68.4% similarity to 26 protein sequences of the triose-phosphate transporters of many other microbes (*Amylocarpus encephaloides*, *Colletotrichum* sp., *Coniochaeta ligniaria*, *Diplogelasinospora grovesii*, *Phialemonium atrogriseum*, *Rhexocercosporidium* sp., *Thozetella* sp., *Xylariaceae* sp., *Zalerion maritima*).

Among the 11 authentic genes outlined in Table 4 that had multiple repetitive copies with slightly reduced AT content (**↓** ≤5), the repetitive copies of 4 authentic genes had mutagenetic patterns similar to those of triose-phosphate transporters, with a combination of more RIP-related C-to-T than T-to-C transitions but more A-to-G than RIP-related G-to-A transitions, or *vice versa*. The repetitive copies of 6 other authentic genes had consistently more T-to-C and A-to-G transitions than the postulated RIP-related C-to-T and G-to-A transitions, in addition to numerous transversion, insertion, and deletion point mutations.

#### III-5.2. Large decreases in the AT content of repetitive genomic copies

Table 6 shows the authentic genes for the heavy metal tolerance protein precursor (2772←5944 within LKHE01001285; 195005→198178 within NGJJ01001482; 2485←5657 within ANOV01008830; 1309808←1312980 within JAAVMX010000003; and 1338987← 1342159 within LWBQ01000001), which had multiple repetitive copies with large decreases (**↓** >5) in the AT content. The repetitive copies of the authentic gene could be divided into 2 groups.

1. The first group of repetitive copies (3819←4233 within LKHE01002445; 150283→150697 within NGJJ01001243; 3175→3589 within ANOV01010958; 833901←834315 within JAAVMX010000002; and 468456←468870 within LWBQ01000010) in the genomes of the *H. sinensis* Strains 1229, CC1406-203, Co18, IOZ07, and ZJB12195, respectively, were 65.4% similar to the authentic gene. These repetitive copies had many more T-to-C and A-to-G transitions (26) and A-to-C and T-to-G transversions (27) than postulated RIP-related C-to-T and G-to-A transitions (14) and C-to-A and G-to-T transversions (14). Overall, the total number (28) of G or C to A or T point mutations was much fewer than the number (53) of A or T to G or C mutations in the repetitive copies, resulting in largely reduced AT content (↓ 6.2%), from 39.5% in the authentic gene to 33.3% in the repetitive copies.
2. The second group of repetitive copies (128317←128839 and 128885←129014 within LKHE01001747; 461774←461903 and 461949←462471 within NGJJ01001310; 4224←4746 and 4792←4921 within ANOV01005573; 85112←85634 and 85680←85809 within JAAVMX010000012; and 141513→142035 and 142081→142210 within LWBQ01000044) of the authentic gene in the 5 *H. sinensis* genome assemblies showed 67.4% similarity to the authentic genes. There were more T-to-C and A-to-G transitions (34) and A-to-C and T-to-G transversions (34) than postulated RIP-related C-to-T and G-to-A transitions (24) and C-to-A and G-to-T transversions (21). Overall, this group of repetitive copies had a total of 45 G or C to A or T point mutations, which was much fewer than the number (68) of A or T to G or C point mutations in the repetitive copies, resulting in slightly reduced AT content (↓ 3.7%), from 38.3% in the authentic gene to 34.6% in the repetitive copies.

Additionally, as shown in Fig. 6, the authentic genes (the genome segments within ANOV01008830, JAAVMX010000003, LKHE01001285, LWBQ01000001, and NGJJ01001482) were normally transcribed. The fragmented transcripts (GCQL01012210, GCQL01006458, and GCQL01000615 of the *H. sinensis* Strain L0106) were 99.9–100% identical to the authentic gene for the heavy metal tolerance protein precursor (accession #EQK98634 containing 1029 amino acids), with query coverages of 12–32%. EQK98634 showed 74.0–82.3% similarity to the heavy metal tolerance protein precursor of 19 other microbes (*Drechmeria coniospora*, *Fusarium* sp., *Hapsidospora chrysogenum*, *Metarhizium* sp., *Moelleriella libera*, *Ophiocordyceps* sp., *Pochonia chlamydosporia*, *Purpureocillium* sp., *Tolypocladium* sp.).

**Figure 5.**
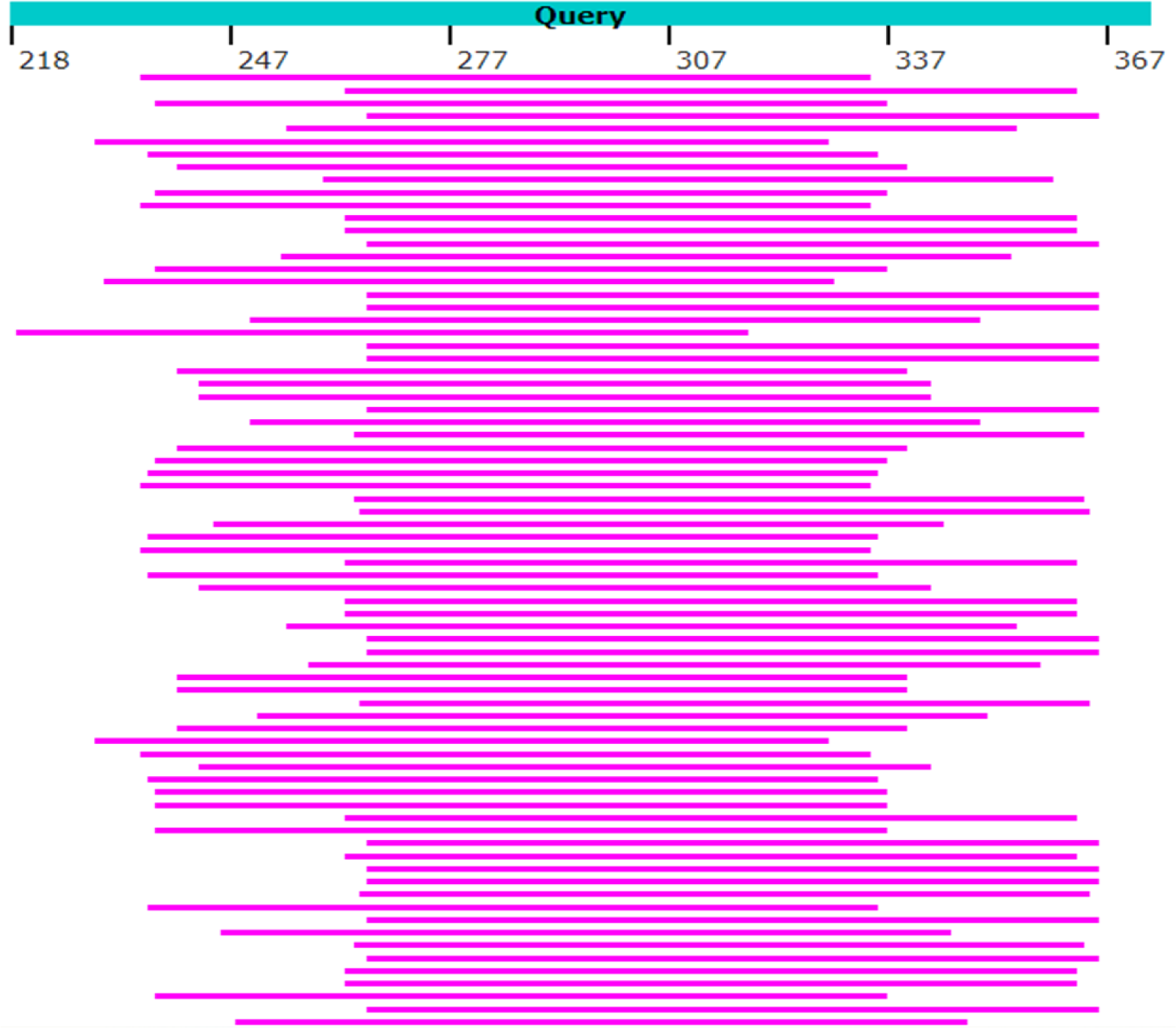
**Graphic alignment summary of the query sequence (in blue), the 5.8S gene (218‒373 segment of Genotype #1 H. sinensis AB067721), and the first 75 in the list of 1894 overlapping shotgun reads (pink bars; GenBank SRR5428527) of mature natural C. sinensis.**

**Figure 6.**
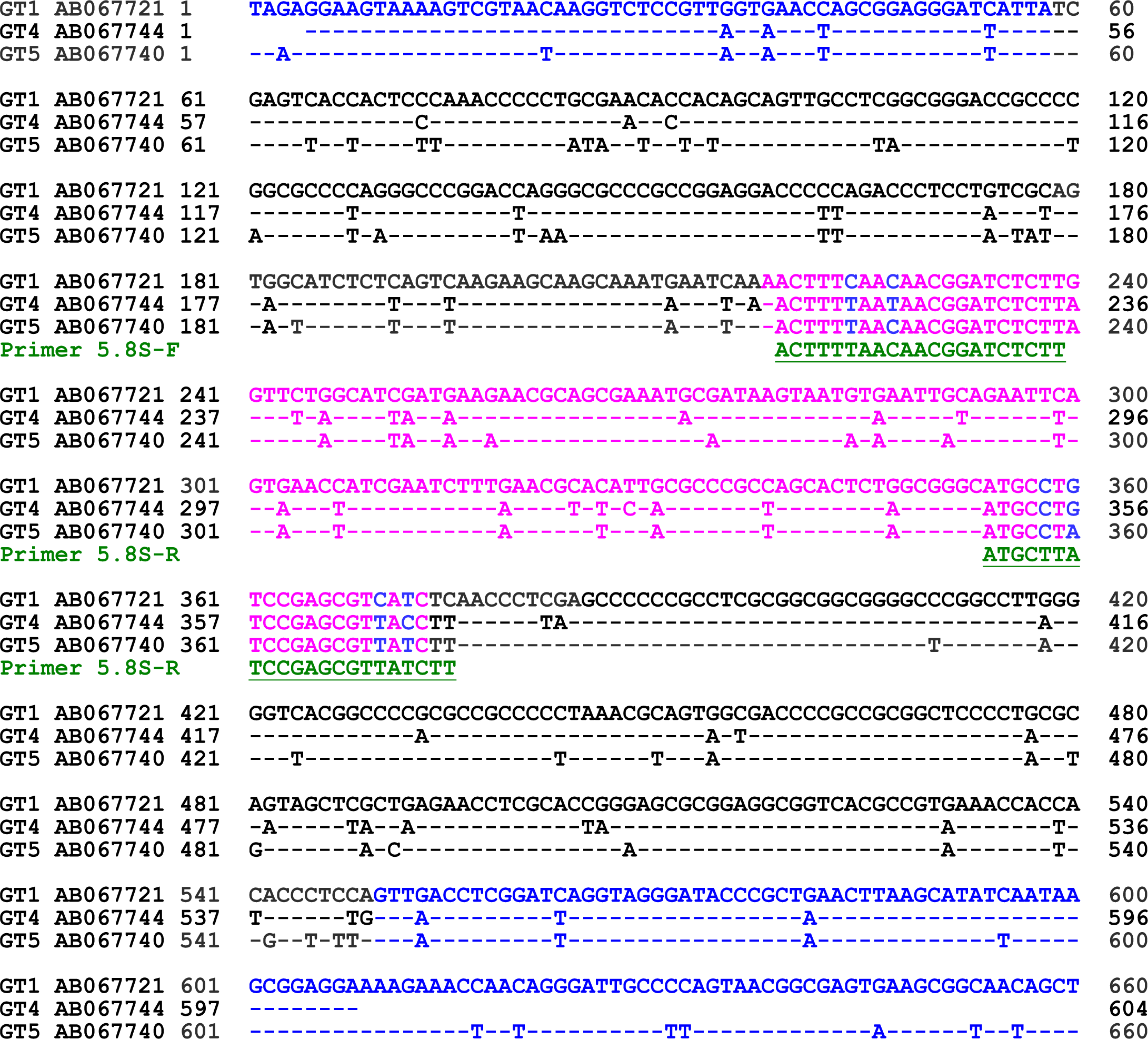
ITS sequence alignment of AB067721 of GC-biased Genotype #1, AB067744 and AB067740 of AT-biased Genotypes #4‒5 of O. sinensis and the locations of 5.8S-F/R primers. “GT1”, “GT4”, and “GT5” represent Genotypes #1 and #4‒5 of O. sinensis, respectively. The sequences in blue are from the 18S and 28S genes (5’ and 3’ end regions, respectively). The sequences in pink represent the sequences of the 5.8S genes. ITS1 is shown in black between the 18S and 5.8S genes, and ITS2 is shown in black between the 5.8S and 28S genes. The sequences underlined in green are for the primers 5.8S-F/R, while primer 5.8S-R was converted to the sense chain sequence. The hyphens indicate identical bases.

The first group of repetitive copies (the genome segments within ANOV01010958, JAAVMX010000002, LKHE01002445, LWBQ01000010, and NGJJ01001243) was 65.4% similar to the authentic gene and was normally transcribed (*cf*. Table 6). The transcript GCQL01017466 was 100% homologous to the DNA sequences of repetitive copies, with a query coverage of 100%; however, it presumptively encoded a different protein, the transporter ATM1 (accession #EQK98286, which contains 598 amino acids), according to the GenBank protein annotation for the genome of the *H. sinensis* Strain Co18. EQK98286 was 84.8–89.4% similar to 59 protein sequences of the transporter ATM1 or the mitochondrial iron–sulfur cluster transporter ATM1 of different microbes (*Akanthomyces lecanii*, *Beauveria* sp., *Colletotrichum* sp., *Conoideocrella luteorostrata*, *Cordyceps* sp., *Dactylonectria macrodidyma*, *Fusarium* sp., *Ilyonectria robusta*, *Lecanicillium* sp., *Mariannaea* sp., *Metarhizium* sp., *Neonectria ditissima*, *Ophiocordyceps* sp., *Paramyrothecium foliicola*, *Purpureocillium* sp., *Tolypocladium* sp., *Trichoderma* sp.).

The second group of repetitive copies (the genome segments within ANOV01005573, JAAVMX010000012, LKHE01001747, LWBQ01000044, and NGJJ01001310) was 67.4% similar to the authentic genes and was also normally transcribed (*cf*. Table 6). The fragmented transcripts (GCQL01013313 and GCQL01016959) were 99.8–100% homologous to the repetitive copies, with query coverages of 38–61%; however, the fragmented transcripts presumably encoded a different protein, the ABC transporter (accession #EQK99315, containing 844 amino acids), according to the GenBank protein annotation for the genome of the *H. sinensis* Strain Co18. EQK99315 was 50.2–89.0% similar to 22 other ABC transporter protein sequences of many different microbes (*Aspergillus* sp., *Beauveria bassiana*, *Blastomyces* sp., *Chaetomium strumarium*, *Histoplasma* sp., *Penicillium* sp., *Purpureocillium* sp., *Ophiocordyceps* sp.).

Like in the case of the mutagenetic pattern of the authentic gene encoding the heavy metal tolerance protein precursor, the repetitive copies of 6 other genes whose AT content decreased substantially (↓ >5%) (*cf*. Table 4) were associated with consistently greater numbers of T-to-C and A-to-G transitions than postulated RIP-related C-to-T and G-to-A transitions, in combination with numerous transversion, insertion and deletion point mutations.

#### III-5.3. Slight increases in the AT content of repetitive genomic copies

The authentic genes for the maltose permease and lipase/serine esterase are shown in Tables 7–8 as representatives of the 28 authentic genes that had multiple repetitive copies with slight increases (↑ ≤5%; Table 7) and large increases (↑ >5%; Table 8) in the AT content, respectively.

**Table 7:**
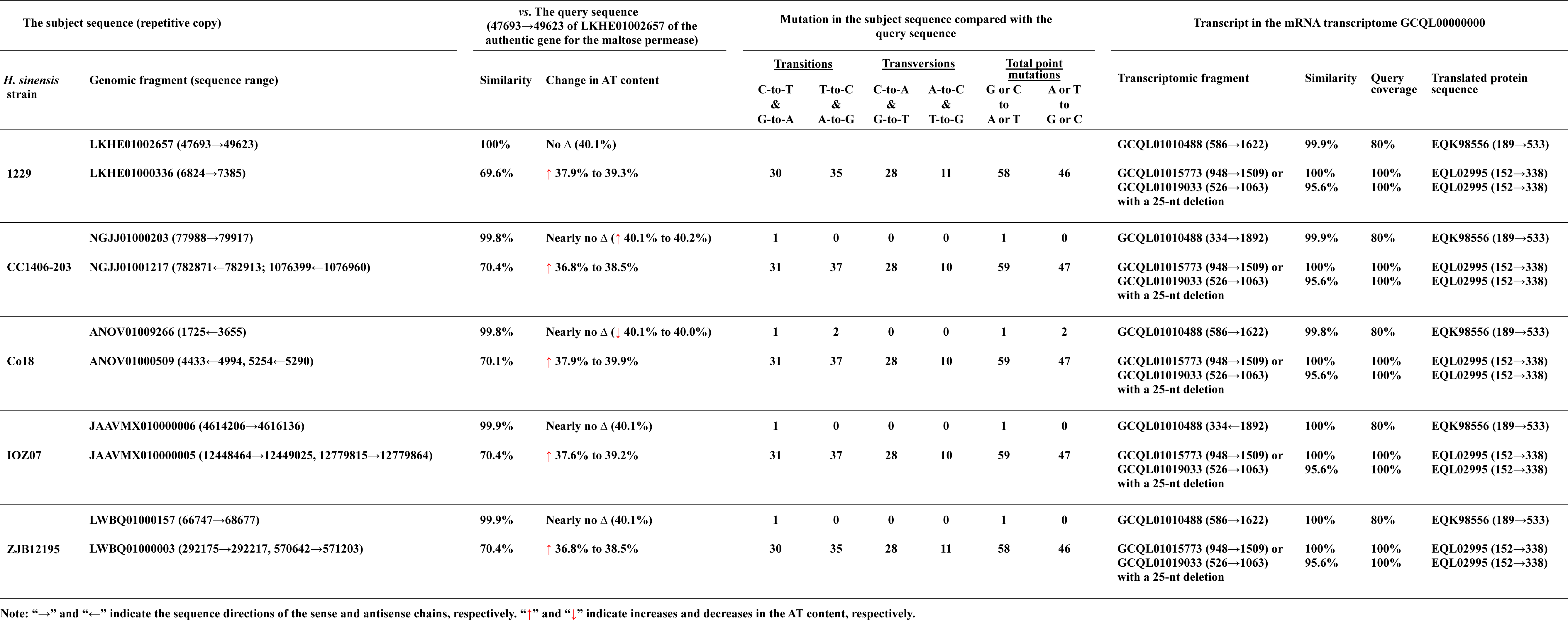
Authentic H. sinensis genes for the maltose permease (query sequence) and repetitive genomic copies with minor increases (≤ 5%) in the AT content.

**Table 8:**
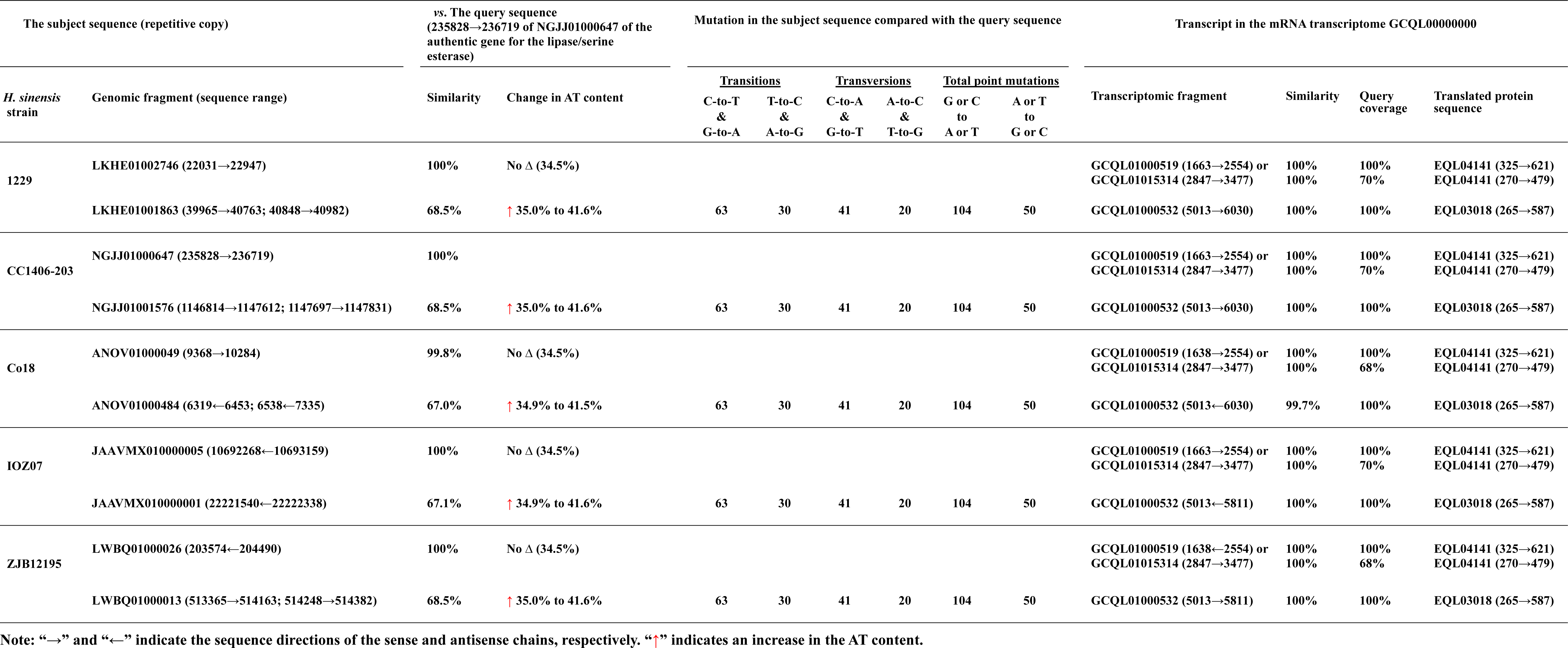
Authentic H. sinensis genes for the lipase/serine esterase (query sequence) and repetitive genomic copies with major increases (> 5%) in the AT content.

Table 7 shows the authentic genes for the maltose permease in 5 *H. sinensis* genome assemblies (47693→49623 within LKHE01002657; 77988→79917 within NGJJ01000203; 1725←3655 within ANOV01009266; 4614206→4616136 within JAAVMX010000006; and 66747→68677 within LWBQ01000157). The repetitive sequences (6824→7385 within LKHE01000336; 782871←782913 and 1076399←1076960 within NGJJ01001217; 4433←4994 and 5254←5290 within ANOV01000509; 12448464→12449025 and 12779815→12779864 within JAAVMX010000005; and 292175→292217 and 570642→571203 within LWBQ01000003; *cf*. Table 7) of the authentic gene showed 69.6–70.4% sequence similarity to the gene sequence. These repetitive sequences had more T-to-C and A-to-G transitions (35–37) than postulated RIP-related C-to-T and G-to-A transitions (30–31), as well as more C-to-A and G-to-T transversions (28) than A-to-C and T-to-G transversions (10–11). Overall, there were more G or C to A or T point mutations (58–59) than A or T to G or C point mutations (46–47) in the repetitive copies, resulting in slight increases (↑ ≤5%) in the AT content, from 36.8–37.9% in the authentic genes to 38.5–39.9% in the repetitive copies.

Table 7 also shows that the authentic genes (the genome segments within ANOV01009266, JAAVMX010000006, LKHE01002657, LWBQ01000157, and NGJJ01000203) were normally transcribed. The GCQL01010488 transcript was 99.8–100% homologous to the authentic genes for the maltose permease (accession #EQK98556 according to the GenBank protein annotation for the genome of the *H. sinensis* Strain Co18), with a query coverage of 80%. EQK98556 showed 74.6–84.9% similarity to the maltose permease or α-glucoside permease of 17 other microbes (*Drechmeria coniospora*, *Fusarium* sp., *Metarhizium* sp., *Ophiocordyceps* sp., *Tolypocladium* sp., *Trichoderma citrinoviride*).

The repetitive copies (the genome segments within ANOV01000509, JAAVMX010000005, LKHE01000336, LWBQ01000003, and NGJJ01001217; *cf*. Table 7) were transcribed, and 2 overlapping transcripts (GCQL01015773 and GCQL01019033) were 98.7% identical, with a query coverage of 100%. GCQL01015773 was 100% homologous to the repetitive copies, and GCQL01019033 was 95.6% similar to the repetitive copies, with a 25-nt deletion. These 2 transcripts encoded a different protein (MRT, a fungal- unique gene, encoding a raffinose family of oligosaccharide transporter; 152→338 of EQL02995, containing 550 amino acids according to the GenBank protein annotation for the genome of the *H. sinensis* Strain Co18); however, the protein encoded by GCQL01019033 had a deletion of 8 amino acids. EQL02995 showed 73.7–77.2% similarity to the MRT of 8 other microbes (*Metarhizium* sp., *Moelleriella libera*, *Ophiocordyceps* sp.).

In addition to the authentic *H. sinensis* genes for the maltose permease shown in Table 7, 16 other authentic genes were found, as outlined in Table 4, with slight increases in the AT content. Seven of them exhibited a mutagenetic pattern with contradictory transitions (T-to-C and A-to-G *vs*. RIP-related C-to-T and G-to-A) and transversions (T-to-G and A-to-C *vs*. C-to-A and G-to-T) in combination with various numbers of insertion and deletion point mutations, similar to the mutagenetic pattern for the maltose permease. Nine other genes had consistently more RIP-related C-to-T and G-to-A transitions than T-to-C and A-to-G transitions; and more C-to-A and G-to-T transversions than T-to-G and A-to-C transversions; and many insertion and deletion point mutations.

#### III-5.4. Large increases in the AT content of repetitive genomic copies

Table 8 shows the authentic genes for the lipase/serine esterase (22031→22947 within LKHE01002746; 235828→236719 within NGJJ01000647; 9368→10284 within ANOV01000049; 10692268←10693159 within JAAVMX010000005; and 203574←204490 within LWBQ01000026) in the 5 *H. sinensis* genomes. The repetitive sequences (39965→40763 and 40848→40982 within LKHE01001863; 1146814→1147612 and 1147697→1147831 within NGJJ01001576; 6319←6453 and 6538←7335 within ANOV01000484; 22221540←22222338 within JAAVMX010000001; and 513365→514163 and 514248→514382 within LWBQ01000013) were 67.0–68.5% similar to the authentic genes in the 5 *H. sinensis* genome assemblies. There were more postulated RIP-related C-to-T and G-to-A transitions (63) than T-to-C and A-to-G transitions (30); and more C-to-A and G-to-T transversions (41) than A-to-C and T-to-G transversions (20) in the repetitive copies. Overall, there were more G or C to A or T point mutations (104) in the repetitive sequences than A or T to G or C point mutations (50), resulting in large increases (↑ >5%) in the AT content, from 34.9–35.0% in the authentic genes to 41.5–41.6% in the repetitive copies.

Table 8 also shows that the highly homologous (99.8–100%) authentic genes (the segments within ANOV01000049, JAAVMX010000005, LKHE01002746, LWBQ01000026, and NGJJ01000647) were transcribed. Two transcripts (GCQL01000519 and GCQL01015314) were 100% homologous to the authentic gene and encoded partially overlapping protein sequences of lipase/serine esterase (accession #EQL04141, according to the protein annotation for the genome of the *H. sinensis* Strain Co18) with query coverages of 68–100%. EQL04141 showed 64.3% similarity to the putative serine esterase of *Hirsutella rhossiliensis* and 40.4–44.5% similarity to the lipase/serine esterase of 19 other microbes (*Beauveria bassiana*, *Dactylonectria estremocensis*, *Fusarium albosuccineum*, *Ilyonectria* sp., *Mariannaea* sp., *Metarhizium* sp., *Ophiocordyceps s*p., *Paramyrothecium foliicola*, *Pochonia chlamydosporia*, *Purpureocillium* sp., *Thelonectria olida*, *Tolypocladium* sp.).

The repetitive copies (the segments within ANOV01000484, JAAVMX010000001, LKHE01001863, LWBQ01000013, and NGJJ01001576; *cf*. Table 8) were 67.0–68.5% similar to the authentic gene and were normally transcribed. The transcript (GCQL01000532) was 99.7–100% homologous to the repetitive copies, with a query coverage of 100%; however, it encoded the hypothetical proteins OCS_01262 and G6O67_000668 (accession #EQL03018 and KAF4513393), according to the protein annotations for the genome of the *H. sinensis* Strains Co18 and IOZ07, respectively. EQL03018 and KAF4513393 were only 46.7% and 47.2% similar to EQL04141, encoded by the authentic gene, with a query coverage of 94%, and were 61.5–83.6% similar to many protein sequences for (1) **<u>the lipase/serine esterase</u>** of *Pochonia chlamydosporia* and *Purpureocillium* sp., (2) **<u>the serine esterase</u>** of *Hirsutella rhossiliensis*, *Ilyonectria robusta*, *Metarhizium acridum*, *Thelonectria olida*, and *Trichoderma breve*, and (3) **<u>the lipase-like proteins</u>**of *Fusarium* sp., *Hapsidospora chrysogenum*, *Metarhizium anisopliae*, *Tolypocladium* sp., and *Trichoderma lentiforme*. Thus, the protein sequence EQL04141 (encoded by the authentic *H. sinensis* gene) and the protein sequences EQL03018 and KAF4513393 (encoded by the repetitive copies) with approximately 47% similarities might represent lipid and ester hydrolases with different substrate specificities, *i.e.*, in theory hydrolyzing water-insoluble long chain triacylglycerols by lipases or water- soluble short acyl chain esters by esterases [59].

The repetitive copies of 10 other *H. sinensis* genes with largely increased AT content (*cf*. Table 4) consistently exhibited more RIP-related C-to-T and G-to-A transitions than T-to-C and A-to-G transitions, in addition to transversion, insertion, and deletion point mutations.

#### III-5.5. Bidirectional changes in the AT content of repetitive genomic copies

Table 9 shows the authentic gene for the β-lactamase/transpeptidase-like protein (19330→20932 within LKHE01000540; 70191→71142 within NGJJ01001580; 2857←4459 within ANOV01000528; 238583←240185 within JAAVMX010000002; and 168259←169259 within LWBQ01000052) among the 15 authentic genes that had multiple repetitive copies with bidirectional changes in the AT content in 5 *H. sinensis* genome assemblies. The multiple repetitive copies could be divided into 3 groups, 2 of which had reduced AT content but slightly different patterns of point mutations, and the other group had increased AT content.

**Table 9:**
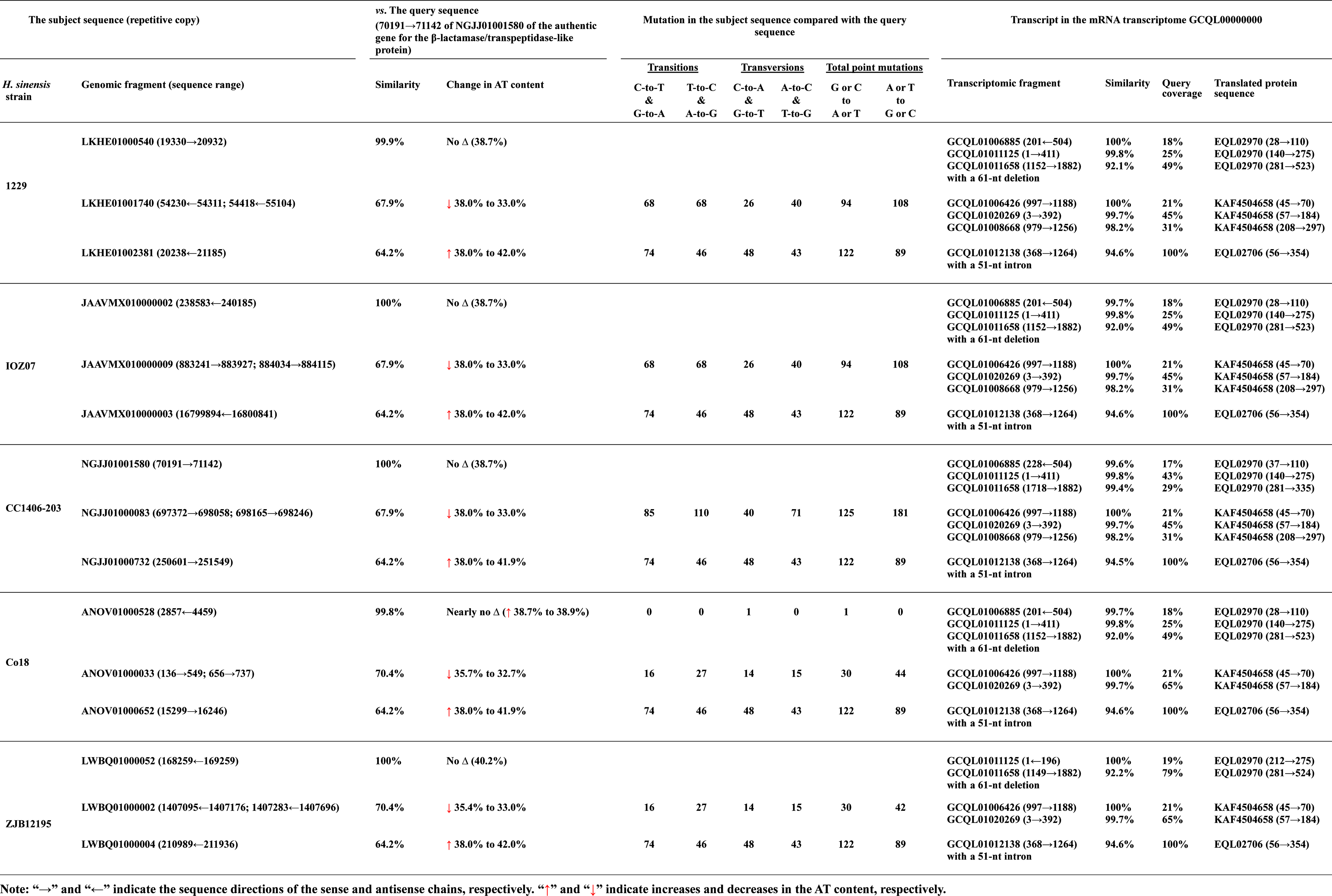
Authentic H. sinensis genes for the β-lactamase/transpeptidase-like protein (the query sequence) and repetitive genomic copies with bidirectional changes in the AT content.

The first group of repetitive copies with slight decreases (↓ ≤5%) in the AT content (54230←54311 and 54418←55104 within LKHE01001740 and 883241→883927 and 884034→884115 within JAAVMX010000009; *cf*. Table 9) had an equal number (68) of RIP-related C-to-T and G-to-A transitions and opposite T-to-C and A-to-G transitions, but more A-to-C and T-to-G transversions (40) than C-to-A and G-to-T transversions (26). Overall, the repetitive copies contained more A or T to G or C point mutations (108) than G or C to A or T point mutations (94), resulting in a sequence similarity of 67.9% to the authentic gene and a 5% decrease in the AT content from 38.0% in the authentic genes to 33.0% in the repetitive copies.

The second group of repetitive copies with slightly reduced AT content (697372→698058 and 698165→698246 within NGJJ01000083; 136→549 and 656→737 within ANOV01000033; and 1407095←1407176 and 1407283←1407696 within LWBQ01000002; *cf*. Table 9) showed 67.9–70.4% similarity to the authentic gene and had more T-to-C and A-to-G transitions (110 and 27) than postulated RIP-related C-to-T and G-to-A transitions (85 and 16) and more A-to-C and T-to-G transversions (71 and 15) than C-to-A and G-to-T transversions (40 and 14). Overall, there were fewer total G or C to A or T point mutations (125 and 30) in the repetitive copies than A or T to G or C point mutations (181 and 44), which was attributed to 2.4–5.0% decreases in the AT content, from 35.4–38.0% in the authentic gene sequences to 32.7–33.0% in the repetitive copies.

The third group of repetitive copies with increased AT content shown in Table 9 (20238←21185 within LKHE01002381; 250601→251549 within NGJJ01000732; 15299→16246 within ANOV01000652; 16799894←16800841 within JAAVMX010000003; and 210989←211936 within LWBQ01000004) exhibited 64.2% similarity to the authentic gene. There were more postulated RIP-related C-to-T and G-to- A transitions (74) than T-to-C and A-to-G transitions (46) and more C-to-A and G-to-T transversions (48) than A-to-C and T-to-G transversions (43) in the repetitive sequences. Overall, the repetitive copies had more G or C to A or T point mutations (122) than A or T to G or C point mutations (89), resulting in 3.9–4.0% increases in the AT content, from 38.0% in the authentic genes to 41.9–42.0% in the repetitive copies.

Additionally, as shown in Table 9, the authentic genes (the genome segments within ANOV01000528, JAAVMX010000002, LKHE01000540, LWBQ01000052, and NGJJ01001580) for the β-lactamase/transpeptidase-like protein were transcribed. Among the 3 fragmented transcripts, GCQL01006885 and GCQL01011125 were 99.6–100% homologous to the authentic gene, with query coverages of 17–43%, while the other transcript, GCQL01011658, was 92.1% similar (with a 58-nt deletion) to the authentic genes, with query coverages of 29–49%. The fragmented transcripts encoded the nonoverlapping sequences of the β-lactamase/transpeptidase-like protein with sequence gaps (accession #EQL02970, containing 531 amino acids), according to the protein annotation for the *H. sinensis* Strain Co18.

The repetitive copies with reduced AT content (the genome segments within ANOV01000033, JAAVMX010000009, LKHE01001740, LWBQ01000002, and NGJJ01000083; *cf*. Table 9) were transcribed. The transcripts (GCQL01006426, GCQL01008668, and GCQL01020269) were 98.2–100% homologous to the sequences of the repetitive copies, with query coverages of 21–65% and short segment overlaps and gaps in protein sequences. The transcripts presumably encoded the hypothetical protein G6O67_008083 (accession #KAF4504658, containing 542 amino acids), according to the protein annotation for the *H. sinensis* Strain IOZ07. KAF4504658 was 49.1–80.4% similar to 24 GenBank protein sequences for the β-lactamase/transpeptidase-like protein of many different microbes (*Dactylonectria* sp., *Fusarium* sp., *Hirsutella* sp., *Ilyonectria* sp., *Mariannaea* sp., *Microthyrium microscopicum*, *Ophiocordyceps* sp., *Thelonectria* olida, *Trichoderma* sp.).

The repetitive copies with increased AT content (the genome segments within ANOV01000652, JAAVMX010000003, LKHE01002381, LWBQ01000004, and NGJJ01000732; *cf*. Table 9) were transcribed. The GCQL01012138 transcript was 94.6% similar (with a 51-nt deletion) to the repetitive sequences and encoded a β-lactamase/transpeptidase-like protein (accession #EQL02706, containing 558 amino acids, according to the protein annotation for the *H. sinensis* Strain Co18), with 100% query coverage. The protein sequences of EQL02706 and KAF4504658, which were encoded by the repetitive sequences, were 49.3% similar to each other, however, were only 50.0% and 46.2% similar, respectively, to that of EQL02970, which is encoded by the authentic gene for the β-lactamase/transpeptidase-like protein. The sequences of EQL02706 and KAF4504658 were (1) 47.9–80.4% similar to those of **<u>the β-lactamase/transpeptidase-like proteins</u>** of various microbes (*Akanthomyces lecanii*, *Dactylonectria* sp., *Fusarium* sp., *Hirsutella rhossiliensis*, *Ilyonectria* sp., *Mariannaea* sp., *Metarhizium* sp., *Microthyrium microscopicum*, *Thelonectria* sp., and *Trichoderma* sp.) and (2) 49.3–63.3% similar to those of **<u>the penicillin-binding proteins</u>** of various microbes (*Drechmeria coniospora*, *Fusarium* sp., *Pochonia chlamydosporia*, *Purpureocillium* sp., and *Trichoderma arundinaceum*).

### III-6. Transcriptional silencing of 5.8S genes

The genetic, genomic, and phylogenetic analyses shown in Tables 1‒3 and Figures 1‒3 demonstrated GC-biased characteristics of the multiple repetitive ITS copies in the *H*. *sinensis* genome assemblies and that all GC- and AT-biased genotypes of *O*. *sinensis* were genome independent. Li *et al*. [1] identified ITS sequences for both Genotypes #1 and #5 from genomic DNA that was isolated from 8 (1206, 1208, 1209, 1214, 1220, 1224, 1227, and 1228) of 15 clones of natural *C. sinensis* multicellular heterokaryotic mono- ascospores [45]. However, Li *et al*. [1] found 5.8S cDNA only for Genotype #1 and not for Genotype #5 in a cDNA library constructed from total RNA extracted from Clone 1220, one of the 8 genetically heterogeneous clones. Li *et al*. [1] did not report cDNA library construction for 7 other genetically heterogeneous clones (1206, 1208, 1209, 1214, 1224, 1227, and 1228) or subsequent cDNA examinations for Genotypes #1 and #5 of *O. sinensis*.

Li *et al*. [1] did not detect the ITS sequences of AT-biased Genotypes #4, #6, and #15‒17 in the genomic DNA isolated from any of the 15 ascosporic clones. Thus, there is insufficient evidence to conclude that the 5.8S genes of all AT-biased genotypes of *O. sinensis* are permanently nonfunctional pseudogenic components coexisting in “a single genome” with a functional copy of GC-biased Genotype #1. To respond to the academic challenge [25], Li *et al*. [60] provided the following logical reasoning underlying the nonfunctional pseudogene hypothesis: “If AT-biased ITS sequences are not pseudogenes, 5.8S cDNA should have been detected because rRNAs are critical and essential for protein synthesis in all living organisms”; this logical reasoning critically touches on 2 serious scientific issues. First, are the 5.8S genes of all the *O. sinensis* genotypes actively transcribed in *C. sinensis* ascospores (discussed in this section)? Second, is it correct to conclude that “5.8S cDNA should have been detected” by the techniques designed by Li *et al*. [1] (discussed in Section III-7)?

Many research groups have reported the differential coexistence of several AT-biased genotypes in various combinations in different compartments of natural *C. sinensis*, as summarized in Table 10 [5–9,21,24,26–28,32,42]. Genotypes #4 and #15 of the AT-biased Cluster B of *O. sinensis* (*cf*. Figure 2) were reported to exist in the stroma and stromal fertile portion (densely covered with numerous ascocarps) of natural *C. sinensis* but not in the ascospores [5–9,32], consistent with the failure to identify Genotype #4 of AT-biased Cluster B in mono-ascospores as reported by Li *et al*. [1] after additional specific efforts. Thus, there is no evidence demonstrating that AT-biased Genotypes #4 and #15 occur in *C. sinensis* ascospores, and there is no ground for claiming that the 5.8S genes of Genotypes #4 and #15 are nonfunctional pseudogenes in the genome of GC-biased Genotype #1.

**Table 10.** Differential occurrence of AT-biased genotypes of O. sinensis in different compartments of natural C. sinensis.

| Genotype # and GenBank accession # | Phylogenetic AT-biased Cluster § | Stroma |  | Stromal Fertile Portion (densely covered with numerous ascocarps) | Ascospores |
| --- | --- | --- | --- | --- | --- |
|  |  | Immature | Mature |  |  |
| #4 AB067744 | B | √ | √ | √ |  |
| #5 AB067740 | A | √ | √ | √ | √ |
| #6 EU555436 | A | √ | √ | √ | √ |
| #15 KT232017 | B | √ | √ | √ |  |
| #16 KT232019 | A | √ |  |  | √ |
| #17 KT232010 | A | √ |  |  |  |

Note: This table was adapted and altered from Li *et al.* [9]. §, Phylogenetic AT-biased Clusters A and B are shown in Figure 2 (the Bayesian tree) and are similarly shown in the phylogenetic trees that were inferred using various phylogenetic algorithms by Li *et al.* [7–9,31–32,42].

Xiang *et al*. [55] reported a metatranscriptomic study of natural *C. sinensis* specimens (unknown maturation stage) collected from Kangding County in Sichuan Province, China. A GenBank BLAST search confirmed that the metatranscriptome GAGW00000000 did not contain any 5.8S gene transcripts of *C. sinensis*-associated fungi (including all genotypes of *O. sinensis*), indicating transcriptional silencing of the 5.8S genes of all cocolonizing fungi in natural settings.

Xia *et al*. [56] reported another metatranscriptomic study of mature natural *C. sinensis* specimens collected from Deqin County in Yunnan Province, China. The metatranscriptomic assembly was uploaded to www.plantkingdomgdb.com/Ophiocordyceps_sinensis/data/cds/Ophiocordyceps_sinensis_CDS.fas. Similarly, this assembled metatranscriptome contained no 5.8S gene transcripts. However, the authors uploaded the unassembled shotgun reads (SRR5428527) to GenBank. A BLAST search of the SRR5428527 Shotgun Read Archive (SRA) using the MegaBLAST algorithm identified 1894 unassembled metatranscriptomic shotgun reads (100 bp each) that were 100% homologous to the 5.8S gene sequence (156 bp) of GC-biased Genotype #1 *H. sinensis*. Figure 5 shows the GenBank BLAST graphic summary of the alignment of the 5.8S gene (the query sequence; segment 218‒373 of Genotype #1 *H. sinensis* AB067721) and the first 75 of the overlapping 1894 metatranscriptomic shotgun reads (100 bp each) of mature natural *C. sinensis*.

In general, fungal genomes contain multiple repetitive copies of rDNA ITS1-5.8S-ITS2 sequences (*cf*. Tables 1 and 3; Figures 1‒3). Because the assembled metatranscriptome contained no 5.8S gene transcripts [55–56] and no segments of the rDNA 18S or 28S gene of *H. sinensis* were homologous to the 5.8S gene, the 1894 unassembled shotgun reads (SRR5428527) of natural *C. sinensis* may have been derived from highly homologous non-rRNA genes that evolved in a conservative manner within fungal metagenomes.

Various degrees of sequence similarity were found when the 1894 unassembled metatranscriptomic shotgun reads were aligned with the 5.8S gene sequences of Genotypes #2–17 of *O. sinensis*. Table 11 shows 6 examples of shotgun reads of SRX2718557 aligned with the 5.8S gene sequences of multiple *O. sinensis* genotypes, *P. hepiali*, *T. sinensis* (Group D), and an AB067719-type Group-E fungus [18,41]. The high homology between the shotgun reads and the various 5.8S gene sequences of *O. sinensis* genotypes may not conclusively define the fungal sources in natural *C. sinensis*, which contains >90 species spanning at least 37 fungal genera and 17 genotypes of *O. sinensis* [5–32,55–56].

**Table 11.** Similarities between the first 6 of 1894 unassembled metatranscriptomic shotgun reads (SRA: SRR5428527) of natural C. sinensis and the 5.8S genes of 17 genotypes of O. sinensis and P. hepiali, T. sinensis (Group D) and an AB067719-type (Group E) fungus.

| Geno-<br>type | GenBank<br>accession # | Percent similarity between the *C. sinensis* metatranscriptomic shotgun reads and the 5.8S gene<br>sequences of multiple *O. sinensis* genotypes and some *C. sinensis*-colonizing fungi |  |  |  |  |  |
| --- | --- | --- | --- | --- | --- | --- | --- |
|  |  | SRR5428527.1 | SRR5428527.1 | SRR5428527.1 | SRR5428527.1 | SRR5428527.1 | SRR5428527.179 |
|  |  | 7546327.1 | 7554054.1 | 7565575.2 | 7679995.2 | 7911498.2 | 29569.2 |
| #1 | AB067721 | <u>100%</u> | <u>100%</u> | <u>100%</u> | <u>100%</u> | <u>100%</u> | <u>100%</u> |
| #2 | MG770309 | 96% | 96% | <u>100%</u> | 96% | <u>98%</u> | 96% |
| #3 | HM595984 | <u>100%</u> | <u>99%</u> | <u>99%</u> | <u>100%</u> | <u>99%</u> | <u>100%</u> |
| #7 | AJ488254 | <u>98%</u> | <u>98%</u> | <u>99%</u> | <u>98%</u> | <u>98%</u> | <u>98%</u> |
| #8 | GU246286 | 94% | 96% | 95% | 94% | <u>97%</u> | 94% |
| #9 | GU246288 | <u>99%</u> | <u>99%</u> | <u>99%</u> | <u>100%</u> | <u>99%</u> | <u>100%</u> |
| #10 | GU246287 | 96% | 94% | 93% | 95% | 93% | <u>97%</u> |
| #11 | JQ695935 | <u>100%</u> | <u>100%</u> | <u>100%</u> | <u>100%</u> | <u>100%</u> | <u>100%</u> |
| #12 | GU246296 | <u>99%</u> | <u>99%</u> | <u>99%</u> | <u>99%</u> | <u>99%</u> | <u>99%</u> |
| #13 | KT339190 | <u>98%</u> | 93% | 93% | <u>98%</u> | 93% | <u>98%</u> |
| #14 | KT339178 | <u>98%</u> | 93% | 93% | <u>98%</u> | 93% | <u>98%</u> |
| #4 | AB067744 | 86% | 87% | 87% | 83% | 87% | 83% |
| #5 | AB067740 | 85% | 87% | 87% | 84% | 87% | 84% |
| #6 | KJ720572 | 85% | 85% | 87% | 84% | 86% | 84% |
| #15 | KT232017 | 85% | 88% | 88% | 84% | 88% | 84% |
| #16 | KT232019 | 86% | 86% | 87% | 85% | 87% | 85% |
| #17 | KT232010 | 87% | 88% | 88% | 87% | 88% | 87% |
| P.h. | EF555097 | <u>99%</u> | <u>98%</u> | <u>97%</u> | <u>99%</u> | <u>98%</u> | <u>99%</u> |
| T.s. | AB067720 | <u>98%</u> | <u>97%</u> | 96% | <u>98%</u> | <u>97%</u> | <u>98%</u> |
|  | AB067719 | <u>98%</u> | 93% | 93% | <u>98%</u> | 93% | <u>98%</u> |

Note: The GenBank sequence ID SRA: SRR5428527 fragments (17546327.1; 17554054.1; 17565575.2; 17679995.2; 17911498.2; and 17929569.2) are the unassembled transcriptome shotgun sequences obtained from the mature natural *C.* *sinensis* specimens collected from Deqin County in Yunnan Province by Xia et al. [56]. GC-biased genotypes are listed in blue, and AT-biased genotypes are listed in red. P.h. refers to *Paecilomyces hepiali*; T.s. refers to Group-D *Tolypocladium sinensis*; and AB067719 refers to an AB067719-type Group-E fungus [18,41]. Highly homologous percent similarities (97–100%) are underlined.

As shown in Table 11, the representative unassembled metatranscriptomic shotgun reads of natural *C. sinensis* were 97‒99% homologous to the 5.8S gene sequence of *P. hepiali*. Some of the shotgun reads were 97‒98% homologous to the 5.8S gene sequences of *T. sinensis* (Group D) and the AB067719-type Group- E fungus [18,41]. A GenBank BLAST search revealed the alignment of the 5.8S gene sequence (segment 218→373 within AB067721) of Genotype #1 *H. sinensis* to 56‒438 unassembled transcriptome shotgun sequence reads, which were identified in 5 data files of RNA-Seq experiments (GenBank assessment #SRX3083435‒SRX3083439) of *P. hepiali* Strain Feng (SRA Study: SRP115168 [61]) (Table 12). Pang *et al*. [61] uploaded 5 transcriptomic experimental datasets to GenBank for the total RNA samples extracted from cultures of *P. hepiali* Strain Feng under different culture conditions and durations.

**Table 12.** Summary of BLAST searches of the transcriptome shotgun reads of P. hepiali Strain Feng vs. the 5.8S gene sequence (218→373 within AB067721) of Genotype #1 H. sinensis.

| *P. hepiali* sample | Culture duration (days) | Additive in culture medium | GenBank transcriptome data file | vs. the 5.8S gene sequence of *H. sinensis* |  |  |
| --- | --- | --- | --- | --- | --- | --- |
|  |  |  |  | # of the total BLAST hits | (% of total filtered reads) | % Similarity |
| P4 | 4 | no | SRX3083438 | 219 | (3.7% x 10<sup>-4</sup>) | 93–99% |
| P9 | 9 | no | SRX3083437 | 56 | (3.6% x 10<sup>-4</sup>) | 95–100% |
| P15 | 15 | no | SRX3083436 | 438 | (6.12% x 10<sup>-4</sup>) | 96–100% |
| Ps | 4 | Adenosine | SRX3083435 | 356 | (4.13% x 10<sup>-4</sup>) | 96–100% |
| Pa | 4 | Adenine | SRX3083439 | 207 | (6.12% x 10<sup>-4</sup>) | 96–99% |

Note: Data are adapted from Pang *et al.* 2018 http://dx.doi.org/10.1016/j.ygeno.2017.09.008.

The extremely high sequence similarities among the 5.8S gene sequences of *H. sinensis*, *P. hepiali* and *T. sinensis,* as shown in Table 11, indicate conservative evolution of the 5.8S gene in many fungal species. Table 12 shows significant differences in the number of BLAST hits and percent similarities of the 5.8S gene sequence of *H. sinensis* aligned with the transcriptomic shotgun reads obtained from culture experiments of *P. hepiali* Strain Feng under different culture conditions and durations [61]. The results indicate the experimental switching on or off of *P. hepiali* genes, including the 5.8S gene and many other non-rRNA genes, in response to the different experimental settings.

Liu *et al*. [54] reported the mRNA transcriptome assembly GCQL00000000, which was derived from the *H. sinensis* Strain L0106 and contained no rRNA or tRNA transcripts. Table 13 summarizes the dynamic, nonlinear alterations during continuous *in vitro* liquid fermentation in the total number of transcriptomic unigenes, average length and GC content of the transcripts of the *H. sinensis* Strain L0106, indicating the nonnatural differential transcriptional activation (switching on) and silencing (switching off) of numerous *H. sinensis* genes in response to continued liquid fermentation of the *H. sinensis* Strain L0106. Table 13 suggests that more dramatic changes might conceivably occur after 9 days of fermentation. The prolonged 25-day liquid incubation period adopted by Li *et al*. [1] might have a greater impact on the unbalanced transcriptional activation and silencing of long and short fungal genes with various GC content biases, in addition to the changes in the culture medium as an important factor influencing the transcriptional activation and silencing of many fungal genes (*cf*. Table 12).

**Table 13.**
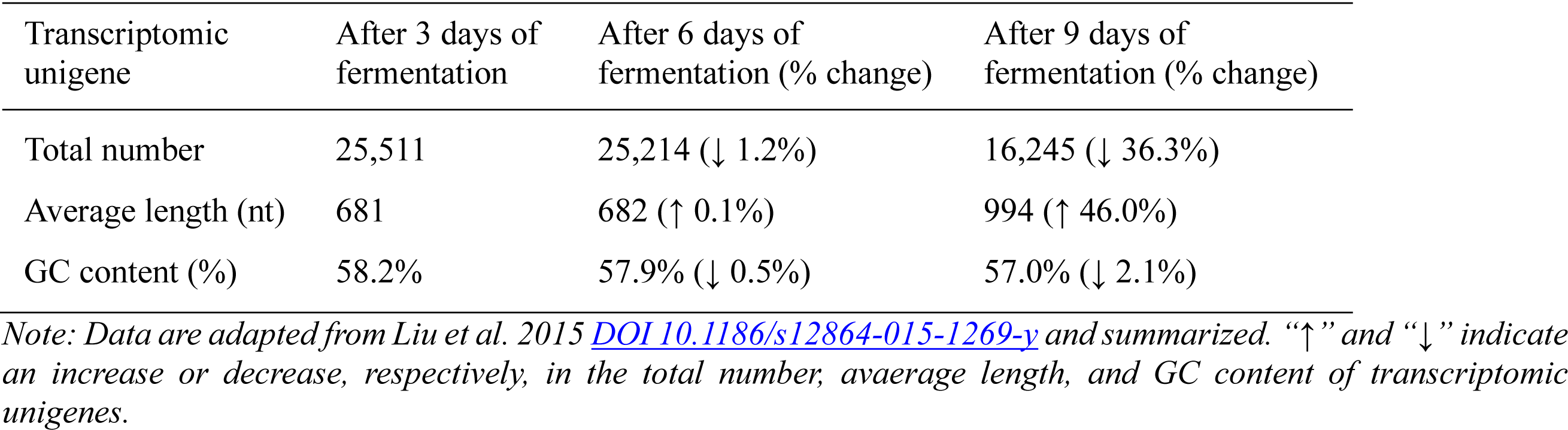
Dynamic alterations of transcripts in the mRNA transcriptome of the H. sinensis Strain L0106 during continuous in vitro fermentation.

| Transcriptomic unigene | After 3 days of fermentation | After 6 days of fermentation (% change) | After 9 days of fermentation (% change) |
| --- | --- | --- | --- |
| Total number | 25,511 | 25,214 (↓ 1.2%) | 16,245 (↓ 36.3%) |
| Average length (nt) | 681 | 682 (↑ 0.1%) | 994 (↑ 46.0%) |
| GC content (%) | 58.2% | 57.9% (↓ 0.5%) | 57.0% (↓ 2.1%) |

*Note: Data are adapted from Liu et al. 2015 [DOI 10.1186/s12864-015-1269-y](https://doi.org/10.1186/s12864-015-1269-y) and summarized. “↑” and “↓” indicate* *an increase or decrease, respectively, in the total number, average length, and GC content of transcriptomic* *unigenes.*

### III-7. Detection of 5.8S cDNA using 5.8S-F/R primers

In addition to the transcriptional silencing of 5.8S genes in natural settings and the significant impact of continuous *in vitro* culture and different experimental settings on unbalanced gene transcription, as discussed above, the second issue related to the logic described by Li *et al*. [60] is whether the methods designed by Li *et al*. [1] were suitable for scientifically identifying 5.8S gene cDNAs of multiple *O. sinensis* genotypes [34].

For instance, Li *et al*. [60] retrospectively explained the research design logic for the 5.8S-F/R primer pair that was used by Li *et al*. [1]: “The 5.8S-F/R primers … located in the most conserved region.” Accordingly, there would be no doubt that such a design would greatly diminish the specificity of the primers for 5.8S cDNA because of the conservative evolution of the 5.8S gene and many other fungal genes. Table 14 and Figure 6 show the similarities and alignments between the sequences of the primers 5.8S-F (5’ ACTTTTAACAACGGATCTCTT 3’) and 5.8S-R (5’ AAGATAACGCTCGGATAAGCAT 3’) and the 5.8S genes of Genotypes #1, #4, and #5.

**Table 14.**
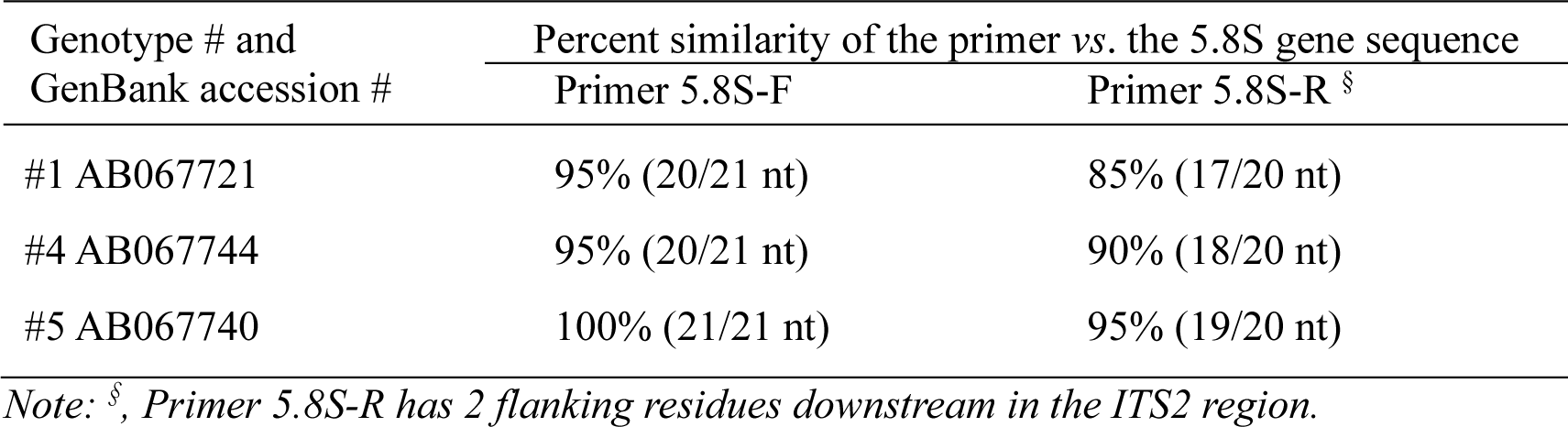
Similarity of the 5.8S-F/R primers and the 5.8S gene sequences of Genotypes #1, #4 and #5 of O. sinensis.

| Genotype # and GenBank accession # | Percent similarity of the primer vs. the 5.8S gene sequence |  |
| --- | --- | --- |
|  | Primer 5.8S-F | Primer 5.8S-R § |
| #1 AB067721 | 95% (20/21 nt) | 85% (17/20 nt) |
| #4 AB067744 | 95% (20/21 nt) | 90% (18/20 nt) |
| #5 AB067740 | 100% (21/21 nt) | 95% (19/20 nt) |

*Note: §, Primer 5.8S-R has 2 flanking residues downstream in the ITS2 region.*

The primer pair 5.8S-F/R was 100% and 95% homologous to the 5.8S gene sequence of AT-biased Genotype #5, as shown in Table 14. In contrast to the 95% and 85% similarity to the 5.8S gene sequence of GC-biased Genotype #1, the 5.8S-F/R primer pair may be favored for amplifying the 5.8S cDNA sequence of Genotype #5 rather than for amplifying the 5.8S cDNA sequence of Genotype #1. The favorability of the primers for the 5.8S cDNA sequence of Genotype #5 is clearly shown in Figure 6. The 5.8S-F/R primers showed variable sequence similarities, ranging from 61.9‒100% and 31.8‒95.5%, respectively, *vs*. the 17 genotypes of *O. sinensis*, preferentially amplifying the most easily amplified cDNA sequence in the cDNA library considering the primary structure and secondary conformation of the PCR templates and consequently causing false-negative results for the other 5.8S cDNA sequences.

Section III-6 describes the 100% homology between the 1894 unassembled metatranscriptomic shotgun reads (100 bp) of mature natural *C. sinensis* and the 5.8S gene sequence (156 bp) of GC-biased Genotype #1 *H. sinensis*, including many non-rRNA transcripts among the highly homologous 1894 shotgun reads in natural settings. A BLAST search of *O. sinensis* transcriptomic sequences (taxid: 72228) in the GenBank database showed that the 5.8S-F/R primer pair was 100% homologous to the mRNA transcriptome assemblies GCQL01001191, GCQL01006777, GCQL01007818, and GCQL01013443 of the Genotype #1 *H. sinensis* Strain L0106 [54], which do not contain rRNA or tRNA transcripts, and to the assembled metatranscriptome sequences GAGW01003691, GAGW01007670, and GAGW01007671 of natural *C. sinensis* [55]. Figure 7 shows that the metatranscriptome GAGW01003691 is 99.8% homologous to a segment of the mRNA transcriptome GCQL01006777 of the *H. sinensis* Strain L0106; the metatranscriptomes GAGW01007670 and GAGW01007671 are 100% identical to the mRNA transcriptome segments of GCQL01007818 and GCQL01001191, respectively [34].

**Figure 7.**
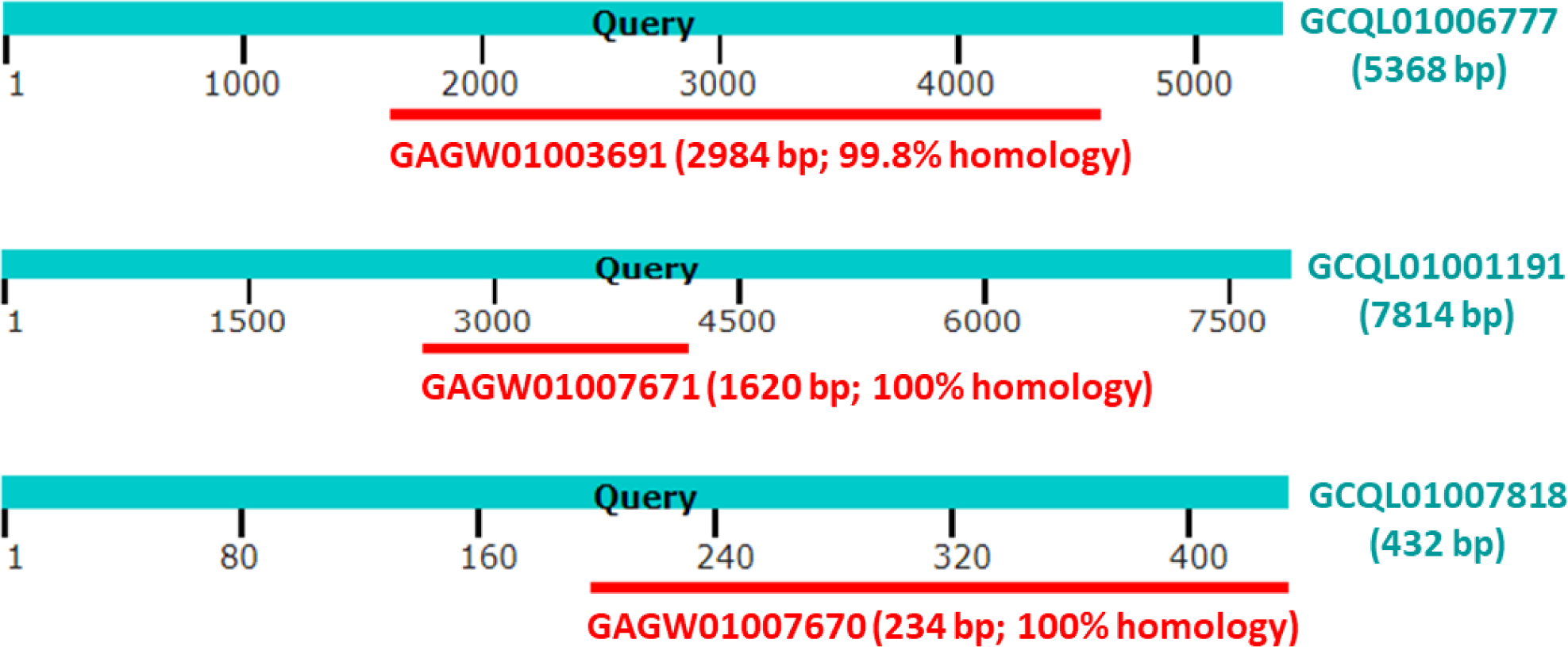
Graphic alignment summary of the mRNA transcriptome assembly sequence (GCQL00000000) of the H. sinensis Strain L0106 and the metatranscriptome assembly sequence (GAGW00000000) of natural C. sinensis. The graphic alignment summary was generated using GenBank BLAST. GCQL01001191, GCQL01006777, and GCQL01007818 are sequences of transcriptome segments that were obtained from the mRNA pool of the H. sinensis Strain L0106 and contained no rRNA or tRNA transcripts. GAGW01003691, GAGW01007670, and GAGW01007671 are segment metatranscriptome sequences obtained from the total RNA of natural C. sinensis specimens collected from Kangding County of Sichuan Province, China.

Because the mRNA transcriptome assembly GCQL00000000 does not contain rRNA or tRNA transcripts, the extremely high homology between the segments of the metatranscriptome GAGW00000000 and transcriptome sequence GCQL00000000 indicates that the 5.8S-F/R primer pair designed by Li *et al*. [1] may be capable of recognizing and nonspecifically amplifying non-rRNA components in the cDNA library derived from the genetically heterogeneous Clone 1220, which is one of the 8 heterogeneous clones (1206, 1208, 1209, 1214, 1220, 1224, 1227, and 1228) of *C. sinensis* mono-ascospore cultures [1].

Unfortunately, Li *et al*. [1] did not disclose the sequences of the identified 5.8S cDNA but instead reported that “Of these clones, 28 showed identical sequences (104 bp excluding the primers) to the most frequent sequence in group A (JQ900148). The remaining 14 sequences … had single substitutions at various positions”, where group A = Genotype #1 *H. sinensis*. The GenBank annotation of the 5.8S gene of JQ900148 is at 192→346, covering 155 nt, which is not a multiple of three and is not consistent with the open reading frame of the gene.

Alternatively, the 5.8S genes were annotated at the following nucleotide positions: 218→373 within AB067721 of Genotype #1 and AB067740 of Genotype #5, as well as 214→369 within AB067744 of Genotype #4, covering 156 nt and 52 codons in the 3 cases [6,31,34]. Excluding the 5.8S-F/R primer regions (*cf*. Figure 6), the remaining sequence of the 5.8S gene consisted of 114 bp, with 10 additional nucleotides (mismatches) beyond the 104 bp in the 28 cDNA clones reported by Li *et al*. [1]. Upon adding “single substitutions at various positions” to 14 other clones, as reported by Li *et al*. [1], the 42 clones presumably contained either 10 or 11 mismatches (8.8‒9.6%) in the 114-bp segment of the 5.8S gene cDNA, corresponding to 90.4‒91.2% sequence similarity to the 5.8S gene of *H. sinensis*. Such low similarities indicated the unlikelihood of the 5.8S gene cDNA detected by Li *et al*. [1] truly belonging to GC-biased Genotype #1. Alternatively, nondisclosure of the detected cDNA sequence did not exclude the possibility that the detected cDNA [1] was derived from the AT-biased Genotype #5 of *O. sinensis*.

## IV. Discussion

### IV-1. The GC-biased repetitive ITS copies in the genome of Genotype #1 *H. sinensis* are unrelated to RIP mutagenesis

Selker and his colleague [62–65] reported frequent C-to-T (cytosine-to-thymine) transitions and concomitant epigenetic methylation of nearly all remaining cytosine residues in fungal 5S RNA genes associated with transcriptional silencing. Gladyshev [66] suggested that RIP “occurs specifically in haploid parental nuclei that continue to divide by mitosis in preparation for karyogamy and ensuing meiosis” and “mutates cytosines on both strands of each DNA duplex in a pairwise fashion (e.g., preferentially affecting none or both repeated sequences).” Hane *et al*. [67] stated that “the RIP process selectively mutated duplicated sequences in both DNA strands by inducing single-nucleotide point (SNP) mutations that converted C:G base pairs to T:A. This often led to the introduction of nonsense or missense mutations which affected the expression of these sequences.” The sequence-specific targets of RIP mutagenesis for *de novo* DNA methylation are under debate [67–69].

Table 3 and Figure 1 of this study demonstrate that the variable repetitive ITS copies were genetically characterized by multiple scattered insertion, deletion and transversion point mutations. These mutations were not generated through RIP mutagenesis, which theoretically causes cytosine-to-thymine (C-to-T) and guanine-to-adenine (G-to-A) transitions. On the other hand, the AT-biased genotypes of *O. sinensis* characterized by multiple scattered transition point mutations (*cf*. Figure 1) were generated in parallel with other GC-biased genotypes through accelerated nuclear rDNA evolution from a common ancestor over the long course of evolution [18], regardless of whether this occurred *via* RIP mutagenesis or other evolutionary mechanisms. Thus, evidence is lacking to support the hypothesis that RIP mutagenesis induces the formation of mutant AT-biased repetitive ITS copies within the single genome of GC-biased Genotype #1 *H. sinensis*, as postulated by Li *et al*. [47]. Even if AT-biased genotypes were indeed induced by RIP mutagenesis over the long course of evolution, there has been no evidence that AT-biased genotypes “could have emerged either before or after” a new GC-biased “ITS haplotype was generated” through “repeat-induced point mutations” (RIP) and became pseudogenic components of the single genome of GC-biased Genotype #1, as assumed by Li *et al*. [1,47,60].

Taranto *et al*. [69] and Hane *et al*. [67] reported that RIP “is observed only in certain fungal taxa; the Pezizomycotina (filamentous Ascomycota) and some species of the Basidiomycota.” Tables 1 and 3 and Figures 1‒3 show that all the multiple genomic repetitive ITS copies were GC biased genetically, with no or only a few transition alleles, and were phylogenetically distinct from AT-biased genotypes, which possess numerous scattered transition alleles. It is thus reasonable to question whether Genotype #1 *H. sinensis* is susceptible to RIP mutagenesis and epigenetic methylation attack.

### IV-2. Genetic and phylogenetic differences in the AT-biased genotypes of *O. sinensis* and GC-biased repetitive ITS copies in the genome of Genotype #1 *H. sinensis*

The sequences of AT-biased Genotypes #4‒6 and #15‒17 of *O. sinensis* shown in red in Figure 1 contain numerous scattered transition alleles. These sequences are genetically distinct and phylogenetically distant from GC-biased repetitive ITS copies containing the predominant insertion, deletion, and transversion point mutations within the genome of GC-biased Genotype #1 *H. sinensis* (*cf*. Figures 1‒2 and Tables 1‒3). The sequences of AT-biased Genotypes #4‒6 and #15‒17 of *O. sinensis*, as well as GC-biased Genotypes #2‒3 and #7‒14, are absent in the genome assemblies ANOV00000000, JAAVMX000000000, LKHE00000000, LWBQ00000000, and NGJJ00000000 of the *H. sinensis* Strains Co18, IOZ07, 1229, ZJB12195, and CC1406-203, respectively (*cf*. Tables 1‒2 and Figures 1 and 3), but instead belong to the genomes of independent *O. sinensis* fungi [5–9,21,25,31,34,41,44,50–53]. The nonresidency of AT-biased *O. sinensis* genotypes does not support the hypothesis that the AT-biased sequences are “ITS pseudogene” components of “a single genome” of GC-biased Genotype #1 *H. sinensis*. Stensrud *et al*. [18] concluded that the variation in the 5.8S gene sequences of AT-biased genotypes “far exceeds what is normally observed in fungi … even at higher taxonomic levels (genera and family)”. Xiao *et al*. [21] concluded that the sequences of AT-biased genotypes likely belong to the genomes of independent *O. sinensis* fungi.

### IV-3. Previously hypothesized centers of origin and evolutionary geography for *O. sinensis*

GC-biased genotypes might have evolved at accelerated rates in parallel with AT-biased genotypes from an ancient common ancestor during long-term evolution, as hypothesized by Stensrud *et al*. [18]. Zhang *et al*. [22] detected GC-biased Genotype #3 in natural *C. sinensis* specimens collected from the Nyingchi District of Tibet and hypothesized that “Nyingchi might have acted as a center of origin of *O. sinensis* and *O. sinensis* might have been transmitted from Nyingchi to other locations”. Accordingly, GC-biased Genotype #3 might represent a modern “immediate” ancestor of Genotype #1 *H. sinensis* that has been consistently identified in other production locations as the result of a relatively modern phase of long-term evolution. The authors then described an evolutionary path from GC-biased Genotype #3 to Genotype #1 and the transmission of *O. sinensis* from the “center of origin … spread eastward and westward first among southern regions on the Tibetan Plateau” and then “spread from south to north … at a later time”.

Unfortunately, this evolutionary geography for GC-biased *O. sinensis* lacks an evolutionary branch or branches related to AT-biased genotypes of *O. sinensis*. Perhaps the generation of AT-biased genotypes of *O. sinensis* might have occurred during more ancient historical evolutionary phases. Undoubtedly, the discovery of AT-biased genotypes of *O. sinensis* will trigger further research on the history of *O. sinensis* evolution and fungus-insect coevolution [70].

The evolutionary geography hypothesis for *O. Sinensis* also overlooks an evolutionary path or paths extending south to (or north of) the production areas in Bhutan, Sikkim. Nepal. Biswa *et al*. [71] identified a mutant sequence (MW990119) from a *C. sinensis* specimen (CBUS3) collected in Yumesamdong (27°84’88” N 88°69’04” E), North Sikkim, which is far from Nyingchi in Tibet (29°38’56” N, 94 21’41” E). MW990119 was 98.8% homologous to HM595984 of the representative sequence of GC-biased Genotype #3, which had 3 insertions and 2 transition alleles but was 95.2% similar to AB067721 of Genotype #1 *H. sinensis*. The discovery of the MW990119 sequence may challenge the “center of origin” hypothesis for Nyingchi, Tibet, and the “evolutionary geography” hypothesis for *O. sinensis*, which may need to be supplemented, amended, or reassessed.

### IV-4. Multicellular heterokaryons of natural *C. sinensis* hyphae and ascospores

Bushley *et al*. [45] illustrated that *C. sinensis* hyphae and ascospores were multicellular heterokaryons with mononucleated, binucleated, trinucleated, and tetranucleated cellular structures. Zhang & Zhang [46] asked whether the binucleated cells (and other monokaryotic and polykaryotic cells) of *C. sinensis* hyphae and ascospores contained homogenous or heterogeneous genetic materials, suggesting that these monokaryotic and polykaryotic cells may contain heterogeneous sets of chromosomes and genomes.

Li *et al*. [1] identified heterogeneous ITS sequences of both GC-biased Genotype #1 and AT-biased Genotype #5 in 8 of 15 ascosporic clones (1206, 1208, 1209, 1214, 1220, 1224, 1227, and 1228) and simultaneously identified 7 other clones (1207, 1218, 1219, 1221, 1222, 1225, and 1229) that contained only homogenous sequences of Genotype #1 after 25 days of *in vitro* inoculation with natural *C. sinensis* mono-ascospores. These culture-dependent molecular mycological findings support the morphological observations of multicellular heterokaryons discovered by their collaborators [45], although molecular mycological strategies may overlook nonculturable fungal components. Apparently, the 2 groups of reported ascosporic clones were derived from different cells of multicellular heterokaryotic mono-ascospores of natural *C. sinensis*, confirming the inference of Zhang & Zhang [46] that there are heterogeneous sets of chromosomes and genomes in mono- and polykaryotic cells.

In addition to the 15 clones reported by Li *et al*. [1], the authors did not disclose any information on other possible ascosporic clones, namely, clones 1210, 1211, 1212, 1213, 1215, 1216, 1217, 1223, and 1226, for which the clone serial numbers are missing among those of the reported 2 groups (*cf*. Table S1 of [1]); nor did the authors state whether these 9 possible clones were simultaneously obtained in the same study, why they were not published along with the 15 reported clones, and whether they contained different fungal genetic materials, although these questions and other relevant questions should have been asked during the journal’s peer review process and should have been answered by the authors. The Editors-in-Chief of Molecular Phylogenetics and Evolution, Drs. Derek Wildman and E.A. Zimmer, encouraged and supported discussions of the relevant scientific issues in the journal with the authors [25,34].

In contrast, Li *et al*. [9], from a different research group, used culture-independent techniques and reported the cooccurrence of GC-biased Genotypes #1 and #13‒14, AT-biased Genotypes #5‒6 and #16 of *O. sinensis*, *P. hepiali*, and an AB067719-type fungus in *C. sinensis* ascospores. These findings from nonculture experiments confirmed the genetic heterogeneity of the multicellular heterokaryotic ascospores of natural *C. sinensis*. Zhu *et al*. [24,42], Gao *et al*. [26–27), and Li *et al*. [9,32) demonstrated that the *O. sinensis* genotypes underwent dynamic alterations in the immature, maturing, and mature stromata, the stromal fertile portion (SFP) that is densely covered with numerous ascocarps, and ascospores of natural *C. sinensis*, indicating the genomic independence of the *O. sinensis* genotypes and that many of the genotypic fungi may not be culturable and detectable in the *in vitro* culture-dependent experimental settings established by Li *et al*. [1].

### IV-5. Genomic variations of *H. sinensis* strains

By aggregating the information from molecular and genomic studies [1,44,50–53], different genomic DNA samples of “pure” *H. sinensis* strains were examined and published and can be summarized and grouped as follows.

1. <u>Group 1 of *H. sinensis* strains</u> consisted of pure, homokaryotic anamorphic *H. sinensis* strains (1229, CC1406-203, Co18, IOZ07, and ZJB12195) [44,50–53]. Total genomic DNA was isolated from these strains and subjected to genome-wide sequencing. The genomes contained no AT-biased genotype sequences.
2. <u>Group 2 of *H. sinensis* strains</u> consisted of 7 clones (Strains 1207, 1218, 1219, 1221, 1222, 1225, and 1229) among 15 clones derived from 25 days of incubation of natural *C. sinensis* mono-ascospores [1]. Total genomic DNA was isolated from these strains and shown to contain the homogenous ITS sequence of Genotype #1 *H. sinensis* but included no AT-biased sequences of *O. sinensis* genotypes.
3. <u>Group 3 of “*H. sinensis*” strains</u> consisted of 8 other clones (Strains 1206, 1208, 1209, 1214, 1220, 1224, 1227, and 1228) obtained from the incubation of *C. sinensis* mono-ascospores for 25 days [1]. Total genomic DNA was isolated from these strains, which exhibited genetic heterogeneity and the coexistence of GC-biased Genotype #1 and AT-biased Genotype #5 of *O. sinensis*.

The *H. sinensis* strains of Groups 1 and 2 were similar, with each strain possessing a homogenous GC-biased Genotype #1 genome, although multiple GC-biased repetitive ITS copies were identified in the genome assemblies JAAVMX000000000 and NGJJ00000000 for the *H. sinensis* Strains IOZ07 and CC1406-203, respectively (*cf*. Tables 1 and 3 and Figures 1–3) [1,44,50–53]. Although a single copy of the ITS sequence was identified in the genome assemblies ANOV00000000, LKHE00000000, and LWBQ00000000 of the *H. sinensis* Strains Co18, 1229, and ZJB12195, respectively [44,51–52], the genomes of these strains may theoretically contain many repetitive ITS copies that could have been discarded during the assembly of genome shotgun reads. Conceivably, if technically possible, reassembling the genome shotgun reads may very likely identify additional repetitive ITS copies in the genomes of Group-1 Strains Co18, 1229, and ZJB12195, among which Strain 1229 is also a Group-2 strain that was derived from mono-ascospores by Li *et al*. [1,51]. Similarly, the genome sequencing of other Group-2 strains may very likely reveal multiple repetitive ITS copies.

The *H. sinensis* strains in Groups 2 and 3 were derived after 25 days of liquid culture of the same *C. sinensis* mono-ascospores [1]; however, they differed genetically. The Group-3 strains may have heterokaryotic mycelial structures, similar to the mycelia derived from the microcycle conidiation of *H. sinensis* ascospores initially cultured on solid medium plates for 30 days and subsequently in liquid medium for 10‒53 days, as shown in Figures 2‒3 of [72]. Li *et al*. [1] reported that the Group-3 strains included cocultured GC-biased Genotype #1 and AT-biased Genotype #5 of *O. sinensis* fungi, which were genomically independent and coexisted in heterokaryotic mycelial cells, indicating either diploid/polyploid mycelial cells containing heterogeneous sets of chromosomes/genomes, or were impure mycelia, each containing a different chromosome/genome. However, the authors did not identify or report any haploid clones containing only AT-biased Genotype #5, indicating that this AT-biased *O. sinensis* fungus may not exhibit independent culturability under *in vitro* experimental settings, although other studies have reported the identification of single AT-biased Genotype #4 or #5 of *O. sinensis* without coculture with GC-biased Genotype #1 in cultures of natural and cultivated *C. sinensis* under different experimental settings [14,20,29]. Unfortunately, Li *et al*. [1,47,60] neglected the differences in genetic material between the strains in Groups 2 and 3, which were presumably derived from the same or different cells of multicellular heterokaryotic ascospores [9,34,45–46].

In addition to the genome sequencing of the homogenous Group-2 Strain 1229 and the genome assembly LKHE00000000 by Li *et al*. [51], the genome sequencing of one of the heterogeneous Group-3 strains will provide critical genomic evidence to validate the hypothesis that AT-biased Genotype #5 of *O. sinensis* coexists as an “ITS pseudogene … in a single genome” of GC-biased Genotype #1. The authors may also consider a multilocus approach to explore pseudogenes in the heterogeneous genomes of Group- 3 strains and broaden the current single locus “ITS pseudogene” hypothesis to multiple genomic loci. Unfortunately, the authors elected not to do so since 2012 or 2013 after obtaining the 8 Group-3 strains or perhaps failed in such an endeavor due to technical difficulties in the assembly of shotgun reads because of heterogeneous sequences from more than one set of genomes of independent genotypic *O. sinensis* fungi having close evolutionary relationships. Consequently, they elected willingly or unwillingly to forgo the opportunity to obtain and publish solid evidence to validate their “ITS pseudogene” hypothesis stating that AT-biased Genotype #5 and GC-biased Genotype #1 physically coexist “in a single genome”. Studies from other research groups have confirmed that the sequences of all AT-biased genotypes belong to the genomes of independent fungi [5–9,18,21,24–31,34,42,51] and that heterokaryotic ascospores and hyphae with multiple mononucleated, binucleated, trinucleated, and tetranucleated cells contain heterogeneous sets of chromosomes and genomes [45–46].

In addition to the 3 groups of *H. sinensis* strains summarized above, many other *H. sinensis* strains have been studied and presumably belong to Groups 1‒3 or other groups. For instance, Li *et al*. [73] isolated 2 *H. sinensis* strains, CH1 and CH2, from the intestine of healthy larvae of *Hepialus lagii* Yan. These strains exhibited *H. sinensis*-like morphological and growth characteristics but contained GC-biased Genotype #1, AT-biased Genotypes #4‒5, and *P. hepiali*. The wild-type strains strongly enhanced the inoculation potency of *H. sinensis* by 15- to 39-fold in the larvae (n=100 in each experimental group) of *Hepialus armoricanus* (*P*<0.001). Additionally, some “pure” *H. sinensis* strains (gifts from a senior mycologist) contained *H. sinensis* and *P. hepiali* [73]. As mentioned above, Li *et al*. [1] did not discuss whether 9 other ascosporic clones (1210, 1211, 1212, 1213, 1215, 1216, 1217, 1223, and 1226) probably simultaneously obtained in the study constituted additional fungal group(s) containing different genetic materials.

### IV-6. Genomic independence of AT-biased genotypes of *O. sinensis*

#### IV-6.1. Differential occurrence of AT-biased genotypes in different compartments of natural *C. sinensis*

A BLAST search of the GenBank database revealed at least 668 ITS sequences of *O. sinensis* (GenBank taxid: 72228), 228 of which belong to AT-biased Genotypes #4‒6 and #15‒17 [6–9,25,31,74]. Stensrud *et al*. [18] and Zhang *et al*. [41] analyzed the phylogeny of 71 and 397 ITS sequences of *O. sinensis*, respectively, and revealed 3 phylogenetic clades, *i.e.*, Groups A‒C (GC-biased Genotype #1 and AT-biased Genotypes 4‒5), which consisted of correct *O. sinensis* sequences. In contrast to the homogenous Group-A sequences reported by Stensrud *et al*. [18], the ITS sequences in Group A (Clade A) reported by Zhang *et al*. [41] appeared to be heterogeneous and diversified in a minimum evolution phylogenetic tree, consisting of Genotype #1 and other GC-biased genotypes of *O. sinensis*. Additional GC- and AT-biased *O. sinensis* genotypes have been successively discovered thereafter (*cf*. Table 2) [1,5–9,13–14,16,21–22,24–29,31–32,42].

Many early studies conducted since 1999 used a single pair of “universal” primers (*e.g*., *ITS1*‒*ITS5*) for PCR amplification and identified one of the *O. sinensis* genotypes in natural *C. sinensis* specimens collected from different production areas or cultivated *C. sinensis* [14–17,19,22,29,75–76]. Importantly, Wei *et al*. [20] reported an industrial cultivation project and identified a single teleomorphic AT-biased Genotype #4 of *O. sinensis* in cultivated *C. sinensis* and a single teleomorphic GC-biased Genotype #1 in natural *C. sinensis*, indicating the existence of at least 2 *O. sinensis* teleomorphs. Unfortunately, this study revealed a species contradiction between the anamorphic inoculants (reported to be 3 GC-biased Genotype #1 *H. sinensis* strains) and the teleomorphic AT-biased Genotype #4 of *O. sinensis* in cultivated *C. sinensis*. These findings are inconsistent with the hypotheses regarding the existence of a sole anamorph of *O. sinensis* for *H. sinensis* and a sole teleomorph of *O. sinensis* proposed by the same key authors 10 years ago [19].

Other studies have optimized and combined the use of a variety of molecular techniques, such as several pairs of genotype-specific primers, touchdown PCR, nested PCR, Southern blotting, restriction fragment length polymorphism (RFLP), cloning-based amplicon sequencing, single-nucleotide polymorphism (SNP) mass spectrometry genotyping techniques, etc. These research approaches greatly improved the specificity and efficiency of DNA amplification and detection and facilitated the identification of differential coexistence and dynamic alterations of AT-biased genotypes as well as GC-biased genotypes of *O. sinensis* in different combinations in the caterpillar body, stroma, stromal fertile portion with numerous ascocarps, and ascospores of natural *C. sinensis* during maturation (*cf*. Table 10) [5–9,21,24–28,32,42,73].

Xiao *et al*. [21] designed pairs of genotype-specific primers for the first time and adopted a nested PCR approach to amplify mutant sequences of *O. sinensis* from the stroma and caterpillar body of immature and mature *C. sinensis* specimens collected from geographically remote production areas. The authors reported the consistent occurrence of GC-biased Genotype #1 and *P. hepiali* and the inconsistent occurrence of AT-biased Genotypes #4‒5 of *O. sinensis*. The authors also uploaded the first sequence (EU555436) of AT-biased Genotype #6 to GenBank. The authors then concluded that the AT-biased genotypes likely belonged to independent *O. sinensis* fungi.

Li *et al*. [9,32] used several pairs of genotype-specific primers, touch-down PCR, cloning-based PCR amplicon sequencing, and MassARRAY SNP mass spectrometry genotyping techniques and identified Genotypes #5‒6 and #16 of AT-biased Cluster A (*cf*. Figure 2) in the ascospores of natural *C. sinensis*, cooccurring with GC-biased Genotype #1, *P. hepiali*, and an AB067719-type fungus. The Genotypes #4 and #15 sequences of AT-biased Cluster-B were identified in the stromal fertile portion, which was densely covered with numerous ascocarps, but not in ascospores obtained from the same *C. sinensis* specimens (*cf*. Table 10). Li *et al*. [9] also identified GC-biased Genotypes #13 and #14 in semiejected and fully ejected ascospores, respectively, collected from the same specimens of natural *C. sinensis*. Although Li *et al*. [77] amended their inoculant information, the inoculant used in the industrial cultivation project was *O. sinensis*, which was isolated from the ascospores of Chinese cordyceps (*i.e.*, natural *C. sinensis*) collected from an alpine meadow of Nyingchi, Tibet Autonomous Region, China, but not the 3 pure GC-biased Genotype #1 *H. sinensis* strains reported by Wei *et al*. [20]. Li *et al*. [9,32] reported that the fully ejected ascospores of natural *C. sinensis* contained GC-biased Genotypes #1 and #14 and AT-biased *O. sinensis* Genotypes #5–6 and #16 within AT-biased Cluster A (*cf*. Figure 2), *P. hepiali* and an AB067719-type fungus. Li *et al*. [1] made additional efforts to design Genotype #4-specific primers for identifying Genotype #4 of AT-biased Cluster B in mono-ascospores but failed. These studies indicated that the ascospores did not contain Genotypes #4 and #15 of AT-biased Cluster B. Thus, Wei *et al*. [20] and Li *et al*. [77] actually left an unsolved puzzle regarding the fungal source of the sole teleomorphic AT-biased Genotype #4 in cultivated *C. sinensis*.

Li *et al*. [1] did not identify the ITS sequences of AT-biased Genotypes #4, #6, and #15‒17 of *O. sinensis* in genomic DNA preparations extracted from the 15 ascosporic strains in both Groups 2 and 3 of *H. sinensis* (*cf*. Section IV-5). Thus, the expansion of the “ITS pseudogene” hypothesis from the identified AT-biased Genotype #5 from the mono-ascospore cultures to all AT-biased genotypes was apparently an overly generalized interpretation with insufficient supporting genomic evidence. As shown in Tables 1‒3 and Figures 1‒2, genomic and phylogenetic analyses did not support the hypothesis that the “diverged ITS paralogs” (*i.e.*, AT-biased Genotypes #4‒6 and #15‒17), as the “ITS pseudogene” components, coexist “in a single genome” of Genotype #1 *H. sinensis*.

IV-6.2. Dynamic alterations of multiple *O. sinensis* genotypes in different compartments of natural *C. sinensis*

Molecular physiological studies have revealed that the biomasses/abundances of *O. sinensis* genotypes are dynamically altered in an asynchronous, disproportional manner during *C. sinensis* maturation, indicating the genomic independence of multiple genotypes from different perspectives [6–9,21,24–28,31–32,42,73].

Zhu *et al*. [42] and Gao *et al*. [26–27] reported that Genotype #4 of AT-biased Cluster B of *O. sinensis* (*cf*. Figure 2) predominated in the immature stroma of natural *C. sinensis,* which was in its asexual growth stage; however, its abundance dramatically declined in the mature stroma, which was in its sexually reproductive stage. The maturational decline in Genotype #4 abundance in the *C. sinensis* stroma was accompanied by a dramatic reciprocal increase in the abundance of Genotype #5 of *O. sinensis*, which was placed in AT-biased Cluster A, indicating that the sequences of the multiple AT-biased genotypes belong to the genomes of independent *O. sinensis* fungi.

Li *et al*. [32] used the MassARRAY SNP mass spectrometry genotyping technique and found dynamic alterations in GC-biased Genotype #1 and several AT-biased genotypes of *O. sinensis* in the stromal fertile portion (densely covered with numerous ascocarps) before and after the ejection of *C. sinensis* ascospores and in fully and semiejected ascospores collected from the same *C. sinensis* specimens. A highly abundant SNP peak that contained both AT-biased Genotypes #6 and #15 was shown in the preejection stromal fertile portion of natural *C. sinensis,* but the abundance dramatically declined in the stromal fertile portion after ascospore ejection, while the coexisting AT-biased Genotype #5 maintained high abundance.

The ITS sequences of the GC-biased genotypes contained an *Eco*RI endonuclease cleavage site (G▾AATTC in purple in Figure 1) at nucleotides 294→299 within AB067721 (a representative sequence of Genotype #1 *H. sinensis*). The *Eco*RI cleavage site was lost (GAATTT) in all AT-biased genotypes due to a single-base cytosine-to-thymine (C-to-T) transition mutation (*cf*. Figure 1). To examine dynamic alterations in the natural biomasses of GC- and AT-biased genotypes of *O. sinensis* without PCR manipulation that causes nonlinear, exponential amplification of DNA sequences, the Southern blotting technique was applied to examine genomic DNA after enzymolysis with the endonucleases *Ava*I, *Dra*I, and *Eco*RI [42]. This non-PCR approach identified a single *Eco*RI-sensitive DNA moiety in genomic DNA isolated from mycelia of *H. sinensis* (*cf*. Figure 4B of [42]) and a doublet (both *Eco*RI-sensitive GC-biased Genotype #1 and *Eco*RI-resistant AT-biased genotypes) in genomic DNA isolated from natural *C. sinensis*. The lack of *Eco*RI-resistant AT-biased genotypes in pure *H. sinensis* (*cf*. Figure 4B of [42]) indicates that the sequences of AT-biased genotypes of *O. sinensis* are absent in the genome of a GC-biased Genotype #1 *H. sinensis* strain (a gift from Prof. Guo Y-L), similar to the *H. sinensis* strains in Groups 1 and 2 summarized in Section IV-5. Because Li *et al*. [1] incorrectly interpreted the biochemical findings obtained by Southern blotting [42], we have provided help to mycologists by reproducing the results (with permission) shown in Figure 8 with detailed explanations.

**Figure 8.**
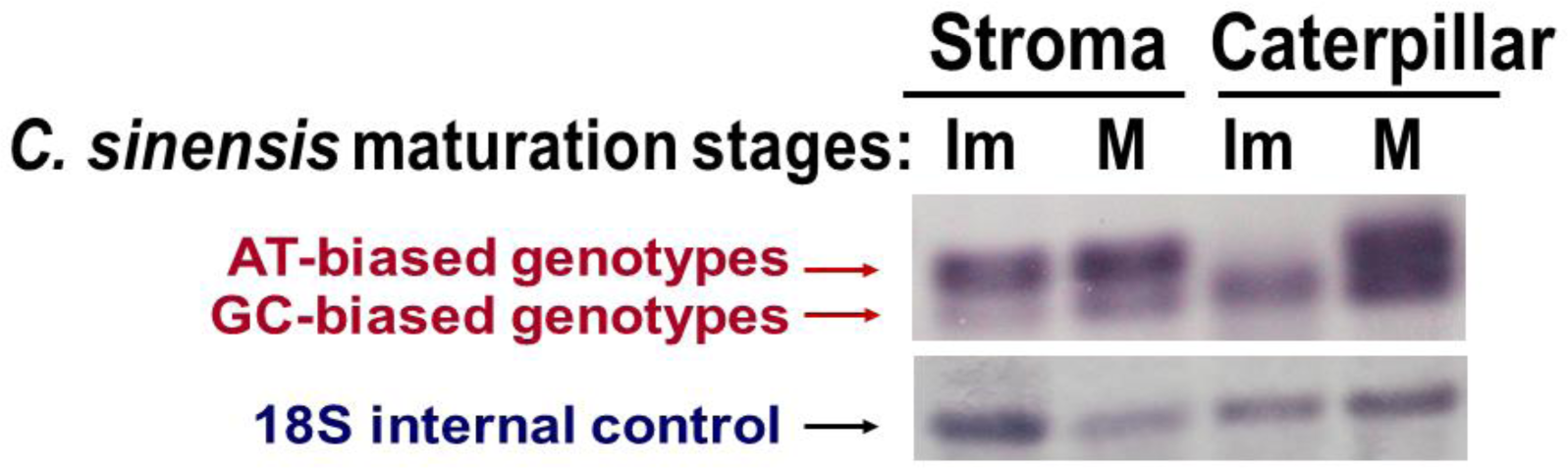
Southern blot of O. sinensis nrDNA in the stroma and caterpillar body of natural C. sinensis during maturation. [Reproduced with permission from AJBMS (www.nwpii.com/ajbms) Am J Biomed Sci 2010; 2(3): 217-238] [42]. The term “caterpillar” refers to the caterpillar body. Genomic DNA templates were isolated from the stromata or caterpillar bodies of immature (Im) or mature (M) C. sinensis specimens collected from Sichuan Province in China and prepared using the restriction enzymes AvaI, DraI, and EcoRI. **<u>The upper panel</u>** was probed with an H. sinensis-specific probe that was designed based on the ITS1 sequence of Genotype #1 H. sinensis (cf. Section II-8). **<u>The lower panel</u>** was probed with a nonspecific 18S internal control probe that was designed based on the 18S gene sequence of Genotype #1 H. sinensis, which is >700 bp upstream of the ITS1 segment.

The upper panel of Figure 8 shows the Southern blot using an *H. sinensis*-specific probe that was designed based on the ITS1 sequence of Genotype #1 *H. sinensis* (*cf*. Section II-8) and demonstrates the following:

1. <u>The *O. sinensis* genotypes identified in the stromata</u>: the left 2 lanes of the upper panel of Figure 8 show that the slower-migrating DNA moieties (*Eco*RI-resistant AT-biased genotypes) predominate naturally without PCR amplification in the stromata of immature (IM) and mature (M) *C. sinensis*. The faster migrating moieties (*Eco*RI-sensitive GC-biased *H. sinensis*) are always minor moieties with much less fluorescence intensities in the stromata during *C. sinensis* maturation, although the *H. sinensis*-specific probe was designed based on the ITS1 sequence of GC-biased Genotype #1 and showed 81.4–92.3% sequence similarity to AT-biased genotypes of *O. sinensis*.
2. <u>The *O. sinensis* genotypes identified in the caterpillar bodies</u>: although the second lane from the right in the upper panel of Figure 8 seems to contain a single *Eco*RI-sensitive moiety in the immature (IM) caterpillar body but a doublet in the mature (M) caterpillar body, as shown in the far right lane, PCR experiments using genotype-specific primers and touch-down and nested PCR protocols were able to identify *Eco*RI-resistant AT-biased genotypes in the immature caterpillar body. These results suggest that the natural quantity of *Eco*RI-resistant AT-biased genotypes in the immature caterpillar body was below the detection limit of the Southern blotting technique, which is much less sensitive than PCR-based techniques. The quantity of AT-biased genotypes dramatically and disproportionally increased from a microquantity in the immature caterpillar body to a level comparable to that of *Eco*RI- sensitive GC-biased *H. sinensis* in the mature caterpillar body.
3. <u>The *O. sinensis* genotypes identified in the stroma and caterpillar body of immature *C. sinensis*</u>: the first and third lanes from the left in Figure 8 show that the abundances and fluorescence intensity ratios of *Eco*RI-resistant AT-biased genotypes and *Eco*RI-sensitive GC-biased *H. sinensis* differed significantly in the stroma and caterpillar body of immature *C. sinensis*, which were sampled from the same specimens.
4. <u>The *O. sinensis* genotypes identified in the stroma and caterpillar body of mature *C. sinensis*</u>: the first and third lanes from the right in Figure 8 show that the abundances and fluorescence intensity ratio of *Eco*RI-resistant and *Eco*RI-sensitive genotypes differed in the stroma and caterpillar body of mature *C. sinensis*, which were sampled from the same specimens.

The disproportional distribution of GC- and AT-biased *O. sinensis* genotypes in the stromata and caterpillar bodies of natural *C. sinensis* and their asynchronous disproportional alterations in natural biomass without PCR amplification during *C. sinensis* maturation indicate that the sequences of AT-biased *O. sinensis* genotypes do not reside in the single genome of Genotype #1 *H. sinensis*. Instead, the genomically independent AT-biased *O. sinensis* genotypes are interindividually accompanied by GC-biased Genotype #1 *H. sinensis* in the stroma and caterpillar body of natural *C. sinensis*.

Zhu *et al*. [24,42] also used the *Eco*RI enzymolysis approach and restriction fragment length polymorphism (RFLP) technique and examined GC-biased Genotypes #1 and #2 of *O. sinensis* in *C. sinensis* stromata. The authors demonstrated dynamic but asynchronous maturational alterations of cooccurring GC-biased Genotypes #1 and #2 in the stromata of immature, maturing, and mature *C. sinensis* specimens, indicating genomic independence of these 2 GC-biased genotypes of *O. sinensis*. Figures 2 and 3 above show that *O. sinensis* Genotype #2 is genetically and phylogenetically distinct from the repetitive ITS copies identified in the genome of Genotype #1 *H. sinensis*, further confirming the genomic independence of the 2 GC-biased genotypes and ruling out the possibility that Genotype #2 represents a repetitive ITS copy in the single *O. sinensis* genome.

If “Different types of ITS sequences were thus proved to coexist in a single genome”, as assumed by Li *et al*. [1], the distributions of GC- and AT-biased *O. sinensis* genotypes should be proportional in the *C. sinensis* stroma and caterpillar body, and any biomass alterations of the cooccurring genotypes should be synchronous and proportional during *C. sinensis* maturation. Southern blotting, as well as RFLP after pretreatment with *Eco*RI for enzymolysis and MassARRAY SNP mass spectrometry genotyping assays, clearly demonstrated asynchronous and disproportional alterations in the GC- and AT-biased *O. sinensis* genotypes [9,24,26–28,32,42], and Figures 1‒3 and Tables 2 and 11 further confirmed the genomic independence of the multiple *O. sinensis* genotypes, the sequences of which belong to independent *O. sinensis* fungi [18,21].

After the hypothesis of “ITS pseudogenes … in a single genome” was published and subjected to academic discussion, Li *et al*. [1] had sufficient time to generate solid evidence and validate their hypothesis by conducting genome sequencing on one of the Group-3 strains (*cf*. Section IV-5). However, scientific papers addressing this topic have yet to appear. Instead, Li *et al*. [1,47,60] repeatedly stated that “Different types of ITS sequences were thus proved to coexist in a single genome”, “from a single genome, and not from different genomes”, and “The coexistence of diverged ITS paralogs in a single genome”. Either these scientists did not conduct genome sequencing of one of the Group-3 strains (*cf*. Section IV-5), or the experiments might have failed due to the high degree of similarity of the genomic sequences of multiple genotypes of *O. sinensis* that evolved from a common ancestor during a long evolutionary history, which caused difficulties in the assembly of shotgun reads to verify their hypothesis that Genotypes #1 and #5 coexist in “a single genome”, although this genome sequencing strategy may not be able to confirm the coexistence of AT-biased Genotypes #4, #6, and #15‒17 as the genomic components of “a single genome” of GC-biased Genotype #1 *H. sinensis*.

### IV-7. Possible synergistic effects of several genotypic fungi on *O. sinensis* sexual reproduction

Based on the identification of mating-type genes of both *MAT1-1* and *MAT1-2* idiomorphs in 2 *H. sinensis* strains, Hu *et al*. [44] and Bushley *et al*. [45] proposed self-fertilization as the reproductive mode of Genotype #1 *H. sinensis* under homothallism or pseudohomothallism. Accordingly, Li *et al*. [47] stated that “*O. sinensis* is one of the only known homothallic species in *Ophiocordycipitaceae* … the homothallic lifestyle may have increased the likelihood and frequency of sex in *O. sinensis*, leading to increased RIP in this species.” However, Zhang & Zhang [46] revealed differential occurrence of the mating-type genes of *MAT1-1* and *MAT1-2* idiomorphs in numerous *H. sinensis* strains or isolates, destabilizing the foundation of the self-fertilization hypothesis for *O. sinensis* at the genomic level, and the authors alternatively hypothesized facultative hybridization as the mode of *O. sinensis* reproduction.

In addition to the analysis of general intraspecific genetic variations and differential expression of *H. sinensis* genes [30–31], Li *et al*. [43] focused on the differential occurrence of mating-type genes of the *MAT1-1* and *MAT1-2* idiomorphs of 236 *H. sinensis* strains: 22 strains (9.3%) contained only the MAT1-1-1 gene, 65 (27.5%) possessed only the MAT1-2-1 gene, and 149 (63.1%) had both genes. The differential occurrence of mating-type genes underlies the genetic control of *O. sinensis* sexual reproduction. The authors also identified differential but reciprocal transcription of mating-type genes in two *H. sinensis* strains. The MAT1-1-1 transcript was absent, but the MAT1-2-1 transcript was present in the transcriptome assembly GCQL00000000 of the *H. sinensis* Strain L0106 [43,54]. The MAT1-1-1 transcript was present in the *H. sinensis* Strain 1229, but the MAT1-2-1 transcript included an unspliced intron I, which contained 3 stop codons [43,45]. Bushley *et al*. [45] specifically commented that a “comparison of DNA and cDNA sequences of MAT1-2-1 revealed only one spliced intron of 55 bp (*i.e.*, intron II)” in the *H. sinensis* Strain 1229, which “did not result from contamination of genomic DNA” because “the RNA was treated with DNase I …… until no DNA contamination could be detected”. Thus, the results indicated the production of a largely truncated and dysfunctional MAT1-2-1 protein lacking the majority of the protein encoded by exons II and III. The transcriptional silencing of one of the mating- type genes and the MAT1-2-1 transcript with an unspliced intron I constitute *O. sinensis* reproduction controls at the transcriptional and coupled transcriptional-translational levels.

Accordingly, Li *et al*. [43] demonstrated the self-sterility of GC-biased Genotype #1 *H. sinensis*, suggesting that mating partners are required for *O. sinensis* sexual reproduction. These mating partners could be intraspecific species, such as the *H. sinensis* Strains L0106 and 1229, for physiological heterothallism, regardless of whether *H. sinensis* is monoecious or dioecious [9,30–31,43,46,54]. Alternatively, the mating partners might be heterospecific fungal species that may undergo hybridization if they are able to overcome interspecific reproductive barriers. The taxonomic positions of Genotypes #2–17 of *O. sinensis* have not been determined, especially for AT-biased genotypes belonging to different *O. sinensis* fungi “even at higher taxonomic levels (genera and family)” [18] and GC-biased Genotypes #2 and #7 of *O. sinensis*, which cooccur with Genotype #1 in the same specimens of natural *C. sinensis* [16,24,42]. The coexistence of several AT-biased genotypes with GC-biased Genotype #1 in the multicellular heterokaryotic ascospores and stromal fertile portion with numerous ascocarps of natural *C. sinensis* may attract special attention and constitute a genetic basis for sexual reproduction of *O. sinensis* involving fungal hybridization [6–9,18,32,43].

In addition, Barseghyan *et al*. [78] reported the findings of macro- and micromycological studies and concluded that both psychrophilic *H. sinensis* and mesophilic *T. sinensis* were anamorphs of *O. sinensis*. Many studies reported the identification of *Tolypocladium* sp. fungi in natural *C. sinensis* [15,78-81]. Engh [13] hypothesized that *Cordyceps* and *Tolypocladium* form a fungal complex in natural *C. sinensis*, and the *Cordyceps* sequence AJ786590 that was discovered by Engh [13] reported and uploaded to the GenBank database by Stensrud *et al*. [17], a Norwegian molecular mycology group. Using a Bayesian phylogenetic algorithm, Stensrud *et al*. [18] clustered AJ786590 and other ITS sequences (AB067741‒AB067749, BD167325) into the Group-B clade of *O. sinensis*; these sequences belong to AT-biased Genotype #4 and are genetically and phylogenetically distinct from GC-biased Genotype #1 *H. sinensis* in the Group-A clade (*cf*. Figures 1‒2).

Li *et al*. [9] discovered GC-biased Genotypes #13‒14 of *O. sinensis* in the semiejected and fully ejected ascospores, respectively, of natural *C. sinensis*. These *O. sinensis* genotypes exhibit the genetic characteristics of large DNA segment reciprocal substitutions and genetic material recombination between the genomes of the 2 parental fungi, *H. sinensis* and an AB067719-type fungus. The discovery of Genotypes #13‒14 of *O. sinensis* likely indicates the possibility of fungal hybridization or parasexuality between the 2 heterospecific parental fungi.

### IV-8. Transcription of the 5.8S gene and PCR amplicons of the 5.8S-F/R primers

The gene transcription analysis (*cf*. Tables 11–13 and Figure 5) indicates that Li *et al*. [1,47,60] have neglected the evidence of transcriptional silencing of 5.8S genes demonstrated in metatranscriptomic studies in natural settings and the evidence of dynamic and unbalanced transcriptional activation and silencing of numerous fungal genes during the continuous *in vitro* fermentation or liquid incubation of Genotype #1 *H. sinensis* [30–31,54–56]. Changes in *in vitro* culture conditions and durations have significant impacts on the transcription of fungal genes (*cf*. Tables 12 and 13), which involves the transcriptional activation and silencing of numerous fungal genes. Because of the conservative evolution of 5.8S genes and many other non-rDNA genes, the incomplete study design and controversial findings reported by Li *et al*. [1] need to be verified using other sophisticated transcription methods, to address the issues whether the cDNA identified by Li *et al*. [1] was truly derived from the 5.8S gene of GC-biased Genotype #1 *H. sinensis*, from mutant AT-biased Genotype #5 of *O. sinensis*, from other colonized fungi, or even from non-rRNA genes, before confidently declaring that the 5.8S genes of the multiple AT-biased genotypes of *O. sinensis* are truly permanently nonfunctional pseudogenes under the “ITS pseudogene … in a single genome” hypothesis.

Li *et al*. [1] did not disclose the amplicon sequences amplified using 5.8SF/R primers or the PCR template of the cDNA library constructed through reverse transcription of total RNA derived from *H. sinensis* Clone 1220. Li *et al*. [60] confirmed that Li *et al*. [1] designed 5.8SF/R primers based on the sequence “in the most conserved region” of *H. sinensis* rRNA, indicating low specificity of the primers and resulting in uncertain genetic sources of the PCR amplicons. Our sequence analysis showed that the cDNA sequences detected by Li *et al*. [1] were only 90.4‒91.2% similar to the 5.8S gene sequence of Genotype #1 of *O. sinensis*. Such high dissimilarity rates (8.8‒9.6%) may not be convincingly attributed to “mismatches during reverse transcription or PCR amplification”, as explained by Li *et al*. [1]. Thus, solid evidence is lacking to conclude that the cDNA sequences were truly derived from the 5.8S gene of Genotype #1 *H. sinensis* and that the *H. sinensis* 5.8S gene was considered the functional gene, while the 5.8S gene of Genotype #5 of *O. sinensis* was considered the nonfunctional “ITS pseudogene”.

Prior to declaring that the 5.8S genes of all AT-biased *O. sinensis* genotypes are permanently nonfunctional pseudogenes cooccurring with a functional 5.8S gene in the genome of GC-biased Genotype #1, several other concerns should be addressed.

1. Scientists [62–69] have repeatedly reported epigenetic methylation of nearly all cytosine residues in 5.8S RNA genes and subsequent transcriptional silencing of GC-biased genes, which occur only in certain fungal species. As shown in Table 3 and Figures 1 and 3 above, the multiple genomic repetitive ITS copies of GC-biased Genotype #1 *H. sinensis* contained mainly insertion, deletion and transversion alleles but no or only a few transition alleles. The repetitive ITS copies of Genotype #1 are genetically and phylogenetically distinct from the AT-biased genotypes of *O. sinensis* (Figures 1–3). Although the 5.8S gene of GC-biased Genotype #1 contains many cytosine residues and is potentially more susceptible to epigenetic methylation attack, introducing nonsense or missense mutations that subsequently cause translational silencing of 5.8S genes, the evidence presented in this paper indicates that *H. sinensis* species may not be the target of RIP mutagenesis and epigenetic methylation attack. It is unlikely that the AT-biased genotypes of *O. sinensis* emerged immediately before or after the generation of a new *H. sinensis* genome; instead, they are likely genomically independent and exist in different *O. sinensis* fungi.
2. Li *et al*. [1] did not detect the ITS sequences of Genotypes #2–4 and #6–17 of *O. sinensis* in the genomic DNA pool of mono-ascosporic cultures. Thus, it is an apparent overgeneralization to infer that the 5.8S genes of these genotypes are nonfunctional “ITS pseudogene” components of the genome of Genotype #1 *H. sinensis*.
3. The sequences of Genotypes #2–17 are absent in the genome assemblies ANOV00000000, JAAVMX000000000, LKHE00000000, LWBQ00000000, and NGJJ00000000 of the *H. sinensis* Strains Co18, IOZ07, 1229, ZJB12195, and CC1406-2031229, respectively [44,50–53], indicating genomic independence of multiple *O. sinensis* genotypes.
4. The culture-dependent approach used by Li *et al*. [1] might have overlooked some fungal species that are nonculturable or difficult to culture under the *in vitro* experimental settings of the study. The culture-dependent strategy might have a significant impact on the transcription of many fungal genes, with many genes being switched on or off nonnaturally, as evidenced by the nonlinear reduction in the total number of transcriptomic unigenes and increases in the average length but decreases in the GC content of transcripts during 3‒9 days of liquid fermentation of the *H. sinensis* Strain L0106 (*cf*. Table 13) [54]. Table 12 also demonstrates that changes in the culture duration from 4‒15 days and the use of culture medium with or without adenosine or adenine supplementation had dramatic impacts on fungal gene transcription [61]. A much greater impact on differential transcriptional activation and silencing of many genes of GC-biased Genotype #1 and AT-biased Genotype #5 of *O. sinensis* may be expected after the prolonged 25-day liquid incubation period adopted by Li *et al*. [1].
5. Three distinct types of secondary steric conformations of 5.8S rRNA were predicted for Groups A–C (*i.e.*, Genotypes #1 and #4–5) of *O. sinensis* by Li *et al*. [1]. The possibility of producing circular RNA through backsplicing-exon circularization [82] should be considered during study design because these distinct steric structures may have a considerable impact on reverse transcription PCR and cDNA sequencing. In addition, the ITS1-5.8S-ITS2 sequences of Genotypes #2 and #6 may adopt different types of steric conformations [5–8,24,31,36,42]; thus, the design of other genotype-specific primers and the combined use of other molecular techniques should be considered.
6. Wei *et al*. [20] reported the identification of a single teleomorph of AT-biased Genotype #4 of *O. sinensis* in cultivated *C. sinensis* and a single teleomorph of GC-biased Genotype #1 in natural *C. sinensis*; the sequences of the 2 teleomorphs resided in the distant phylogenetic clades (*cf*. Figure 6 of [20] and Figures 1–2 of this paper). If AT-biased Genotype #4 represents a nonfunctional ITS pseudogene, as believed by Li *et al*. [1,60], the single AT-biased *O. sinensis* teleomorph in cultivated *C. sinensis* might have disturbed teleomorphic functions, leading to reproductive sterility and an abnormal, disturbed lifecycle of cultivated *C. sinensis*. Wei *et al*. [20] did not share any information about formation of stromal fertile portion and ascospore production in cultivated *C. sinensis*, which is the critical feature of *O. Sinensis* sexual reproduction in natural and cultivated *C. sinensis* [48].
7. Li *et al*. [9,31–32,34] reported the identification of teleomorphic Genotypes #4 and #15 of *O. sinensis* in AT-biased Cluster B in the stromal fertile portion densely covered with numerous ascocarps but not in ascospores collected from the same pieces of natural *C. sinensis*. AT-biased Genotypes #6 and #15 were found at high abundance in the stromal fertile portion prior to ascospore ejection, and their abundance drastically declined after ascospore ejection, while teleomorphic Genotype #5 maintained its high abundance in the stromal fertile portion before and after ascospore ejection [8–9,32].
8. The 5.8S genes of multiple *O. sinensis* genotypes may be transcriptionally activated or silenced in different developmental and maturational stages of natural *C. sinensis*. Significant changes in the proteomic expression profiles of the stroma and caterpillar body of natural *C. sinensis* during maturation provide evidence of such dynamic alterations in epigenetic, transcriptional, posttranscriptional, translational, and posttranslational modifications of numerous genes [83].

Apparently, Li *et al*. [1] improperly designed the study and provided insufficient and inconclusive evidence to support their “ITS pseudogene” hypothesis. The whole package of genes expressed in *O. sinensis* fungi and natural *C. sinensis* needs to be further explored before the 5.8S genes of AT-biased genotypes of *O. sinensis* can be defined as permanently nonfunctional pseudogenes.

### IV-9. Multilocus analysis of the repetitive copies of numerous authentic genes in the genome of Genotype #1 *H. sinensis*

In addition to the reassessment of the “ITS pseudogene” hypothesis and the underlying “RIP” as the root cause of pseudogenic mutagenesis by Li *et al*. [1,47,60] through single ITS-locus analysis of 5 *H. sinensis* genomes, we extended the genomic repetitive copy analysis to multiple authentic *H. sinensis* genes at multiple genomic loci in the *H. sinensis* genome assemblies, namely, ANOV00000000, JAAVMX000000000, LKHE00000000, LWBQ00000000, and NGJJ00000000, as well as their transcripts in the *H. sinensis* transcriptome assembly GCQL00000000. Based on the high degree of sequence similarity among the fungi in the genus *Ophiocordyceps* Petch, authentic *H. sinensis* genes were positioned and annotated in the *H. sinensis* genome through cross-reference annotation of the 1271 authentic genes in the genome assembly JAACLJ010000002, which is one of the 13 genomic contigs of the *O. camponoti-floridani* Strain EC05 [57].

Of the 1271 genes in the *H. sinensis* genome, 104 (approximately 8.2%) had repetitive copies (*cf*. Table 4); 37 of the 104 authentic genes had repetitive genomic copies in only one *H. sinensis* genome, and 67 had multiple repetitive copies in all 5 *H. sinensis* genomes. Tables 5–9 show 5 representative *H. sinensis* genes with a variety of point mutations, including various the postulated RIP-related C-to-T and G-to-A transitions and opposite T-to-C and A-to-G transitions, as well as C-to-A and G-to-T transversions and opposite A-to- C and T-to-G transversions, in addition to insertions and deletions. The point mutations caused low sequence similarity of repetitive copies compared with the sequences of authentic genes and increases, decreases or bidirectional changes in the total AT content.

In general, the repetitive copies of most authentic genes (83 of 104) at multiple *H. sinensis* genomic loci contained fewer postulated RIP-related C-to-T and G-to-A transitions than opposite T-to-C and A-to-G transitions. In contrast, the repetitive copies with large increases in the AT content included more RIP-related C-to-T and G-to-A transitions than opposite T-to-C and A-to-G transitions in only 11 of the 104 authentic genes (*cf*. Table 8). The slight increases in the AT content in the repetitive copies of 8 authentic genes (*cf.* Table 7) were the result of transversion mutations; however, the postulated RIP-related C-to-T and G-to-A transitions were less frequent than opposite T-to-C and A-to-G transitions.

As shown in Tables 5–9, the transcripts for the repetitive genomic copies of the authentic genes could be identified in the mRNA transcriptome GCQL00000000 for the *H. sinensis* Strain L0106, regardless of the increases, decreases, or bidirectional changes in the AT content and low sequence similarity of the repetitive copies compared with the authentic genes. The authentic genes and repetitive genomic copies likely encode different proteins or the same proteins with altered specificities. Thus, the genetic results of multilocus analysis and the transcription feature of the repetitive copies further invalidated the pseudogene hypothesis and its root cause of “RIP” mutagenesis, as proposed by Li *et al*. [1,47,60]. *O. sinensis*, especially the GC-biased Genotype #1 *H. sinensis*, may not be the mutagenetic target of “RIP”, which causes nonsense and missense mutations of authentic *O. sinensis* genes, concurrent epigenetic methylation attacks, and subsequent dysfunction of multiple repetitive copies of authentic genes.

## V. Conclusions

This study analyzed multiple repetitive ITS copies in the genome of GC-biased Genotype #1 *H. sinensis* and multiple genotypes of *O. sinensis* fungi and evaluated the “ITS pseudogene … in a single genome” hypothesis for all AT-biased *O. sinensis* genotypes [1,47,60]. The repetitive ITS copies in the genome of Genotype #1 contained multiple scattered insertion, deletion and transversion point mutations and may not be generated through RIP mutagenesis because theoretically RIP mutagenesis causes C-to-T and G-to-A transitions. The GC-biased repetitive ITS copies are genetically and phylogenetically distinct from AT-biased *O. sinensis* genotypes that contain multiple scattered transition alleles. Genotype #1 *H. sinensis* may not be the target of RIP mutagenesis or concurrent epigenetic methylation attack. The AT-biased genotypes are independent of the genome of Genotype #1 and might have been generated from a common ancestor through certain mutagenic and evolutionary mechanisms during the long course of evolution in response to the extremely harsh environment of the Qinghai-Tibet Plateau; these genotypes became independent *O. sinensis* fungi with independent genomes coexisting in parallel with GC-biased genotypes in natural *C. sinensis*. The analysis of multiple genomic loci revealed numerous repetitive copies of 104 (approximately 8.2%) of 1271 authentic *H. sinensis* genes. The repetitive copies contained various transition and transversion alleles and insertion and deletion point mutations, causing decreases, increases, or bidirectional changes in the AT content. However, the repetitive copies were functionally transcribed, regardless of whether the point mutations were caused by RIP-related or non-RIP-related mutagenic evolutionary mechanisms. Thus, Li *et al*. [1,47,60] provided unsound and controversial information to support their “ITS pseudogene … in a single genome” and the causative “RIP” mutagenesis hypotheses, which have not been validated by genomic and transcriptomic evidence.

## VI. Acknowledgments

This research was supported by a grant from the Science and Technology Department of Qinghai Province, China (grant number 2021-SF-A4 “Study on key technologies of conservation of natural resource and industrial upgrading of *Cordyceps sinensis*”) and by the major science and technology projects in Qinghai Province.

