## Supplementary material for "Genetic and phylogenetic features of the repetitive ITS copies in the *Hirsutella sinensis* genome and the genomically independent AT-biased *Ophiocordyceps sinensis* genotypes": Tables 5-9: Table 5.docx (from another version of this preprint: version 4, 2024-01-03)

### ***Table 5: The authentic* H. sinensis *genes for the triose-phosphate transporter (the query sequence) and the repetitive genomic copies with slight decrease (≤5%) in AT content.***

| **The subject sequence (repetitive copy)** | |  | ***vs*. The query sequence**  **(202613→203498 of NGJJ01001434 of the authentic gene for** **the triose-phosphate transporter)** | |  | **Mutation in the subject sequence comparing with the query sequence** | | | | | |  | **Transcript in mRNA transcriptome GCQL00000000** | | | |
| --- | --- | --- | --- | --- | --- | --- | --- | --- | --- | --- | --- | --- | --- | --- | --- | --- |
| ***H. sinensis* strain** | **Genomic fragment (sequence range)** |  | **Similarity** | **Change in AT content** |  | **Transitions** | | **Transversions** | | **Total point mutations** | |  | **Transcriptomic fragment** | **Similarity** | **Query coverage** | **Translated protein sequence** |
|  |  |  |  |  |  | **C-to-T**  **&**  **G-to-A** | **T-to-C**  **&**  **A-to-G** | **C-to-A**  **&**  **G-to-T** | **A-to-C**  **&**  **T-to-G** | **G or C**  **to**  **A or T** | **A or T**  **to**  **G or C** |  |  |  |  |  |
| **1229** | **LKHE01003487 (71694←72581)** |  | **99.9%** | **Nearly no change (↓ 44.2% to 44.1%)** |  |  |  |  |  |  |  |  | **[GCQL01008460](https://www.ncbi.nlm.nih.gov/nuccore/GCQL01008460" \t "lnkP32TN0VV013" \o "Show report for GCQL01008460.1) (451→1271)** | **100%** | **92%** | **[EQL01658](https://www.ncbi.nlm.nih.gov/protein/EQL01658.1?report=genbank&log$=protalign&blast_rank=2&RID=RF3KEU86016" \t "lnkRF3KEU86016" \o "Show report for EQL01658.1) (67→439)** |
|  | **LKHE01003657 (****124108←124983)** |  | **68.1%** | **↓ 44.2% to 42.1%** |  | **62** | **70** | **21** | **58** | **83** | **128** |  | **GCQL01008629 (1→821) with a 48-nt deletion** | **94.3%** | **99%** | [**EQL02567**](https://www.ncbi.nlm.nih.gov/protein/EQL02567.1?report=genbank&log$=protalign&blast_rank=1&RID=REUAB265016) **(44→316)** |
| **CC1406-203** | **NGJJ01001434 (202613→203498)** |  | **100%** | **No change (44.2%)** |  |  |  |  |  |  |  |  | [**GCQL01008460**](https://www.ncbi.nlm.nih.gov/nuccore/GCQL01008460) **(451→1080)** | **100%** | **92%** | [**EQL01658**](https://www.ncbi.nlm.nih.gov/protein/EQL01658.1?report=genbank&log$=protalign&blast_rank=2&RID=RF3KEU86016) **(67→376)** |
|  | **[NGJJ01000759](https://www.ncbi.nlm.nih.gov/nuccore/NGJJ01000759" \t "lnkPCVT0931016" \o "Show report for NGJJ01000759.1) (751126→752000)** |  | **68.1%** | **↓ 44.2% to 42.1%** |  | **62** | **70** | **21** | **58** | **83** | **128** |  | **GCQL01008629 (1→821) with a 48-nt deletion** | **94.3%** | **99%** | [**EQL02567**](https://www.ncbi.nlm.nih.gov/protein/EQL02567.1?report=genbank&log$=protalign&blast_rank=1&RID=REUAB265016) **(44→316)** |
| **Co18** | **ANOV01001461 (5923→6447)** |  | **100%** | **No change (43.5%)** |  |  |  |  |  |  |  |  | [**GCQL01008460**](https://www.ncbi.nlm.nih.gov/nuccore/GCQL01008460) **(810→1271)** | **100%** | **88%** | [**EQL01658**](https://www.ncbi.nlm.nih.gov/protein/EQL01658.1?report=genbank&log$=protalign&blast_rank=2&RID=RF3KEU86016) **(187→439)** |
|  | **ANOV01001461 (5261→5919)** |  | **99.8%** | **Nearly no change (↓ 43.9% to 43.7%)** |  |  |  |  |  |  |  |  | [**GCQL01008460**](https://www.ncbi.nlm.nih.gov/nuccore/GCQL01008460) **(451→1080)** | **100%** | **95%** | [**EQL01658**](https://www.ncbi.nlm.nih.gov/protein/EQL01658.1?report=genbank&log$=protalign&blast_rank=2&RID=RF3KEU86016) **(67→376)** |
|  | **ANOV01000747 (27035→27910)** |  | **68.1%** | **↓ 44.2% to 42.1%** |  | **62** | **70** | **21** | **58** | **83** | **128** |  | **GCQL01008629 (1→821) with a 48-nt deletion** | **94.3%** | **99%** | [**EQL02567**](https://www.ncbi.nlm.nih.gov/protein/EQL02567.1?report=genbank&log$=protalign&blast_rank=1&RID=REUAB265016) **(44→316)** |
| **IOZ07** | **JAAVMX010000003 (1710624→1711511)** |  | **99.9%** | **Nearly no change (↓ 44.2% to 44.1%)** |  |  |  |  |  |  |  |  | [**GCQL01008460**](https://www.ncbi.nlm.nih.gov/nuccore/GCQL01008460) **(451→1271)** | **100%** | **92%** | [**EQL01658**](https://www.ncbi.nlm.nih.gov/protein/EQL01658.1?report=genbank&log$=protalign&blast_rank=2&RID=RF3KEU86016) **(67→439)** |
|  | **JAAVMX010000005 (9078524←9079399)** |  | **68.1%** | **↓ 44.2% to 42.1%** |  | **62** | **70** | **21** | **58** | **83** | **128** |  | **GCQL01008629 (1→821) with a 48-nt deletion** | **94.3%** | **99%** | [**EQL02567**](https://www.ncbi.nlm.nih.gov/protein/EQL02567.1?report=genbank&log$=protalign&blast_rank=1&RID=REUAB265016) **(44→316)** |
| **ZJB12195** | **LWBQ01000001 (1755599→1756257)** |  | **99.8%** | **Nearly no change (43.9%)** |  | **0** | **1** | **0** | **0** | **0** | **1** |  | [**GCQL01008460**](https://www.ncbi.nlm.nih.gov/nuccore/GCQL01008460) **(451→1080)** | **100%** | **95%** | [**EQL01658**](https://www.ncbi.nlm.nih.gov/protein/EQL01658.1?report=genbank&log$=protalign&blast_rank=2&RID=RF3KEU86016) **(67→376)** |
|  | **LWBQ01000001 (1756277→1756590)** |  | **98.4%** | **Nearly no change (↓ 43.5% to 43.3%)** |  | **2** | **1** | **1** | **0** | **3** | **1** |  | [**GCQL01008460**](https://www.ncbi.nlm.nih.gov/nuccore/GCQL01008460) **(1022→1271)** | **100%** | **79%** | [**EQL01658**](https://www.ncbi.nlm.nih.gov/protein/EQL01658.1?report=genbank&log$=protalign&blast_rank=2&RID=RF3KEU86016) **(258→439)** |
|  | **LWBQ01000030 (268003→268878)** |  | **68.1%** | **↓ 44.2% to 42.1%** |  | **62** | **70** | **21** | **58** | **83** | **128** |  | **GCQL01008629 (1←821) with a 48-nt deletion** | **94.5%** | **99%** | [**EQL02567**](https://www.ncbi.nlm.nih.gov/protein/EQL02567.1?report=genbank&log$=protalign&blast_rank=1&RID=REUAB265016) **(44→316)** |

**Note: “****→” and “←” indicate the sequence directions of the sense and antisense chains, respectively. “↓” indicates a decrease in AT content.**
