## Supplementary material for "Genetic and phylogenetic features of the repetitive ITS copies in the *Hirsutella sinensis* genome and the genomically independent AT-biased *Ophiocordyceps sinensis* genotypes": Tables 5-9: Table 6.docx (from another version of this preprint: version 4, 2024-01-03)

### ***Table 6: The authentic* H. sinensis *genes for the heavy metal tolerance protein precursor (the query sequence) and the repetitive genomic copies with major*** ***decrease (>5%) in the AT content.***

| **The subject sequence (repetitive copy)** | |  | ***vs*. The query sequence**  **(195005→198178 of NGJJ01001482 of the authentic gene for the heavy metal tolerance protein precursor)** | |  | **Mutation in the subject sequence comparing with the query sequence** | | | | | |  | **Transcript in mRNA transcriptome GCQL00000000** | | | |
| --- | --- | --- | --- | --- | --- | --- | --- | --- | --- | --- | --- | --- | --- | --- | --- | --- |
| ***H. sinensis* strain** | **Genomic fragment (sequence range)** |  | **Similarity** | **Change in AT content** |  | **Transitions** | | **Transversions** | | **Total point mutations** | |  | **Transcriptomic fragment** | **Similarity** | **Query coverage** | **Translated protein sequence** |
|  |  |  |  |  |  | **C-to-T**  **&**  **G-to-A** | **T-to-C**  **&**  **A-to-G** | **C-to-A**  **&**  **G-to-T** | **A-to-C**  **&**  **T-to-G** | **G or C**  **to**  **A or T** | **A or T**  **to**  **G or C** |  |  |  |  |  |
| **1229** | **LKHE01001285 (2772←5944)** |  | **99.97%** | **Nearly no ∆ (41.2%)** |  |  |  |  |  |  |  |  | [**GCQL01012210**](https://www.ncbi.nlm.nih.gov/nuccore/GCQL01012210) **(442←1471)** [**GCQL01006458**](https://www.ncbi.nlm.nih.gov/nuccore/GCQL01006458) **(2←683)** [**GCQL01000615**](https://www.ncbi.nlm.nih.gov/nuccore/GCQL01000615) **(1←403)** | **100% 99.9% 100%** | **32% 21% 12%** | [**EQK98634**](https://www.ncbi.nlm.nih.gov/protein/EQK98634.1?report=genbank&log$=protalign&blast_rank=1&RID=REW7ECTF013) **(1→344)** [**EQK98634**](https://www.ncbi.nlm.nih.gov/protein/EQK98634.1?report=genbank&log$=protalign&blast_rank=1&RID=REW7ECTF013) **(572→798)** [**EQK98634**](https://www.ncbi.nlm.nih.gov/protein/EQK98634.1?report=genbank&log$=protalign&blast_rank=1&RID=REW7ECTF013) **(796→929)** |
|  | **LKHE01002445 (3819←4233)** |  | **65.4%** | **↓ 39.5% to 33.3%** |  | **14** | **26** | **14** | **27** | **28** | **53** |  | [**GCQL01017466**](https://www.ncbi.nlm.nih.gov/nuccore/GCQL01017466) **(726←1140)** | **100%** | **100%** | [**EQK98286**](https://www.ncbi.nlm.nih.gov/protein/EQK98286.1?report=genbank&log$=protalign&blast_rank=1&RID=REWFHX1U013) **(378→515)** |
|  | **LKHE01001747 (128317←128839; 128885←129014)** |  | **67.4%** | **↓ 38.3% to 34.6%** |  | **24** | **34** | **21** | **34** | **45** | **68** |  | [**GCQL01013313**](https://www.ncbi.nlm.nih.gov/nuccore/GCQL01013313) **(1←426)** [**GCQL01016959**](https://www.ncbi.nlm.nih.gov/nuccore/GCQL01016959) **(1→271)** | **99.8% 100%** | **61% 38%** | [**EQK99315**](https://www.ncbi.nlm.nih.gov/protein/EQK99315.1?report=genbank&log$=protalign&blast_rank=1&RID=REWC9H69013) **(592→732)** [**EQK99315**](https://www.ncbi.nlm.nih.gov/protein/EQK99315.1?report=genbank&log$=protalign&blast_rank=1&RID=REWC9H69013) **(734→823)** |
| **CC1406-203** | **NGJJ01001482 (195005→198178)** |  | **100%** | **No ∆ (40.2%)** |  |  |  |  |  |  |  |  | [**GCQL01012210**](https://www.ncbi.nlm.nih.gov/nuccore/GCQL01012210) **(442←1471)** [**GCQL01006458**](https://www.ncbi.nlm.nih.gov/nuccore/GCQL01006458) **(2←683)** [**GCQL01000615**](https://www.ncbi.nlm.nih.gov/nuccore/GCQL01000615) **(1←403)** | **100% 99.9% 100%** | **32% 21% 12%** | [**EQK98634**](https://www.ncbi.nlm.nih.gov/protein/EQK98634.1?report=genbank&log$=protalign&blast_rank=1&RID=REW7ECTF013) **(1→344)** [**EQK98634**](https://www.ncbi.nlm.nih.gov/protein/EQK98634.1?report=genbank&log$=protalign&blast_rank=1&RID=REW7ECTF013) **(572→798)** [**EQK98634**](https://www.ncbi.nlm.nih.gov/protein/EQK98634.1?report=genbank&log$=protalign&blast_rank=1&RID=REW7ECTF013) **(796→929)** |
|  | **NGJJ01001243 (150283→150697)** |  | **65.4%** | **↓ 39.5% to 33.3%** |  | **14** | **26** | **14** | **27** | **28** | **53** |  | [**GCQL01017466**](https://www.ncbi.nlm.nih.gov/nuccore/GCQL01017466) **(726←1140)** | **100%** | **100%** | [**EQK98286**](https://www.ncbi.nlm.nih.gov/protein/EQK98286.1?report=genbank&log$=protalign&blast_rank=1&RID=REWFHX1U013) **(378→515)** |
|  | **NGJJ01001310 (461774←461903; 461949←462471)** |  | **67.4%** | **↓ 38.3% to 34.6%)** |  | **24** | **34** | **21** | **34** | **45** | **68** |  | [**GCQL01013313**](https://www.ncbi.nlm.nih.gov/nuccore/GCQL01013313) **(1→426)** [**GCQL01016959**](https://www.ncbi.nlm.nih.gov/nuccore/GCQL01016959) **(1←271)** | **99.8% 100%** | **61% 38%** | [**EQK99315**](https://www.ncbi.nlm.nih.gov/protein/EQK99315.1?report=genbank&log$=protalign&blast_rank=1&RID=REWC9H69013) **(592→732)** [**EQK99315**](https://www.ncbi.nlm.nih.gov/protein/EQK99315.1?report=genbank&log$=protalign&blast_rank=1&RID=REWC9H69013) **(734→823)** |
| **Co18** | **ANOV01008830 (2485←5657)** |  | **99.97%** | **Nearly no ∆ (40.2%)** |  |  |  |  |  |  |  |  | [**GCQL01012210**](https://www.ncbi.nlm.nih.gov/nuccore/GCQL01012210) **(442←1471)** [**GCQL01006458**](https://www.ncbi.nlm.nih.gov/nuccore/GCQL01006458) **(2←683)** [**GCQL01000615**](https://www.ncbi.nlm.nih.gov/nuccore/GCQL01000615) **(1←403)** | **100% 99.9% 100%** | **32% 21% 12%** | [**EQK98634**](https://www.ncbi.nlm.nih.gov/protein/EQK98634.1?report=genbank&log$=protalign&blast_rank=1&RID=REW7ECTF013) **(1→344)** [**EQK98634**](https://www.ncbi.nlm.nih.gov/protein/EQK98634.1?report=genbank&log$=protalign&blast_rank=1&RID=REW7ECTF013) **(572→798)** [**EQK98634**](https://www.ncbi.nlm.nih.gov/protein/EQK98634.1?report=genbank&log$=protalign&blast_rank=1&RID=REW7ECTF013) **(796→929)** |
|  | **ANOV01010958 (3175→3589)** |  | **65.4%** | **↓ 39.5% to 33.3%** |  | **14** | **26** | **14** | **27** | **28** | **53** |  | [**GCQL01017466**](https://www.ncbi.nlm.nih.gov/nuccore/GCQL01017466) **(726←1140)** | **100%** | **100%** | [**EQK98286**](https://www.ncbi.nlm.nih.gov/protein/EQK98286.1?report=genbank&log$=protalign&blast_rank=1&RID=REWFHX1U013) **(378→515)** |
|  | [**ANOV01005573**](https://www.ncbi.nlm.nih.gov/nuccore/ANOV01005573) **(4224←4746; 4792←4921)** |  | **67.4%** | **↓ 38.3% to 34.6%** |  | **24** | **34** | **21** | **34** | **45** | **68** |  | [**GCQL01013313**](https://www.ncbi.nlm.nih.gov/nuccore/GCQL01013313) **(1←426)** [**GCQL01016959**](https://www.ncbi.nlm.nih.gov/nuccore/GCQL01016959) **(1→271)** | **99.8% 100%** | **61% 38%** | [**EQK99315**](https://www.ncbi.nlm.nih.gov/protein/EQK99315.1?report=genbank&log$=protalign&blast_rank=1&RID=REWC9H69013) **(592→732)** [**EQK99315**](https://www.ncbi.nlm.nih.gov/protein/EQK99315.1?report=genbank&log$=protalign&blast_rank=1&RID=REWC9H69013) **(734→823)** |
| **IOZ07** | **JAAVMX010000003 (1309808←1312980)** |  | **99.97%** | **Nearly no ∆ (40.2%)** |  |  |  |  |  |  |  |  | [**GCQL01012210**](https://www.ncbi.nlm.nih.gov/nuccore/GCQL01012210) **(442←1471)** [**GCQL01006458**](https://www.ncbi.nlm.nih.gov/nuccore/GCQL01006458) **(2←683)** [**GCQL01000615**](https://www.ncbi.nlm.nih.gov/nuccore/GCQL01000615) **(1←403)** | **100% 99.9% 100%** | **32% 21% 12%** | [**EQK98634**](https://www.ncbi.nlm.nih.gov/protein/EQK98634.1?report=genbank&log$=protalign&blast_rank=1&RID=REW7ECTF013) **(1→344)** [**EQK98634**](https://www.ncbi.nlm.nih.gov/protein/EQK98634.1?report=genbank&log$=protalign&blast_rank=1&RID=REW7ECTF013) **(572→798)** [**EQK98634**](https://www.ncbi.nlm.nih.gov/protein/EQK98634.1?report=genbank&log$=protalign&blast_rank=1&RID=REW7ECTF013) **(796→929)** |
|  | **JAAVMX010000002 (833901←834315)** |  | **65.4%** | **↓ 39.5% to 33.3%** |  | **14** | **26** | **14** | **27** | **28** | **53** |  | [**GCQL01017466**](https://www.ncbi.nlm.nih.gov/nuccore/GCQL01017466) **(726←1140)** | **100%** | **100%** | [**EQK98286**](https://www.ncbi.nlm.nih.gov/protein/EQK98286.1?report=genbank&log$=protalign&blast_rank=1&RID=REWFHX1U013) **(378→515)** |
|  | **JAAVMX010000012 (85112←85634; 85680←85809)** |  | **67.4%** | **↓ 38.3% to 34.6%** |  | **24** | **34** | **21** | **34** | **45** | **68** |  | [**GCQL01013313**](https://www.ncbi.nlm.nih.gov/nuccore/GCQL01013313) **(1←426)** [**GCQL01016959**](https://www.ncbi.nlm.nih.gov/nuccore/GCQL01016959) **(1→271)** | **99.8% 100%** | **61% 38%** | [**EQK99315**](https://www.ncbi.nlm.nih.gov/protein/EQK99315.1?report=genbank&log$=protalign&blast_rank=1&RID=REWC9H69013) **(592→732)** [**EQK99315**](https://www.ncbi.nlm.nih.gov/protein/EQK99315.1?report=genbank&log$=protalign&blast_rank=1&RID=REWC9H69013) **(734→823)** |
| **ZJB12195** | **LWBQ01000001 (****1338987←1342159)** |  | **99.97%** | **Nearly no ∆ (41.2%)** |  |  |  |  |  |  |  |  | [**GCQL01012210**](https://www.ncbi.nlm.nih.gov/nuccore/GCQL01012210) **(442←1471)** [**GCQL01006458**](https://www.ncbi.nlm.nih.gov/nuccore/GCQL01006458) **(2←683)** [**GCQL01000615**](https://www.ncbi.nlm.nih.gov/nuccore/GCQL01000615) **(1←403)** | **100% 99.9% 100%** | **32% 21% 12%** | [**EQK98634**](https://www.ncbi.nlm.nih.gov/protein/EQK98634.1?report=genbank&log$=protalign&blast_rank=1&RID=REW7ECTF013) **(1→344)** [**EQK98634**](https://www.ncbi.nlm.nih.gov/protein/EQK98634.1?report=genbank&log$=protalign&blast_rank=1&RID=REW7ECTF013) **(572→798)** [**EQK98634**](https://www.ncbi.nlm.nih.gov/protein/EQK98634.1?report=genbank&log$=protalign&blast_rank=1&RID=REW7ECTF013) **(796→929)** |
|  | **LWBQ01000010 (468456←468870)** |  | **65.4%** | **↓ 39.5% to 33.3%** |  | **14** | **26** | **14** | **27** | **28** | **53** |  | [**GCQL01017466**](https://www.ncbi.nlm.nih.gov/nuccore/GCQL01017466) **(726←1140)** | **100%** | **100%** | [**EQK98286**](https://www.ncbi.nlm.nih.gov/protein/EQK98286.1?report=genbank&log$=protalign&blast_rank=1&RID=REWFHX1U013) **(378→515)** |
|  | [**LWBQ01000044**](https://www.ncbi.nlm.nih.gov/nuccore/LWBQ01000044) **(141513→142035; 142081→142210)** |  | **67.4%** | **↓ 38.3% to 34.6%** |  | **24** | **34** | **21** | **34** | **45** | **68** |  | [**GCQL01013313**](https://www.ncbi.nlm.nih.gov/nuccore/GCQL01013313) **(1←426)** [**GCQL01016959**](https://www.ncbi.nlm.nih.gov/nuccore/GCQL01016959) **(1→271)** | **99.8% 100%** | **61% 38%** | [**EQK99315**](https://www.ncbi.nlm.nih.gov/protein/EQK99315.1?report=genbank&log$=protalign&blast_rank=1&RID=REWC9H69013) **(592→732)** [**EQK99315**](https://www.ncbi.nlm.nih.gov/protein/EQK99315.1?report=genbank&log$=protalign&blast_rank=1&RID=REWC9H69013) **(734→823)** |

**Note: “****→” and “←” indicate the sequence directions of the sense and antisense chains, respectively. “↓” indicates a decrease in AT content.**
