## Supplementary material for "Genetic and phylogenetic features of the repetitive ITS copies in the *Hirsutella sinensis* genome and the genomically independent AT-biased *Ophiocordyceps sinensis* genotypes": Tables 5-9: Table 7.docx (from another version of this preprint: version 4, 2024-01-03)

### ***Table 7: The authentic* H. sinensis *genes for the maltose permease (query sequence) and the repetitive genomic copies with minor increase (≤5%) in the AT content.***

| **The subject sequence (repetitive copy)** | |  | ***vs*. The query sequence**  **(47693→49623 of** [**LKHE01002657**](https://www.ncbi.nlm.nih.gov/sites/entrez?cmd=Search&db=nucleotide&term=LKHE01002657.1&dopt=GenBank) **of the authentic gene for the** **maltose permease)** | |  | **Mutation in the subject sequence comparing with the query sequence** | | | | | |  | **Transcript in mRNA transcriptome GCQL00000000** | | | |
| --- | --- | --- | --- | --- | --- | --- | --- | --- | --- | --- | --- | --- | --- | --- | --- | --- |
| ***H. sinensis* strain** | **Genomic fragment (sequence range)** |  | **Similarity** | **Change in AT content** |  | **Transitions** | | **Transversions** | | **Total point mutations** | |  | **Transcriptomic fragment** | **Similarity** | **Query coverage** | **Translated protein sequence** |
|  |  |  |  |  |  | **C-to-T**  **&**  **G-to-A** | **T-to-C**  **&**  **A-to-G** | **C-to-A**  **&**  **G-to-T** | **A-to-C**  **&**  **T-to-G** | **G or C**  **to**  **A or T** | **A or T**  **to**  **G or C** |  |  |  |  |  |
| **1229** | [**LKHE01002657**](https://www.ncbi.nlm.nih.gov/sites/entrez?cmd=Search&db=nucleotide&term=LKHE01002657.1&dopt=GenBank) **(47693→49623)** |  | **100%** | **No ∆ (40.1%)** |  |  |  |  |  |  |  |  | [**GCQL01010488 (586→1622)**](https://www.ncbi.nlm.nih.gov/nuccore/GCQL01010488) | **99.9%** | **80%** | [**EQK98556**](https://www.ncbi.nlm.nih.gov/protein/EQK98556.1?report=genbank&log$=protalign&blast_rank=1&RID=REY3K5XS016) **(189→533)** |
|  | **LKHE01000336 (6824→7385)** |  | **69.6%** | **↑ 37.9% to 39.3%** |  | **30** | **35** | **28** | **11** | **58** | **46** |  | **[GCQL01015773](https://www.ncbi.nlm.nih.gov/nuccore/GCQL01015773" \t "lnkP5J5JJA101N" \o "Show report for GCQL01015773.1) (948→1509) or** [**GCQL01019033 (526→1063)**](https://www.ncbi.nlm.nih.gov/nuccore/GCQL01019033) **with a 25-nt deletion** | **100% 95.6%** | **100% 100%** | [**EQL02995**](https://www.ncbi.nlm.nih.gov/protein/EQL02995.1?report=genbank&log$=protalign&blast_rank=2&RID=RF5SZSD101N) **(152→338)** [**EQL02995**](https://www.ncbi.nlm.nih.gov/protein/EQL02995.1?report=genbank&log$=protalign&blast_rank=2&RID=RF5SZSD101N) **(152→338)** |
| **CC1406-203** | **NGJJ01000203 (77988→79917)** |  | **99.8%** | **Nearly no ∆ (↑ 40.1% to 40.2%)** |  | **1** | **0** | **0** | **0** | **1** | **0** |  | [**GCQL01010488 (334→1892)**](https://www.ncbi.nlm.nih.gov/nuccore/GCQL01010488) | **99.9%** | **80%** | [**EQK98556**](https://www.ncbi.nlm.nih.gov/protein/EQK98556.1?report=genbank&log$=protalign&blast_rank=1&RID=REY3K5XS016) **(189→533)** |
|  | **NGJJ01001217 (782871←782913; 1076399←1076960)** |  | **70.4%** | **↑ 36.8% to 38.5%** |  | **31** | **37** | **28** | **10** | **59** | **47** |  | [**GCQL01015773**](https://www.ncbi.nlm.nih.gov/nuccore/GCQL01015773) **(948→1509) or** [**GCQL01019033 (526→1063)**](https://www.ncbi.nlm.nih.gov/nuccore/GCQL01019033) **with a 25-nt deletion** | **100% 95.6%** | **100% 100%** | [**EQL02995**](https://www.ncbi.nlm.nih.gov/protein/EQL02995.1?report=genbank&log$=protalign&blast_rank=2&RID=RF5SZSD101N) **(152→338)** [**EQL02995**](https://www.ncbi.nlm.nih.gov/protein/EQL02995.1?report=genbank&log$=protalign&blast_rank=2&RID=RF5SZSD101N) **(152→338)** |
| **Co18** | **ANOV01009266 (1725←3655)** |  | **99.8%** | **Nearly no ∆ (↓ 40.1% to 40.0%)** |  | **1** | **2** | **0** | **0** | **1** | **2** |  | [**GCQL01010488 (586→1622)**](https://www.ncbi.nlm.nih.gov/nuccore/GCQL01010488) | **99.8%** | **80%** | [**EQK98556**](https://www.ncbi.nlm.nih.gov/protein/EQK98556.1?report=genbank&log$=protalign&blast_rank=1&RID=REY3K5XS016) **(189→533)** |
|  | **ANOV01000509 (4433←4994, 5254←5290)** |  | **70.1%** | **↑ 37.9% to 39.9%** |  | **31** | **37** | **28** | **10** | **59** | **47** |  | [**GCQL01015773**](https://www.ncbi.nlm.nih.gov/nuccore/GCQL01015773) **(948→1509) or** [**GCQL01019033 (526→1063)**](https://www.ncbi.nlm.nih.gov/nuccore/GCQL01019033) **with a 25-nt deletion** | **100% 95.6%** | **100% 100%** | [**EQL02995**](https://www.ncbi.nlm.nih.gov/protein/EQL02995.1?report=genbank&log$=protalign&blast_rank=2&RID=RF5SZSD101N) **(152→338)** [**EQL02995**](https://www.ncbi.nlm.nih.gov/protein/EQL02995.1?report=genbank&log$=protalign&blast_rank=2&RID=RF5SZSD101N) **(152→338)** |
| **IOZ07** | **JAAVMX010000006 (4614206→4616136)** |  | **99.9%** | **Nearly no ∆ (40.1%)** |  | **1** | **0** | **0** | **0** | **1** | **0** |  | [**GCQL01010488 (334←1892)**](https://www.ncbi.nlm.nih.gov/nuccore/GCQL01010488) | **100%** | **80%** | [**EQK98556**](https://www.ncbi.nlm.nih.gov/protein/EQK98556.1?report=genbank&log$=protalign&blast_rank=1&RID=REY3K5XS016) **(189→533)** |
|  | **JAAVMX010000005 (12448464→12449025, 12779815→12779864)** |  | **70.4%** | **↑ 37.6% to 39.2%** |  | **31** | **37** | **28** | **10** | **59** | **47** |  | [**GCQL01015773**](https://www.ncbi.nlm.nih.gov/nuccore/GCQL01015773) **(948→1509) or** [**GCQL01019033 (526→1063)**](https://www.ncbi.nlm.nih.gov/nuccore/GCQL01019033) **with a 25-nt deletion** | **100% 95.6%** | **100% 100%** | [**EQL02995**](https://www.ncbi.nlm.nih.gov/protein/EQL02995.1?report=genbank&log$=protalign&blast_rank=2&RID=RF5SZSD101N) **(152→338)** [**EQL02995**](https://www.ncbi.nlm.nih.gov/protein/EQL02995.1?report=genbank&log$=protalign&blast_rank=2&RID=RF5SZSD101N) **(152→338)** |
| **ZJB12195** | **LWBQ01000157 (66747→68677)** |  | **99.9%** | **Nearly no ∆ (40.1%)** |  | **1** | **0** | **0** | **0** | **1** | **0** |  | [**GCQL01010488 (586→1622)**](https://www.ncbi.nlm.nih.gov/nuccore/GCQL01010488) | **100%** | **80%** | [**EQK98556**](https://www.ncbi.nlm.nih.gov/protein/EQK98556.1?report=genbank&log$=protalign&blast_rank=1&RID=REY3K5XS016) **(189→533)** |
|  | **LWBQ01000003 (292175→292217, 570642→571203)** |  | **70.4%** | **↑ 36.8% to 38.5%** |  | **30** | **35** | **28** | **11** | **58** | **46** |  | [**GCQL01015773**](https://www.ncbi.nlm.nih.gov/nuccore/GCQL01015773) **(948→1509) or** [**GCQL01019033 (526→1063)**](https://www.ncbi.nlm.nih.gov/nuccore/GCQL01019033) **with a 25-nt deletion** | **100% 95.6%** | **100% 100%** | [**EQL02995**](https://www.ncbi.nlm.nih.gov/protein/EQL02995.1?report=genbank&log$=protalign&blast_rank=2&RID=RF5SZSD101N) **(152→338)** [**EQL02995**](https://www.ncbi.nlm.nih.gov/protein/EQL02995.1?report=genbank&log$=protalign&blast_rank=2&RID=RF5SZSD101N) **(152→338)** |

**Note: “****→” and “←” indicate the sequence directions of the sense and antisense chains, respectively. “↑” and “↓” indicate increases and decreases in AT content.**
