## Supplementary material for "Genetic and phylogenetic features of the repetitive ITS copies in the *Hirsutella sinensis* genome and the genomically independent AT-biased *Ophiocordyceps sinensis* genotypes": Tables 5-9: Table 8.docx (from another version of this preprint: version 4, 2024-01-03)

### ***Table 8: The authentic* H. sinensis *genes for the lipase/serine esterase (query sequence) and the repetitive genomic copies with major increase (>5%) in the AT content.***

| **The subject sequence (repetitive copy)** | |  | ***vs*. The query sequence**  **(235828→236719 of NGJJ01000647 of the authentic gene for the lipase/serine esterase)** | |  | **Mutation in the subject sequence comparing with the query sequence** | | | | | |  | **Transcript in mRNA transcriptome GCQL00000000** | | | |
| --- | --- | --- | --- | --- | --- | --- | --- | --- | --- | --- | --- | --- | --- | --- | --- | --- |
| ***H. sinensis* strain** | **Genomic fragment (sequence range)** |  | **Similarity** | **Change in AT content** |  | **Transitions** | | **Transversions** | | **Total point mutations** | |  | **Transcriptomic fragment** | **Similarity** | **Query coverage** | **Translated protein sequence** |
|  |  |  |  |  |  | **C-to-T**  **&**  **G-to-A** | **T-to-C**  **&**  **A-to-G** | **C-to-A**  **&**  **G-to-T** | **A-to-C**  **&**  **T-to-G** | **G or C**  **to**  **A or T** | **A or T**  **to**  **G or C** |  |  |  |  |  |
| **1229** | **LKHE01002746 (22031→22947)** |  | **100%** | **No ∆ (34.5%)** |  |  |  |  |  |  |  |  | [**GCQL01000519**](https://www.ncbi.nlm.nih.gov/nuccore/GCQL01000519) **(1663→2554) or** [**GCQL01015314**](https://www.ncbi.nlm.nih.gov/nuccore/GCQL01015314) **(2847→3477)** | **100% 100%** | **100% 70%** | [**EQL04141**](https://www.ncbi.nlm.nih.gov/protein/EQL04141.1?report=genbank&log$=protalign&blast_rank=1&RID=REYZYNV5013) **(325→621)** [**EQL04141**](https://www.ncbi.nlm.nih.gov/protein/EQL04141.1?report=genbank&log$=protalign&blast_rank=1&RID=REYZYNV5013) **(270→479)** |
|  | **LKHE01001863 (39965→40763; 40848→40982)** |  | **68.5%** | **↑ 35.0% to 41.6%** |  | **63** | **30** | **41** | **20** | **104** | **50** |  | [**GCQL01000532**](https://www.ncbi.nlm.nih.gov/nuccore/GCQL01000532) **(5013→6030)** | **100%** | **100%** | [**EQL03018**](https://www.ncbi.nlm.nih.gov/protein/EQL03018.1?report=genbank&log$=protalign&blast_rank=2&RID=RF5D4UYG016) **(265→587)** |
| **CC1406-203** | [**NGJJ01000647**](https://www.ncbi.nlm.nih.gov/nuccore/NGJJ01000647) **(235828→236719)** |  | **100%** |  |  |  |  |  |  |  |  |  | [**GCQL01000519**](https://www.ncbi.nlm.nih.gov/nuccore/GCQL01000519) **(1663→2554) or** [**GCQL01015314**](https://www.ncbi.nlm.nih.gov/nuccore/GCQL01015314) **(2847→3477)** | **100% 100%** | **100% 70%** | [**EQL04141**](https://www.ncbi.nlm.nih.gov/protein/EQL04141.1?report=genbank&log$=protalign&blast_rank=1&RID=REYZYNV5013) **(325→621)** [**EQL04141**](https://www.ncbi.nlm.nih.gov/protein/EQL04141.1?report=genbank&log$=protalign&blast_rank=1&RID=REYZYNV5013) **(270→479)** |
|  | **NGJJ01001576 (1146814→1147612; 1147697→1147831)** |  | **68.5%** | **↑ 35.0% to 41.6%** |  | **63** | **30** | **41** | **20** | **104** | **50** |  | [**GCQL01000532**](https://www.ncbi.nlm.nih.gov/nuccore/GCQL01000532) **(5013→6030)** | **100%** | **100%** | [**EQL03018**](https://www.ncbi.nlm.nih.gov/protein/EQL03018.1?report=genbank&log$=protalign&blast_rank=2&RID=RF5D4UYG016) **(265→587)** |
| **Co18** | **ANOV01000049 (9368→10284)** |  | **99.8%** | **No ∆ (34.5%)** |  |  |  |  |  |  |  |  | [**GCQL01000519**](https://www.ncbi.nlm.nih.gov/nuccore/GCQL01000519) **(1638→2554) or** [**GCQL01015314**](https://www.ncbi.nlm.nih.gov/nuccore/GCQL01015314) **(2847→3477)** | **100% 100%** | **100% 68%** | [**EQL04141**](https://www.ncbi.nlm.nih.gov/protein/EQL04141.1?report=genbank&log$=protalign&blast_rank=1&RID=REYZYNV5013) **(325→621)** [**EQL04141**](https://www.ncbi.nlm.nih.gov/protein/EQL04141.1?report=genbank&log$=protalign&blast_rank=1&RID=REYZYNV5013) **(270→479)** |
|  | **ANOV01000484 (6319←6453; 6538←7335)** |  | **67.0%** | **↑ 34.9% to 41.5%** |  | **63** | **30** | **41** | **20** | **104** | **50** |  | [**GCQL01000532**](https://www.ncbi.nlm.nih.gov/nuccore/GCQL01000532) **(5013←6030)** | **99.7%** | **100%** | [**EQL03018**](https://www.ncbi.nlm.nih.gov/protein/EQL03018.1?report=genbank&log$=protalign&blast_rank=2&RID=RF5D4UYG016) **(265→587)** |
| **IOZ07** | **JAAVMX010000005 (10692268←10693159)** |  | **100%** | **No ∆ (34.5%)** |  |  |  |  |  |  |  |  | [**GCQL01000519**](https://www.ncbi.nlm.nih.gov/nuccore/GCQL01000519) **(1663→2554) or** [**GCQL01015314**](https://www.ncbi.nlm.nih.gov/nuccore/GCQL01015314) **(2847→3477)** | **100% 100%** | **100% 70%** | [**EQL04141**](https://www.ncbi.nlm.nih.gov/protein/EQL04141.1?report=genbank&log$=protalign&blast_rank=1&RID=REYZYNV5013) **(325→621)** [**EQL04141**](https://www.ncbi.nlm.nih.gov/protein/EQL04141.1?report=genbank&log$=protalign&blast_rank=1&RID=REYZYNV5013) **(270→479)** |
|  | **JAAVMX010000001 (22221540←22222338)** |  | **67.1%** | **↑ 34.9% to 41.6%** |  | **63** | **30** | **41** | **20** | **104** | **50** |  | [**GCQL01000532**](https://www.ncbi.nlm.nih.gov/nuccore/GCQL01000532) **(5013←5811)** | **100%** | **100%** | [**EQL03018**](https://www.ncbi.nlm.nih.gov/protein/EQL03018.1?report=genbank&log$=protalign&blast_rank=2&RID=RF5D4UYG016) **(265→587)** |
| **ZJB12195** | **LWBQ01000026 (203574←204490)** |  | **100%** | **No ∆ (34.5%)** |  |  |  |  |  |  |  |  | [**GCQL01000519**](https://www.ncbi.nlm.nih.gov/nuccore/GCQL01000519) **(1638←2554) or** [**GCQL01015314**](https://www.ncbi.nlm.nih.gov/nuccore/GCQL01015314) **(2847→3477)** | **100% 100%** | **100% 68%** | [**EQL04141**](https://www.ncbi.nlm.nih.gov/protein/EQL04141.1?report=genbank&log$=protalign&blast_rank=1&RID=REYZYNV5013) **(325→621)** [**EQL04141**](https://www.ncbi.nlm.nih.gov/protein/EQL04141.1?report=genbank&log$=protalign&blast_rank=1&RID=REYZYNV5013) **(270→479)** |
|  | **LWBQ01000013 (513365→514163; 514248→514382)** |  | **68.5%** | **↑ 35.0% to 41.6%** |  | **63** | **30** | **41** | **20** | **104** | **50** |  | [**GCQL01000532**](https://www.ncbi.nlm.nih.gov/nuccore/GCQL01000532) **(5013→5811)** | **100%** | **100%** | [**EQL03018**](https://www.ncbi.nlm.nih.gov/protein/EQL03018.1?report=genbank&log$=protalign&blast_rank=2&RID=RF5D4UYG016) **(265→587)** |

**Note: “****→” and “←” indicate the sequence directions of the sense and antisense chains, respectively. “↑” indicates an increase in AT content.**
