## Supplementary material for "Genetic and phylogenetic features of the repetitive ITS copies in the *Hirsutella sinensis* genome and the genomically independent AT-biased *Ophiocordyceps sinensis* genotypes": Tables 5-9: Table 9.docx (from another version of this preprint: version 4, 2024-01-03)

### ***Table 9: The authentic* H. sinensis *genes for the β-lactamase/transpeptidase-like protein (the query sequence) and the repetitive genomic copies with bidirectional changes in AT content.***

| **The subject sequence (repetitive copy)** | |  | ***vs*. The query sequence**  **(70191→71142 of NGJJ01001580 of the authentic gene for the β-lactamase/transpeptidase-like protein)** | |  | **Mutation in the subject sequence comparing with the query sequence** | | | | | |  | **Transcript in mRNA transcriptome GCQL00000000** | | | |
| --- | --- | --- | --- | --- | --- | --- | --- | --- | --- | --- | --- | --- | --- | --- | --- | --- |
| ***H. sinensis* strain** | **Genomic fragment (sequence range)** |  | **Similarity** | **Change in AT content** |  | **Transitions** | | **Transversions** | | **Total point mutations** | |  | **Transcriptomic fragment** | **Similarity** | **Query coverage** | **Translated protein sequence** |
|  |  |  |  |  |  | **C-to-T**  **&**  **G-to-A** | **T-to-C**  **&**  **A-to-G** | **C-to-A**  **&**  **G-to-T** | **A-to-C**  **&**  **T-to-G** | **G or C**  **to**  **A or T** | **A or T**  **to**  **G or C** |  |  |  |  |  |
| **1229** | **LKHE01000540 (19330→20932)** |  | **99.9%** | **No ∆ (38.7%)** |  |  |  |  |  |  |  |  | [**GCQL01006885**](https://www.ncbi.nlm.nih.gov/nuccore/GCQL01006885) **(201←504)** [**GCQL01011125**](https://www.ncbi.nlm.nih.gov/nuccore/GCQL01011125) **(1→411)** [**GCQL01011658**](https://www.ncbi.nlm.nih.gov/nuccore/GCQL01011658) **(1152→1882) with** **a 61-nt deletion** | **100% 99.8% 92.1%** | **18% 25% 49%** | [**EQL02970**](https://www.ncbi.nlm.nih.gov/protein/EQL02970.1?report=genbank&log$=protalign&blast_rank=2&RID=RF8PWZME016) **(28→110)** [**EQL02970**](https://www.ncbi.nlm.nih.gov/protein/EQL02970.1?report=genbank&log$=protalign&blast_rank=2&RID=RF8PWZME016) **(140→275)** [**EQL02970**](https://www.ncbi.nlm.nih.gov/protein/EQL02970.1?report=genbank&log$=protalign&blast_rank=2&RID=RF8PWZME016) **(281→523)** |
|  | **LKHE01001740 (54230←54311; 54418←55104)** |  | **67.9%** | **↓ 38.0% to 33.0%** |  | **68** | **68** | **26** | **40** | **94** | **108** |  | **[GCQL01006426](https://www.ncbi.nlm.nih.gov/nuccore/GCQL01006426" \t "lnkRPS6PD1W016" \o "Show report for GCQL01006426.1) (997→1188)** [**GCQL01020269**](https://www.ncbi.nlm.nih.gov/nuccore/GCQL01020269) **(3→392)** **[GCQL01008668](https://www.ncbi.nlm.nih.gov/nuccore/GCQL01008668" \t "lnkPA8BS5GY016" \o "Show report for GCQL01008668.1) (979→1256)** | **100% 99.7% 98.2%** | **21% 45% 31%** | **KAF4504658 (45→70) KAF4504658 (57→184) KAF4504658 (208→297)** |
|  | **LKHE01002381 (20238←21185)** |  | **64.2%** | **↑ 38.0% to 42.0%** |  | **74** | **46** | **48** | **43** | **122** | **89** |  | [**GCQL01012138**](https://www.ncbi.nlm.nih.gov/nuccore/GCQL01012138) **(368→1264) with** **a 51-nt intron** | **94.6%** | **100%** | [**EQL02706**](https://www.ncbi.nlm.nih.gov/protein/EQL02706.1?report=genbank&log$=protalign&blast_rank=1&RID=REZZHMC2013) **(56→354)** |
| **CC1406-203** | [**NGJJ01001580**](https://www.ncbi.nlm.nih.gov/nuccore/NGJJ01001580) **(70191→71142)** |  | **100%** | **No ∆ (38.7%)** |  |  |  |  |  |  |  |  | [**GCQL01006885**](https://www.ncbi.nlm.nih.gov/nuccore/GCQL01006885) **(228←504)** [**GCQL01011125**](https://www.ncbi.nlm.nih.gov/nuccore/GCQL01011125) **(1→411)** [**GCQL01011658**](https://www.ncbi.nlm.nih.gov/nuccore/GCQL01011658) **(1718→1882)** | **99.6% 99.8% 99.4%** | **17% 43% 29%** | [**EQL02970**](https://www.ncbi.nlm.nih.gov/protein/EQL02970.1?report=genbank&log$=protalign&blast_rank=2&RID=RF8PWZME016) **(37→110)** [**EQL02970**](https://www.ncbi.nlm.nih.gov/protein/EQL02970.1?report=genbank&log$=protalign&blast_rank=2&RID=RF8PWZME016) **(140→275)** [**EQL02970**](https://www.ncbi.nlm.nih.gov/protein/EQL02970.1?report=genbank&log$=protalign&blast_rank=2&RID=RF8PWZME016) **(281→335)** |
|  | **NGJJ01000083 (697372→698058; 698165→698246)** |  | **67.9%** | **↓ 38.0% to 33.0%** |  | **85** | **110** | **40** | **71** | **125** | **181** |  | [**GCQL01006426 (997→1188)**](https://www.ncbi.nlm.nih.gov/nuccore/GCQL01006426) [**GCQL01020269**](https://www.ncbi.nlm.nih.gov/nuccore/GCQL01020269) **(3→392)** [**GCQL01008668**](https://www.ncbi.nlm.nih.gov/nuccore/GCQL01008668) **(979→1256)** | **100% 99.7% 98.2%** | **21% 45% 31%** | **KAF4504658 (45→70) KAF4504658 (57→184) KAF4504658 (208→297)** |
|  | **NGJJ01000732 (250601→251549)** |  | **64.2%** | **↑ 38.0% to 41.9%** |  | **74** | **46** | **48** | **43** | **122** | **89** |  | [**GCQL01012138**](https://www.ncbi.nlm.nih.gov/nuccore/GCQL01012138) **(368→1264) with** **a 51-nt intron** | **94.5%** | **100%** | [**EQL02706**](https://www.ncbi.nlm.nih.gov/protein/EQL02706.1?report=genbank&log$=protalign&blast_rank=1&RID=REZZHMC2013) **(56→354)** |
| **Co18** | **ANOV01000528 (2857←4459)** |  | **99.8%** | **Nearly no ∆ (↑ 38.7% to 38.9%)** |  | **0** | **0** | **1** | **0** | **1** | **0** |  | [**GCQL01006885**](https://www.ncbi.nlm.nih.gov/nuccore/GCQL01006885) **(201←504)** [**GCQL01011125**](https://www.ncbi.nlm.nih.gov/nuccore/GCQL01011125) **(1→411)** [**GCQL01011658**](https://www.ncbi.nlm.nih.gov/nuccore/GCQL01011658) **(1152→1882) with** **a 61-nt deletion** | **99.7% 99.8% 92.0%** | **18% 25% 49%** | [**EQL02970**](https://www.ncbi.nlm.nih.gov/protein/EQL02970.1?report=genbank&log$=protalign&blast_rank=2&RID=RF8PWZME016) **(28→110)** [**EQL02970**](https://www.ncbi.nlm.nih.gov/protein/EQL02970.1?report=genbank&log$=protalign&blast_rank=2&RID=RF8PWZME016) **(140→275)** [**EQL02970**](https://www.ncbi.nlm.nih.gov/protein/EQL02970.1?report=genbank&log$=protalign&blast_rank=2&RID=RF8PWZME016) **(281→523)** |
|  | **ANOV01000033 (136→549; 656→737)** |  | **70.4%** | **↓ 35.7% to 32.7%** |  | **16** | **27** | **14** | **15** | **30** | **44** |  | [**GCQL01006426 (997→1188)**](https://www.ncbi.nlm.nih.gov/nuccore/GCQL01006426) [**GCQL01020269**](https://www.ncbi.nlm.nih.gov/nuccore/GCQL01020269) **(3→392)** | **100% 99.7%** | **21% 65%** | **KAF4504658 (45→70) KAF4504658 (57→184)** |
|  | **ANOV01000652 (15299→16246)** |  | **64.2%** | **↑ 38.0% to 41.9%** |  | **74** | **46** | **48** | **43** | **122** | **89** |  | [**GCQL01012138**](https://www.ncbi.nlm.nih.gov/nuccore/GCQL01012138) **(368→1264) with** **a 51-nt intron** | **94.6%** | **100%** | [**EQL02706**](https://www.ncbi.nlm.nih.gov/protein/EQL02706.1?report=genbank&log$=protalign&blast_rank=1&RID=REZZHMC2013) **(56→354)** |
| **IOZ07** | **JAAVMX010000002 (238583←240185)** |  | **100%** | **No ∆ (38.7%)** |  |  |  |  |  |  |  |  | [**GCQL01006885**](https://www.ncbi.nlm.nih.gov/nuccore/GCQL01006885) **(201←504)** [**GCQL01011125**](https://www.ncbi.nlm.nih.gov/nuccore/GCQL01011125) **(1→411)** [**GCQL01011658**](https://www.ncbi.nlm.nih.gov/nuccore/GCQL01011658) **(1152→1882) with** **a 61-nt deletion** | **99.7% 99.8% 92.0%** | **18% 25% 49%** | [**EQL02970**](https://www.ncbi.nlm.nih.gov/protein/EQL02970.1?report=genbank&log$=protalign&blast_rank=2&RID=RF8PWZME016) **(28→110)** [**EQL02970**](https://www.ncbi.nlm.nih.gov/protein/EQL02970.1?report=genbank&log$=protalign&blast_rank=2&RID=RF8PWZME016) **(140→275)** [**EQL02970**](https://www.ncbi.nlm.nih.gov/protein/EQL02970.1?report=genbank&log$=protalign&blast_rank=2&RID=RF8PWZME016) **(281→523)** |
|  | **JAAVMX010000009 (883241→883927; 884034→884115)** |  | **67.9%** | **↓ 38.0% to 33.0%** |  | **68** | **68** | **26** | **40** | **94** | **108** |  | [**GCQL01006426 (997→1188)**](https://www.ncbi.nlm.nih.gov/nuccore/GCQL01006426) [**GCQL01020269**](https://www.ncbi.nlm.nih.gov/nuccore/GCQL01020269) **(3→392)** [**GCQL01008668**](https://www.ncbi.nlm.nih.gov/nuccore/GCQL01008668) **(979→1256)** | **100% 99.7% 98.2%** | **21% 45% 31%** | **KAF4504658 (45→70) KAF4504658 (57→184) KAF4504658 (208→297)** |
|  | **JAAVMX010000003 (16799894←16800841)** |  | **64.2%** | **↑ 38.0% to 42.0%** |  | **74** | **46** | **48** | **43** | **122** | **89** |  | [**GCQL01012138**](https://www.ncbi.nlm.nih.gov/nuccore/GCQL01012138) **(368→1264) with** **a 51-nt intron** | **94.6%** | **100%** | [**EQL02706**](https://www.ncbi.nlm.nih.gov/protein/EQL02706.1?report=genbank&log$=protalign&blast_rank=1&RID=REZZHMC2013) **(56→354)** |
| **ZJB12195** | **LWBQ01000052 (168259←169259)** |  | **100%** | **No ∆ (40.2%)** |  |  |  |  |  |  |  |  | [**GCQL01011125**](https://www.ncbi.nlm.nih.gov/nuccore/GCQL01011125) **(1←196)** [**GCQL01011658**](https://www.ncbi.nlm.nih.gov/nuccore/GCQL01011658) **(1149→1882) with** **a 61-nt deletion** | **100% 92.2%** | **19% 79%** | [**EQL02970**](https://www.ncbi.nlm.nih.gov/protein/EQL02970.1?report=genbank&log$=protalign&blast_rank=2&RID=RF8PWZME016) **(212→275)** [**EQL02970**](https://www.ncbi.nlm.nih.gov/protein/EQL02970.1?report=genbank&log$=protalign&blast_rank=2&RID=RF8PWZME016) **(281→524)** |
|  | **LWBQ01000002 (1407095←1407176; 1407283←1407696)** |  | **70.4%** | **↓ 35.4% to 33.0%** |  | **16** | **27** | **14** | **15** | **30** | **42** |  | [**GCQL01006426 (997→1188)**](https://www.ncbi.nlm.nih.gov/nuccore/GCQL01006426) [**GCQL01020269**](https://www.ncbi.nlm.nih.gov/nuccore/GCQL01020269) **(3→392)** | **100% 99.7%** | **21% 65%** | **KAF4504658 (45→70) KAF4504658 (57→184)** |
|  | **LWBQ01000004 (210989←211936)** |  | **64.2%** | **↑ 38.0% to 42.0%** |  | **74** | **46** | **48** | **43** | **122** | **89** |  | **[GCQL01012138](https://www.ncbi.nlm.nih.gov/nuccore/GCQL01012138" \t "lnkPA8U1KZ3016" \o "Show report for GCQL01012138.1) (368→1264) with** **a 51-nt intron** | **94.6%** | **100%** | [**EQL02706**](https://www.ncbi.nlm.nih.gov/protein/EQL02706.1?report=genbank&log$=protalign&blast_rank=1&RID=REZZHMC2013) **(56→354)** |

**Note: “****→” and “←” indicate the sequence directions of the sense and antisense chains, respectively. “↑” and “↓” indicate increases and decreases in AT content, respectively.**
